# Membrane curvature association of amphipathic helix 8 drives constitutive endocytosis of GPCRs

**DOI:** 10.1101/2024.03.27.587027

**Authors:** Jan Hendrik Schmidt, Rasmus Herlo, Joscha Rombach, Andreas Haahr Larsen, Mikkel Stoklund, Mathias Perslev, Tommas Theiss Ehlers Nielsen, Keenie Ayla Andersen, Carmen Klein Herenbrink, Matthew D. Lycas, Aske Lykke Ejdrup, Nikolaj Riis Christensen, Jan P. Christensen, Mootaz Salman, Freja Herborg, Ulrik Gether, Alexander Sebastian Hauser, Patricia Bassereau, David Perrais, Kenneth Lindegaard Madsen

**Author notes:** Correspondence and requests for materials should be addressed to Kenneth Lindegaard Madsen.

## Abstract

Cellular signaling relies on the activity of transmembrane receptors and their presentation on the cellular surface. Their continuous insertion in the plasma membrane is balanced by constitutive and activity dependent internalization, which is orchestrated by adaptor proteins recognizing semi-specific motifs within the receptors’ intracellular regions. Here we describe a complementary and evolutionary conserved and refined trafficking mechanism for G-protein coupled receptors (GPCR). This mechanism relies on the insertion of their amphipathic helix 8 into the inner leaflet of lipid membranes, orthogonal to the transmembrane helices. These amphipathic helices dictate subcellular localization of the receptors and autonomously drive their endocytosis by cooperative assembly and association with areas of high membrane curvature. The strength of helix 8 membrane insertion propensity quantitatively predicts the rate of constitutive internalization of GPCRs. This discovery advances our understanding of membrane protein trafficking and highlights a new principle of receptor-lipid interactions that may have broader implications for cellular signaling and therapeutic targeting.

**One-Sentence Summary:** Receptor proteins navigate cellular membranes by interacting with their curvature using an evolutionary conserved mechanism that relies on amphipathic helices and complements direct coupling to the endocytic protein machinery.

## INTRODUCTION

Cellular trafficking and sorting processes direct transmembrane proteins (TMPs) to their proper destination within the cell. Consequently, these processes are of fundamental importance for cellular homeostasis, metabolism, and communication. Malfunction results in diseases including cancer, metabolic syndromes, neuropathies, neuro-degenerative disease and psychiatric disorders ^1^.

The transport of TMPs between different cellular compartments is mediated through the action of vesicular membrane carriers. Specificity in trafficking itineraries is achieved by selective TMP cargo sorting into these carriers, eventually giving rise to differential localizations of TMPs within the cell. This sorting relies partly on semi-specific protein-protein interactions between compartment specific adaptor proteins (e.g. AP1-5 and GGAs) and short linear motifs within the cargo TMPs (e.g. FXNPXY, tyrosine- and dileucine-based signals) ^2,3^. However, these signals are highly ambiguous and offers little predictive power, leaving critical gaps in our understanding of how TMP localization is precisely regulated within cells.

The initial budding and formation of vesicular carriers involves generation of high membrane curvature which serves to recruit specific scaffolding and regulatory cytosolic proteins ^4,5^. This mechanism, termed membrane curvature sensing (MCS), is frequently mediated by amphipathic helices (AHs) that insert partially into exposed lipid packing defects in the convex leaflet of curved membranes ^6,7^. We hypothesized that AHs in the intracellular termini of TMPs, positioned at the membrane-cytoplasm interface, serve to partition TMPs into areas of high curvature during formation of vesicular carriers thereby serving as a generic mechanism driving differential forward trafficking of TMPs in the endolysosomal system.

Using G protein-coupled receptors (GPCRs) as example, our data demonstrates that AHs direct constitutive trafficking and redistribution of TMPs with rates quantitatively predicted by their membrane insertion propensity (MIP). We demonstrate that this new trafficking mechanism of TMPs, which relies on direct association with highly curved membranes in budding membrane carriers, is fundamentally distinct in terms of kinetics and molecular association from conventional adaptor-based TMP trafficking processes, such as agonist-induced internalization. Moreover, we present data suggesting that MIP of GPCR AHs is conserved across GPCR families and animal species and has been refined during the course of evolution. We refer to this novel mechanism as Trafficking of Transmembrane proteins by Amphipathic Motifs (ToTAM) and propose it as a complementary means of cellular sorting and localization to adaptor-based sorting of TMPs.

## RESULTS

### Amphipathic helices drive constitutive internalization of transmembrane proteins

G protein-coupled receptors (GPCRs) constitute the largest family of TMPs in the human genome and include receptors for diverse signaling molecules, such as small molecule neurotransmitters, peptides, glycoproteins, lipids, nucleotides, and ions ^8^. They control vital physiological functions and represent targets for roughly a third of FDA approved drugs ^9^. GPCRs consist of 7 transmembrane α-helices and a commonly amphipathic 8th helix (H8) that can insert into the cytosolic leaflet of the membrane orthogonal to the transmembrane helices (Figure 1A) ^10–13^.

**Figure 1.**
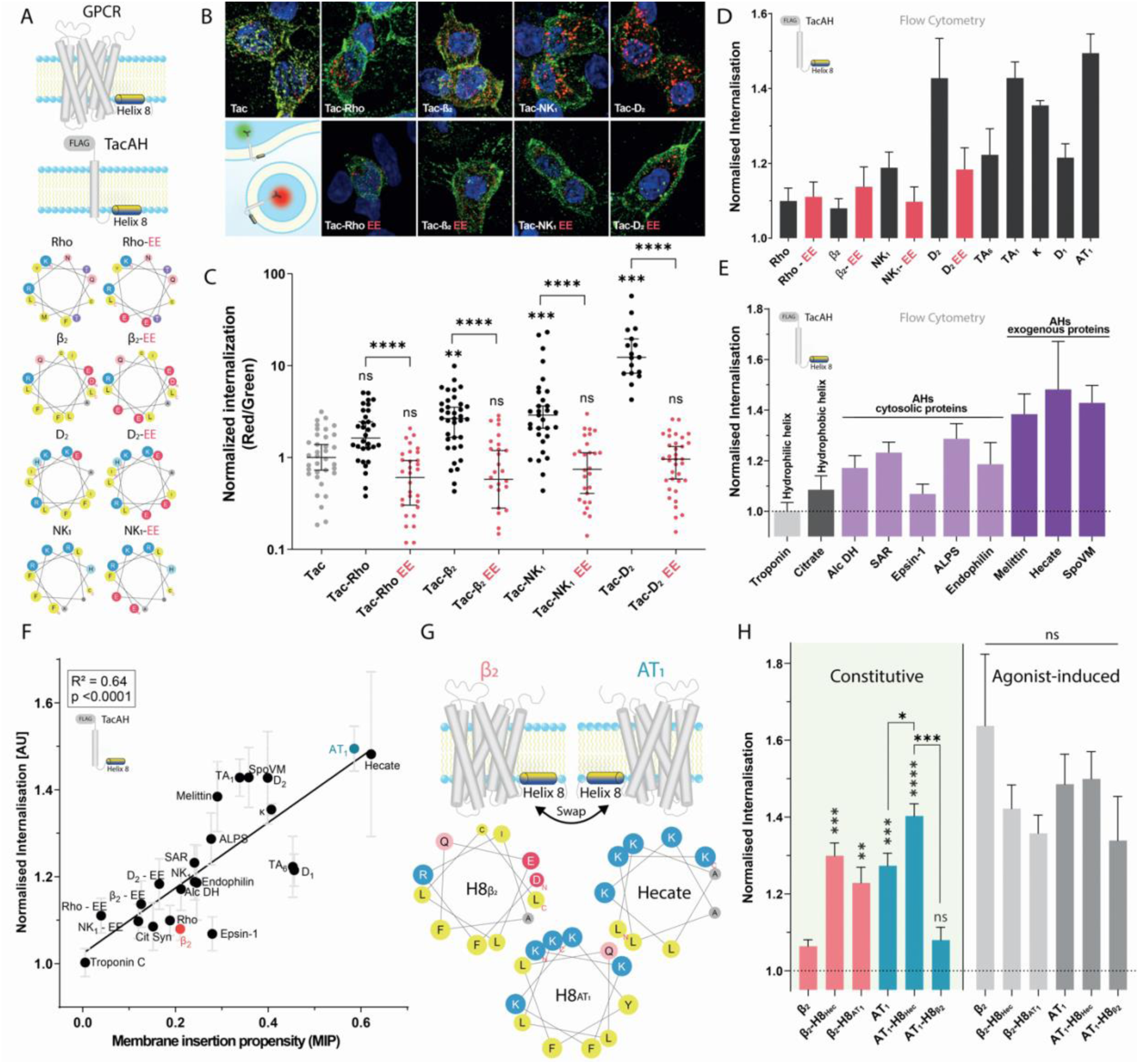
Amphipathic helices in transmembrane proteins autonomously drive their endocytosis. **(A)** Illustration of GPCR H8 sequences transferred to the single transmembrane protein Tac for determination of their autonomous endocytic drive alongside mutational disruption of amphipathicity by introduction of negative charges in the hydrophobic face of H8. **(B)** Representative images of HEK293 cells transfected with Tac-H8 constructs and subjected to antibody feeding (illustrated in insert, lower left). **(C)** Quantification of internalization (Red/Green fluorescence ratio), relative to Tac, from confocal images (every dot represents a cell) N=3. **(D)** Internalization of Tac-H8 constructs assessed by flow cytometry and normalized to Tac. Mean ± SEM, N=3. **(E)** Internalization of Tac with AHs from cytosolic proteins or exogenous AHs assessed by flow cytometry. Mean ± SEM, N=3. **(F)** Linear correlation (R^2^ = 0.64 p <0.0001) between Membrane Insertion Propensity (MIP) of Tac-AH constructs and their respective endocytic rate (relative to Tac). Mean ± SEM, N=3. **(G)** Illustration of chimeric H8 GPCRs with swapped part highlighted in helical wheel representations using HeliQuest ^43^. **(H)** Constitutive and agonist-induced (10 µM agonists; ISO = Isoproterenol and ANGII = angiotensin II) internalization rates of chimeric FLAG-tagged GPCR constructs with swapped H8 sequences relative to Tac. Mean ± SEM, N=3.

Interactions between H8 and cytosolic adaptor proteins are known to mediate various trafficking processes, particularly agonist-induced endocytosis, but also constitutive internalization ^14^.

To test our hypothesis, that AHs in the intracellular termini of TMPs can drive their forward trafficking in the endolysosomal system, we tested the autonomous function of H8 in endocytosis. We fused peptide sequences harboring H8 (17-18 residues) from a number of well-characterized class A GPCRs (rhodopsin (Rho), β2 adrenergic receptor (β_2_), substance P receptor (NK_1_), and dopamine D2 receptor (D_2_)) to the cytosolic C-terminus of the internalization inert transmembrane interleukin-2 receptor a-subunit (IL-2a), hereafter referred to as Tac (Figure 1A) ^15,16^. The addition of AHs increased internalization of Tac as determined by quantitative confocal microscopy of antibody feeding experiments in HEK293 cells (up to 10-fold for Tac-D2) (Figure 1B-C). Disrupting the hydrophobic wedge of the amphipathic helices by introducing negative charges (Figure 1A) abrogated this internalization. We recapitulated these findings in HeLa cells (Figure S1). Alanine substitution of cysteines, previously shown to modify trafficking of GPCRs through acylation ^17^, did not further affect internalization in this context (Figure S2).

After confirming the relative internalization rates with high-throughput quantitative flow cytometry ^18^ (Figure S3), we showed that five additional H8s with distinct amphipathic nature, i.e., with high hydrophobic moment, also increased Tac internalization (Figure 1D). These were H8 from trace amine receptor 1 and 6 (TA_1_ and TA_6_), kappa opioid receptor (κ), dopamine D1 receptor (D_1_) and angiotensin 2 type 1 receptor (AT_1_) (biochemical parameters outlined in Figure S4). To consolidate and broaden these findings, we also tested non-GPCR helices. Fusion of a hydrophilic helix from troponin^19^ or the hydrophobic helix from citrate^20^ did not increase Tac internalization, whereas AHs of several membrane-associated proteins involved in membrane trafficking, including Amphipathic Lipid Packing Sensor (ALPS), Sar1 and Endophilin A1, but not Epsin-1^21,22^ facilitated internalization (Figure 1E). Similarly, exogenous AHs, including those from the bee venom Melittin and the bacterial protein SpoVM as well as the artificial AH Hecate, strongly facilitated internalization of Tac. Together these findings demonstrate that AHs, diverse in origin and devoid of known adaptor motifs, autonomously increase endocytosis of a transmembrane protein. We refer to this mechanism as Trafficking of Transmembrane proteins by Amphipathic Motifs (ToTAM).

### The membrane insertion propensity of AHs quantitatively predicts their effect on TMP internalization

To quantify the propensity of individual AHs to associate with positive membrane curvature, we calculated the hydrophobic moment of AHs and adjusted it by their intrinsic helical propensity (see methods)^23^. The resulting value, membrane insertion propensity (MIP), correlated with empirical measures of membrane association strength as determined by liposome binding to peptides on a SPOT array (Figure S5). Furthermore, plotting internalization of the AH-fused Tac- constructs as a function of MIP yielded a remarkably strong correlation irrespective of the origin of the AHs (Figure 1F), demonstrating that AHs drive internalization of TMPs in a predictive manner. In comparison, net charge and hydrophobicity demonstrated much lower predictive power on their own (Figure S6). In absolute terms, AHs increased internalization from a basal level of 20% of Tac over a 30-minute period to a complete (100%) turnover of Tac-AHs for the strongest AHs (Figure S7).

### The amphipathic nature of H8 drives constitutive endocytosis of full length GPCRs

To address the endocytic function of H8 in context of full-length receptors, we focused on β_2_ and AT_1_, which contain H8s with pronounced differences in their autonomous endocytic drive (Figure 1F, red and blue, respectively). We made chimeric constructs by interchanging their H8s (Figure 1G) and assessed their internalization. In concordance with the autonomous drive of their H8s, AT_1_ displayed significantly higher constitutive internalization than β_2_, and this pattern was fully reversed by swapping their helices (Figure 1H and S8). Furthermore, the introduction of the synthetic, potent AH Hecate (Figure 1G) gave rise to an even higher internalization rate than that of AT_1_ H8, in context of both β_2_ and AT_1_. In contrast, internalization in presence of agonist was robust for all constructs irrespective of AH (Figure 1H and S8). Further, while constitutive internalization scaled linearly with expression, agonist induced internalization for the H8 containing constructs deviated from linearity consistent with saturation of adaptor-based internalization machinery (Figure S8)^24^. This suggests that MIP of H8 drives constitutive, but not agonist-induced, internalization of GPCRs in a mechanistically novel fashion.

### AHs in TMPs regulate their cellular distribution at steady state

To assess how AHs might affect the steady state distribution of TMPs we adapted a proximity biotinylation approach. We chose a series of five Tac-AH constructs, spanning from low to high MIP values and fused the small (26.4kDa) biotin ligase BioID2^25^ to their C-terminal end (Figure 2A). Western blot analysis confirmed expression of proteins of the expected size (62kDa, lower band unglycosylated) and further suggested a reduced glycosylation (>75kDA, Upper band) with increasing MIP (Figure 2B, top). The BioID2-fusion constructs recapitulated ToTAM driven internalization as assessed by quantitative confocal microscopy (Figure 2B, bottom and S9A-B).

**Figure 2.**
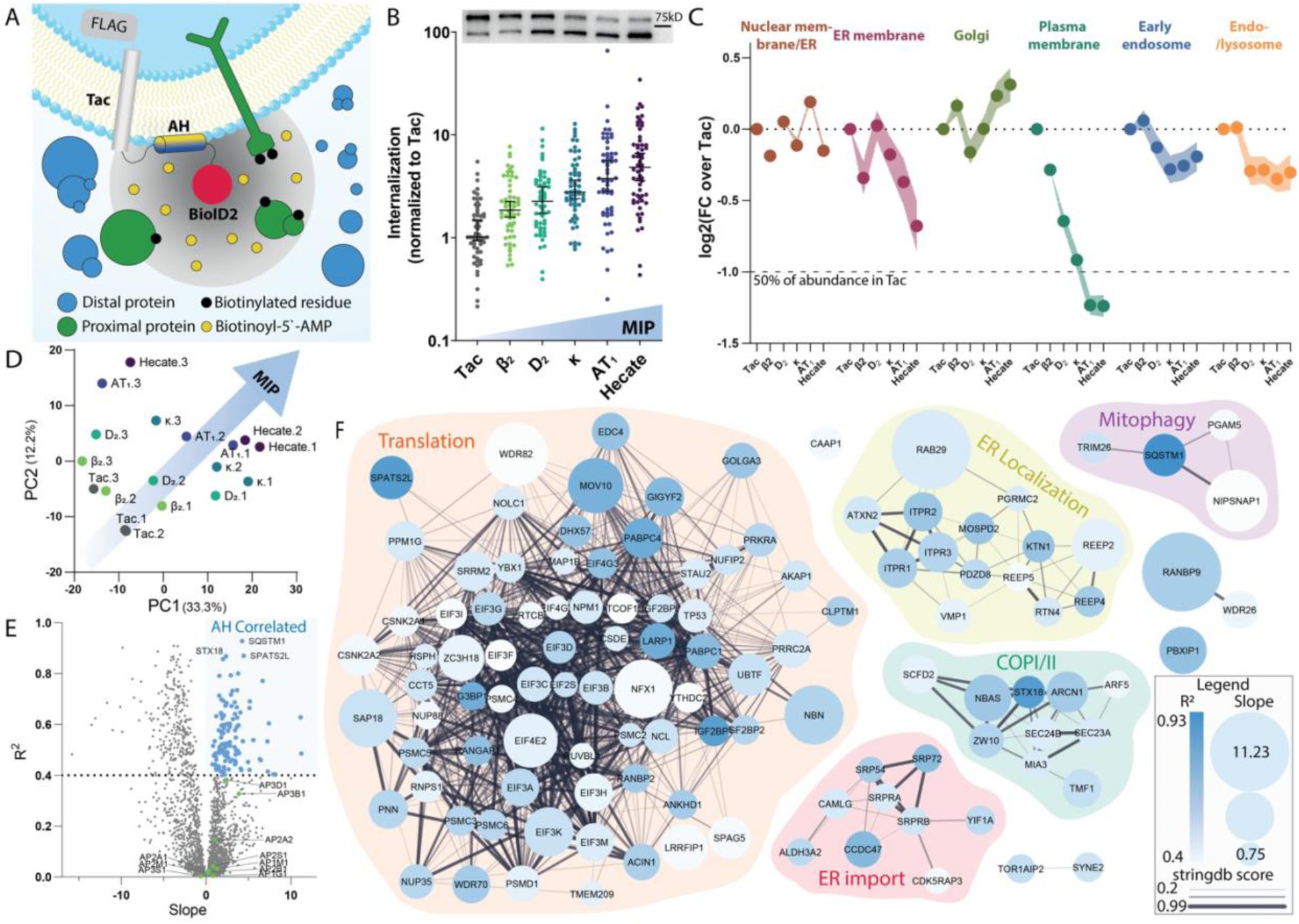
Increasing MIP drives association with the biosynthetic pathway and away from the plasma membrane. **(A)** Schematic of proximity biotinylation assay with BioID2. **(B)** (Upper) Western blot (M1 ab) of Flag-Tac AH-BioID2 constructs transfected in HEK293 cells showing mature proteins (high MW band) and immature protein (low MW band). (Lower) Quantification of internalization of Flag-Tac AH BioID2 constructs from confocal images (every dot represents a cell, compiled from three independent experiments). **(C)** Differences in label-free quantification (LFQ) values for indicated constructs versus basal levels (pcDNA) normalized to Tac compiled for all identified proteins belonging to endolysosomal compartments of MMF grouping in ^26^. **(D)** Principal component analysis biplot showing spread of MS samples along principal component 1 and 2 indicating a MIP driven separation. **(E)** Volcano plot highlighting slope and r^2^ values of linear correlations for each identified protein with MIP of the Tac-AH-BioID2 constructs. Protein correlating with MIP (slope > 0 and r^2^ > 0.4) highlighted in blue. Identified adaptor protein (AP) complex subunits (indicated in green). **(F)** STRING protein network analysis of the proteins identified as MIP correlated in (E) in functional groups based on database annotations.

Following a 24-hour incubation with excess biotin, we performed affinity purification of the biotinylated proteins to assess their differential association with the AH containing BioID2-fusion constructs by quantitative label-free LC-MS^2^ ^25^. We identified a total of 2569 unique proteins and grouped them by sub-cellular localization according to a negative matrix factorization (NMF) approach previously applied to proximity biotinylation experiments ^26^ and evaluated relative enrichment of proteins found in TacAH-BioID2 transfected cells versus mock (pcDNA). Proteins belonging to the endolysosomal pathway were among the most enriched (Figure S9C), validating the proximity labelling strategy. Among the subcellular compartments belonging to the endosomal system only “plasma membrane” exhibited significantly differential enrichment across the BioID2-fusion proteins revealing a strong negative association with increasing MIP values consistent with reduced maturation and increased internalization (Figure 2C). Weaker, non-significant trends were observed for the remaining endolysosomal compartments.

Upon analysis of the total variance of the data by principal component analysis, we observed a MIP-dependent distinction of protein labelling intensities by TacAH-BioID2 contructs (Figure 2D). We next asked whether MIP—value was associated with any individual proteins. Linear regression analysis of all identified proteins highlighted a subset of proteins which showed strong positive correlation (R^2^ >0.4) with MIP (indicated in light blue). Of note, none of the identified adaptor protein (AP) complex subunits (indicated in green) fulfilled these criteria (Figure 2E). Protein network analysis of the strongly MIP correlated subset revealed several tightly connected clusters annotated as related to translation, ER localization, ER import, COPI/II complex and mitophagy (Figure 2F). Interestingly, plotting all identified core proteins of the COPI and COPII complexes revealed an overall increased abundance with MIP (Figure S9D+E). Together these findings suggest a MIP dependent steady state enrichment of TMP within the early biosynthetic pathway as well as a depletion at the plasma membrane.

### ToTAM is associated with clathrin

Since we found no significant associations between MIP and any classical or putative endocytic adapter proteins, we asked more qualitatively how the Tac-BioID2 constructs differentially labelled established components of the endocytic machinery similar to the analysis for the COPI and COPII complexes. The MIP of the expressed Tac-AHs did not significantly affect the enrichment of actin or dynamin, (Figure 3A) consistent with a low impact of treatment with the actin polymerization inhibitor Latrunculin A, EIPA and Dyngo-4A on internalization of Tac-AHs (Figure S10A-C). In comparison, ATP-depletion resulted in a relative reduction of internalization for all constructs (Figure S10D-F), indicating involvement of energy-dependent processes in ToTAM.

**Figure 3.**
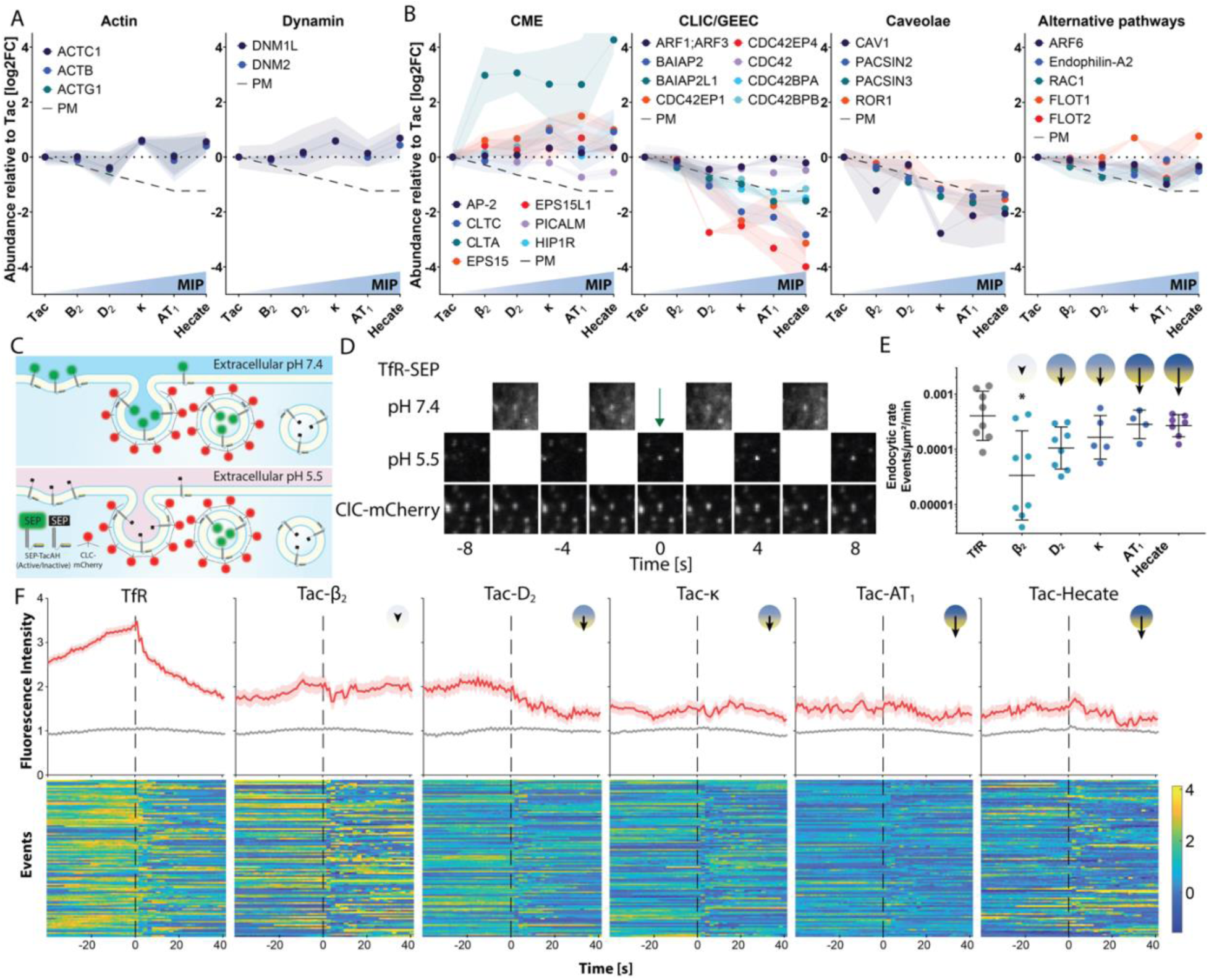
ToTAM associates with CME machinery in a non-canonical manner. **(A+B)** Absolute difference in LFQ values (normalized to Tac) of identified actin and dynamin subunits (A) and selected identified proteins associated with CME, CLIG/GEEC, Caveolae and other endocytic pathways (B). **(C)** Schematic of (ppH-TIRFM) protocol enabling definition of time of scission for individual endocytic events. **(D)** Images of representative endocytic event from ppHTIRFM experiments of Super Ecliptic pHluorin (SEP) tagged transferrin receptor (TfR) together with clathrin light chain (CLC)-mCherry. **(E)** Endocytic event rate for the 5 different SEP-TacAH constructs compared to TfR-SEP. Statistical analysis using Kruskal-Wallis multiple comparisons test. **(F)** (Upper) Average of individually normalized CLC-mCherry traces for TfR-SEP and 5 SEP-TacAH constructs. (Lower) Heatmaps of normalized individual CLC-mCherry traces pre- and post-scission. N = 5-10 cells (846-2797 events) per condition.

Next, we addressed association of the Tac-BioID2 constructs with components of specific endocytic pathways. Key proteins of clathrin-mediated endocytosis (CME)^27^, in particular clathrin and EPS15, were enriched for all Tac-AHs (Figure 3B). On the other hand, identified components of the CLIC/GEEC pathway, and caveolae^28,29^, were consistently less enriched with increasing MIP of the Tac-AHs. Finally, enrichment of proteins involved in alternative endocytic pathways, including Arf6, Rac1, Endophilin A2, and Flotillins (FLOT1 and FLOT2 ^30^), was relatively independent of MIP. Taking into account the reduced surface expression of Tac-BioID2 constructs as a function of MIP (dashed line), we interpret that the AHs likely do not actively drive sorting away from CLIC/GEEC or caveolae, but that AHs potentially sort TMPs toward alternative endocytic pathways involving Arf6, Rac1, Endophilin A2, and Flotillins, in addition to CME.

Grouping all identified CME associated proteins according to temporally defined modules ^31^, we observed no marked increase in association with nucleation and assembly, neck constriction or scission and movement modules, while proteins involved in uncoating as well as endosome fusion were enriched for all AHs (Figure S11A). Notably, proteins of the nucleation and formation module that showed increased abundance, including clathrin and EPS15, represented a group that has been shown to be continuously associated with CME events until uncoating ^27^. EPS15 is known to recruit ubiquitinated cargo to endocytic sites ^32^. In accordance, we detected minor increases in ubiquitin binding proteins of the ESCRT machinery and E3 ubiquitin ligases ^33^ (Figure S11B+C), as well as the ubiquitin-receptor SQSTM1 (p62) as the top hit among MIP correlated associations (Figure 2E+F). This suggests an interaction with the ubiquitin machinery, despite no evident ubiquitination of the Tac-AH constructs (Figure 2B). Taken together, ToTAM associates with CME and likely with components of alternative pathways, while seemingly avoiding GLIC/GEEC and caveolae in addition to the downstream ubiquitin machinery.

### ToTAM is temporally uncoupled from classical CME

To directly evaluate the trafficking dynamics of Tac-AH constructs and the CME structures they associate with, we turned to pulsed pH (ppH) TIRFM assay ^34^ using super ecliptic pHluorin (SEP) tagged constructs co-expressed with mCherry-tagged clathrin light chain (CLC) (Figure 3C-F and S12). Corroborating the previous findings, the number of endocytic events increased by almost an order of magnitude with increased MIP of the AHs, approaching the rate of the transferrin receptor (TfR) for the strongest helices (H8 from AT_1_, and Hecate) (Figure 3E). Interestingly, exocytosis increased correspondingly, suggesting efficient intracellular sorting to recycling endosomes (Figure S13). CLC fluorescence intensity traces associated with TfR endocytic events (red line showing averaged traces) displayed a typical profile with gradual accumulation until point of scission followed by a rapid decrease after scission (Figure 3F). This distinct profile was gradually lost with increasing MIP of the Tac-AH constructs indicating that scission is temporally uncoupled from classical CME, although some spatial association with CLC was retained for all constructs (red line is above the gray line showing the average level of CLC away from endocytic events). Examination of individual normalized traces displayed a range of CLC recruitment profiles (Figure 3F (Lower) and Figure S14). Principal component analysis revealed a decreasing fraction of traces similar to the typical TfR-associated CLC profile for Tac-AHs with increasing MIP (Figure S15). Conversely, an increased fraction of events displayed accumulation of CLC immediate post scission. In summary, increased MIP directs ToTAM towards bulk association with CME core machinery at the expense of other endocytic machineries, while simultaneously decreasing the temporal synchronization and coordination with clathrin-dynamics.

### ToTAM involves accelerated endocytic maturation and scission

Next, we analyzed the recruitment dynamics of the SEP-Tac-AH constructs themselves prior to scission (Figure 4A). While SEP-TfR accumulated at a steady rate during vesicle formation ^35^, the AH-linked profiles showed fast and accelerating kinetics with increasing MIP of AHs. Fitting these profiles with a logistic equation indicated an increasingly cooperative clustering mechanism consistent with an auto-nucleating function underlying ToTAM (Figure 4B). To determine if this effect translates into a generalized increase in maturation rate of endocytic structures, we performed Tf-uptake in cells expressing the Tac-AH constructs. This demonstrated a strong and MIP dependent increase in transferrin uptake *in trans* (Figure 4C). Cargo load of individual endocytic events determined from the ppH-TIRFM experiments (pH 5.5 fluorescence at the time of scission) was decreased with increasing MIP of AHs (Figure 4D), whereas quenching rates as result of endosomal acidification increased with MIP of AHs (Figure 4E). Relating the cargo load to the acidification rates as a measure of endosome size (3D volume) further suggests an increased cargo density with increasing MIP of AHs (Figure S16). Taken together, we find that ToTAM is characterized by numerous, fast, small and cooperative endocytic events with the potential to accelerate overall trafficking rates.

**Figure 4.**
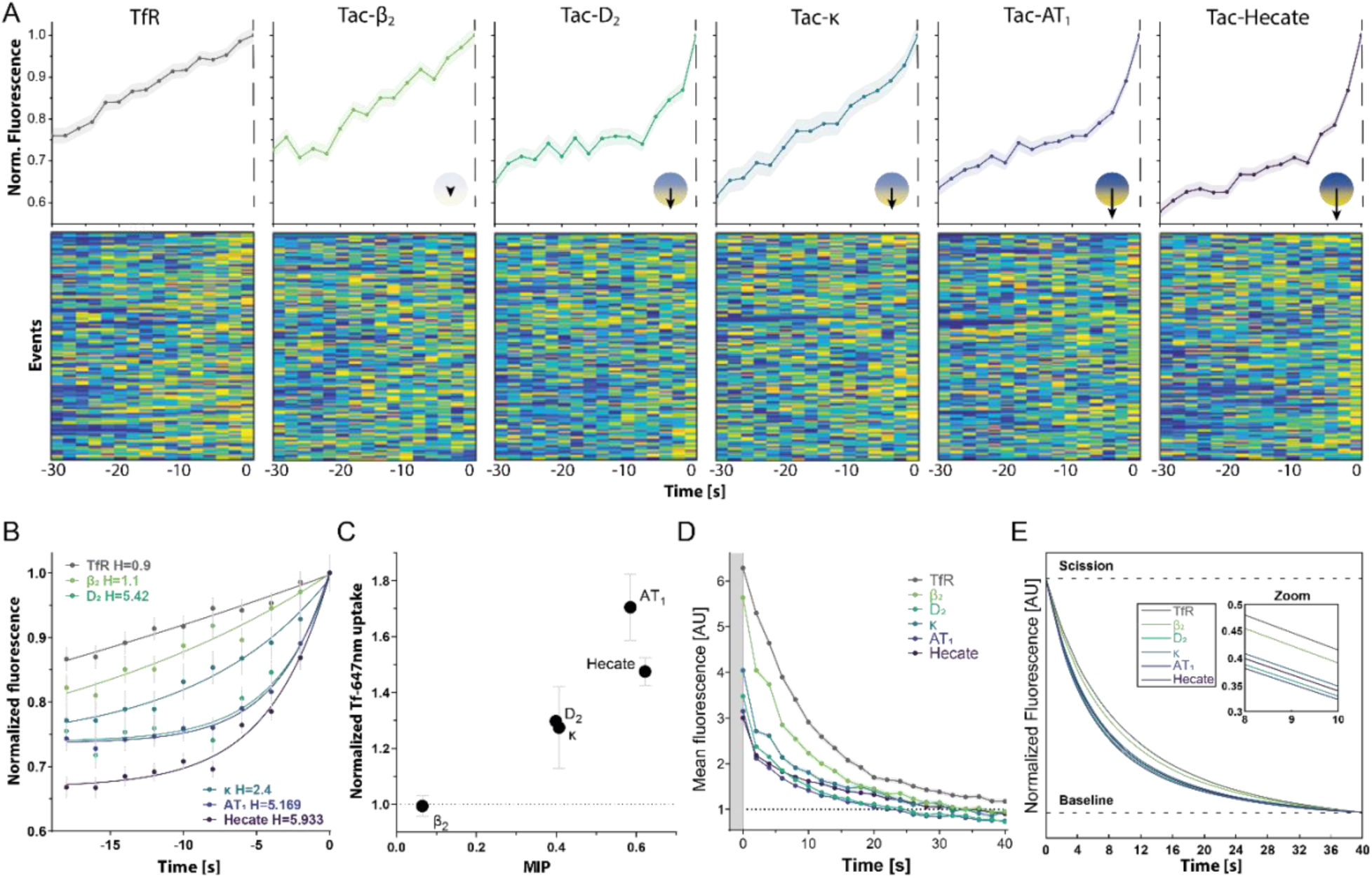
ToTAM involves accelerated endocytic maturation and scission. **(A)** (Upper) Average of individually normalized pH 7.4 TfR-SEP and SEP-TacAH fluorescence intensity traces building up to point of scission (Time = 0). (Lower) Heatmaps of normalized individual SEP-fluorescence profiles at pH 7.4 of TfR-SEP and SEP-TacAH constructs prior to scission. N = 5-10 cells per condition, n = 518-2367 total traces. **(B)** Non-linear regression fits of averaged normalized pHluorin traces up to point of scission. H is the Hill coefficient. **(C)** Effect of TacAH constructs on AF647-labelled transferrin endocytosis in trans, by flow cytometry. **(D)** Averaged fluorescent traces measured at pH 5.5 in cell populations expressing either of the six indicated SEP-TacAH-constructs. Absolute values shown post-scission (t=0). **(E)** Average traces from (D) are normalized to initial fluorescence observed at time of scission (t=0) and fitted to the “Acidification Kinetics Function” (see methods). Fits shown for all six SEP-TacAH-constructs with TfR eliciting slowest acidification kinetics, while AT_1_ elicits the fastest acidification kinetics.

### ToTAM associated with high membrane curvature during endocytosis

The temporal uncoupling with canonical endocytic machinery as well as the cooperative, high-density recruitment and the inductive properties of ToTAM collectively point to an underlying autonomous and recursive mechanism, consistent with concurrent membrane curvature sensing and induction, which is a common characteristic of AH association with lipid membranes. We therefore investigated the membrane curvature sensing properties of Hecate (the AH tested with highest MIP) in a buckled membrane system using molecular dynamics simulation. This simulation predicted a curvature sensitive membrane sorting (Figure 5A-B), akin to previous findings for ALPS ^36^. To experimentally confirm association with high membrane curvature we incubated Bodipy-Tr labelled giant unilamellar vesicles (GUVs) (red) with Oregon green 488 labelled Hecate peptide (green). Consistent with the cooperative clustering observed in the ppH- TIRF experiments, the Hecate peptide occasionally formed bulging microdomains on GUVs in the absence of manipulation indicative of membrane curvature induction (Figure 5C).

**Figure 5.**
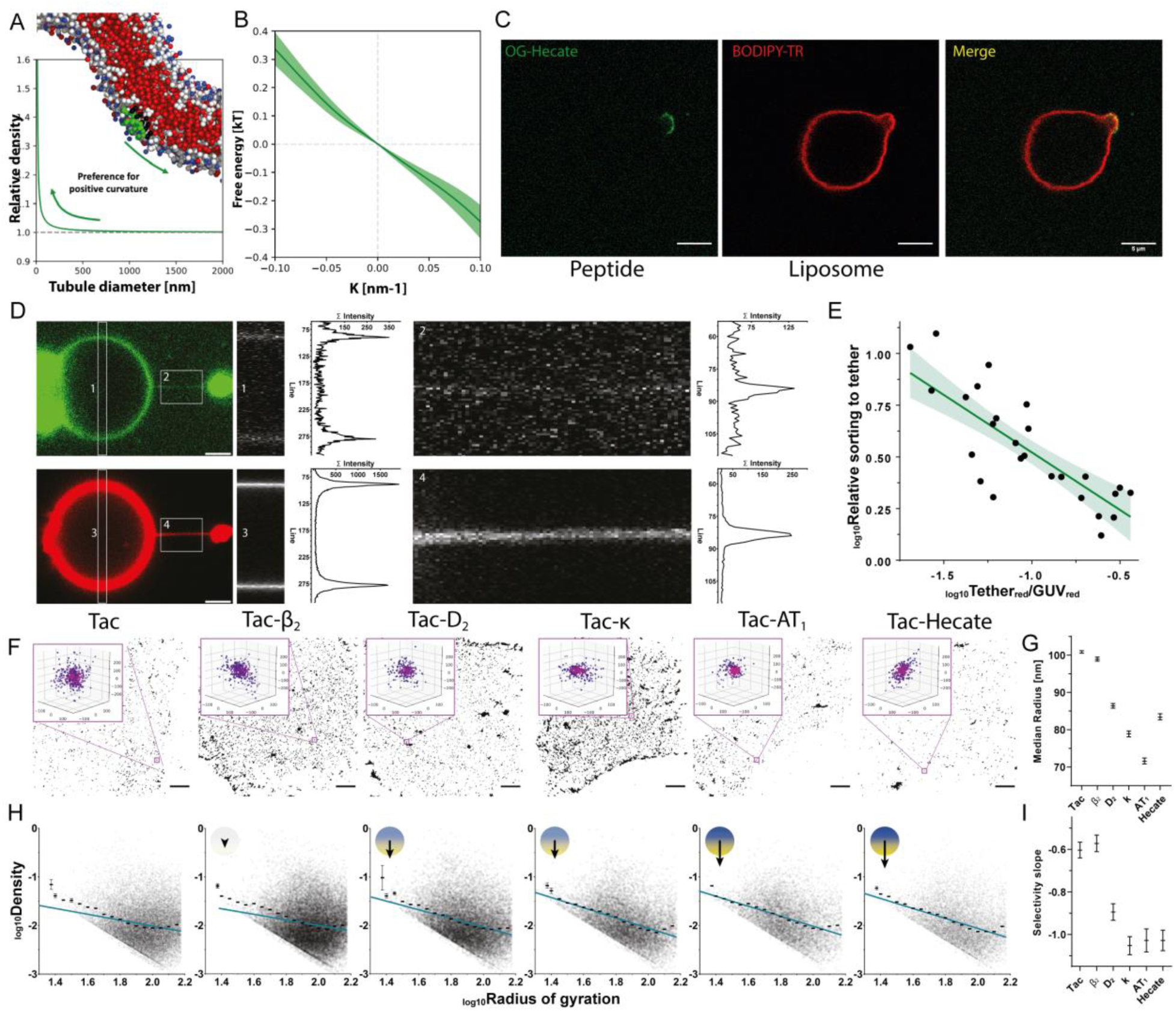
ToTAM involves association with high membrane curvature during endocytosis. **(A)** Relative protein density (Hecate peptide) as a function of tubule diameter determined in buckled membrane MD analysis indicating a strong preference for positive curvature. **(B)** Free energy from (A) as a function of membrane curvature. **(C)** Representative images of Oregon Green 488 labeled Hecate (green) bound to free floating BODIPY-TR labeled GUVs (red) revealing selective organization of peptides in deformed microdomain structures on the GUVs. Scale bars indicate 5 µm. **(D)** Tethers pulled from GUVs (red) pre-incubated with Oregon Green labelled Hecate peptide (green). Insert 1 and 3, zooms of GUV and summed intensity profile. Insert 2 and 4, zooms of tether and summed intensity profile. **(E)** Quantification of the relative sorting to the tether as a function of relative size of the tether (Tethered/GUVred). **(F)** Representative 3D-dSTORM images of FLAG-Tac and 5 FLAG-TacAH constructs after surface labelling with Alexa Fluor 647-conjugated M1 α-FLAG antibody, scale bar 5 µm. Inserts show representative single clusters color coded for localization density. **(G)** Average size of clusters for the individual constructs, median ± 95% CI, N=6. **(H)** Logarithmic relationship of cluster size and cargo density (FLAG-Tac-AH). Black bars show binned averages ± SEM and the blue line represents a linear fit of the data. Better adherence of the fit to the binned averages indicates curvature dependent sorting for Tac-AH constructs with high MIP. **(I)** Selectivity slope (i.e., curvature sensitivity) derived for linear fits with error on log-log plots of the data in (H) (see Figure S18).

Next, we pulled tethers from GUVs preincubated with OG488-Hecate to generate a high membrane curvature domain, and indeed observed a significant enrichment of the Hecate peptide on the tether (peptide on tether relative to GUV) (Figure 5D-E). A control peptide with similar charge but reduced helical propensity and MIP of the helix, did not bind to GUVs (Figure S17A+B). Given this *in vitro* evidence, we next tested whether the Tac-AH constructs displayed membrane curvature association at the plasma membrane of cells. To assess the size and curvature of the endocytic budding structures encompassing the cargo prior to scission, we antibody-labelled surface expressed Tac-AHs for 3D-dSTORM imaging and identified protein clusters using DBSCAN analysis (Figure 5F and S18). While the median radius of clusters was 100 nm for Tac, consistent with the size of clathrin-coated pits ^37^ cluster sizes were consistently smaller with increasing MIP of AHs (Figure 5G). We further extracted the number of localizations per cluster, enabling us to plot protein densities of individual budding structures as a function of size. In our previous studies, we have demonstrated and rationalized by modelling the membrane curvature sensing by amphipathic insertion into lipid membrane defects give rise to a log-log linear relationship ^6^. Indeed, the averaged cargo densities for the different size bins gradually approached a linear relationship (indicated by the dashed line) with increasing MIP of the AHs (Figure 5H). Moreover, the negative slope, which indicates the degree of membrane curvature sensing nominally increased as a function of the MIP of the AH (Figure 5I)^6^. In summary, we propose that ToTAM mechanistically serves both to localize cargo to endocytic pits as well as to facilitate the membrane curvature induction and possibly scission of these pits to drive endocytosis.

### Mapping of MIP in the C-termini of human GPCRs

To characterize the membrane association of C-termini across the full range of Human GPCRs, ^35^ we developed the liposome binding SPOT assay (Figure S5) into a large-scale chip array format using consecutive 16-mer peptides with a 2AA offset (Figure 6A-B). Reconstruction of binding profiles from individual peptide binding values revealed confined membrane binding regions of different strength as illustrated by the C-terminal regions of rhodopsin, β_2_ and AT_1_ (Figure 6C). These regions overlap with the H8 region as indicated by their high MIP-values. All receptor profiles are displayed in Figure S19. Averaged membrane binding profiles for each of the GPCR families confirmed this pattern for rhodopsin (disregarding olfactory receptors) as well as for secretin and frizzled families, albeit with membrane binding extending slightly further away from the TM region for the latter. Membrane binding was less pronounced for the remaining families and absent from the glutamate receptor family (Figure 6D). Moreover, these binding profiles are in good agreement with the conservation of amphipathic periodicity in H8 across the GPCR families^38^ (Figure S20A). Clustering of binding profiles of C-termini from individual receptors revealed a surprisingly large group of GPCRs (cluster 1) without membrane binding (Figure 6E+F). Several of these receptors are from the glutamate receptor family in agreement with the low average profile of the family (Figure 6D). However, many additional receptors belonging to other families and without any clear evolutionary origin, also show no membrane binding. Notably, this is true for many receptors belonging to the Rhodopsin family (Figure S19+20B). Clusters 2-5 showed membrane binding profiles consistent with membrane proximal H8, albeit with different intensities of the binding within each cluster and increasing length and distance from TM7 going from cluster 2 to 5 (Figure 6D-E and Figure S19). Finally, cluster 6 contained a wide range of different membrane binding profiles, which are more distant from TM7 and/or contain multiple membrane binding sites.

**Figure 6.**
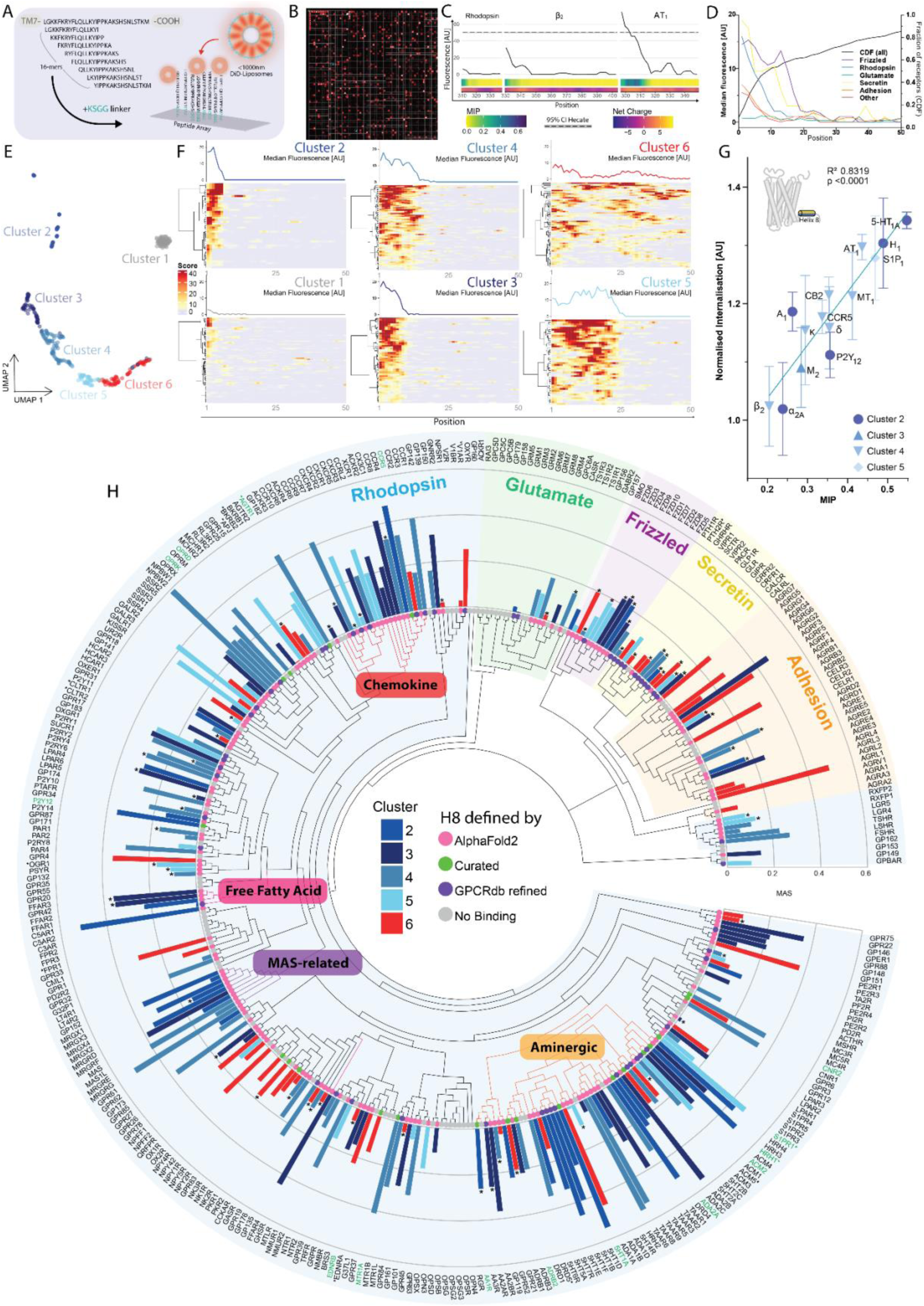
Proteome-wide characterization of C-terminal membrane interactions in GPCRs and their role in constitutive endocytosis of GPCRs. **(A)** Schematic of chip array for scanning of liposome binding strength throughout the full range of human GPCR C-termini at dual AA resolution. Individual peptides are randomly positioned on the chip. **(B)** Representative image of fluorescent signal from bound liposomes on zoomed in area of array. Regular pattern in top left corner is control Hecate peptide. **(C)** Compiled binding profile of the C-termini of Rhodopsin, β_2_ and AT_1_ along with corresponding MIP and net charge values displayed below. **(D)** Average liposome binding profiles of peptide sequences corresponding to 16 amino acid stretches of GPCR C-termini split by GRAFS family (left y-axis) and cumulative distribution function of all human GPCR C-terminals length (right y-axis) starting at 16 amino acids. **(E)** UMAP-based clustering of all human GPCR C-termini based on position and strength of liposome binding. **(F)** Heatmaps of all individual GPCR C-terminal liposome binding profiles within the six clusters and profile plots of the median fluorescence intensity (line above) (see S20 from detailed view). **(G)** Linear correlation (R^2^ = 0.83, p <0.0001) between MIP of H8 from clusters 2-5, based on sequences (Figure S22) in most pharmacologically relevant GPCRs and their respective endocytic rate (relative to Tac) determined by flow cytometry ^39^. Mean ± SEM, N=3. **(H)** Circular phylogenetic tree of human GPCRs based on sequence homology and divided into GRAFS families and clusters, alongside the corresponding calculated MIP based on H8 from either refined representative structures as annotated on GPCRdb (purple) ^11^, AlphaFold2 prediction (magenta) ^44^ or manually curated. Receptors used in (G) are highlighted by green and receptors described as mechanosensitive ^41^ are highlighted by asterisks (name). GPCRs from cluster 1 (no binding) in grey. Receptors with a structural H8 predicted longer than 20 amino acids marked with asterisk (bar).

We did not observe a significant enrichment of individual clusters within GPCR receptor families (Figure S20B), or specific tissues (Figure S20C). However, functional enrichment analysis within the studied GPCRs using Gene Ontology (GO) terms revealed a significant enrichment of receptors located to endolysosomal membranes for the bona fide H8 clusters (Clusters 2-5), while non-binding receptors (Cluster 1) are associated with disc-like membranes or membranes with topologies that restrict membrane insertion (Photoreceptor discs and exosomes, respectively) (Figure S21A). Cluster 6 receptors showed a weak association with caveola and growth cones, although driven by a relatively low number of proteins. A wide range of both Molecular Function and Biological Process (BP) terms were associated with all clusters (Figure S21B+C), leading us to perform a semantic similarity GO term enrichment analysis of the significant BP terms (Figure S21D). Consistent with the enrichment of Cluster 1 in Photoreceptor discs, we see a strong association with light related BP terms. Clusters 2-5 show the strongest association with synaptic transmission, particularly driven by receptors with membrane proximal H8 (cluster 2), while the longer and more distal H8 of cluster 5 associates with cellular differentiation processes. Finally, we find that receptors of cluster 6 associate with terms related to peripheral, cardiovascular regulation (Figure S21D).

### MIP of H8’s predicts constitutive internalization of human GPCRs

The experimentally derived membrane binding profiles enabled us to delineate bona fide membrane binding H8s (cluster 2-5) and calculate MIP values for these regions (Figure S22). We previously characterized the constitutive internalization of a range of pharmacologically relevant GPCRs ^39^ and here asked if this internalization could be explained by MIP values of the regions defined by their membrane binding profiles. Indeed, constitutive internalization significantly correlated with MIP values (cluster 2-5) (Figure 6G), demonstrating that the amphipathic nature of H8 can quantitatively predict constitutive endocytosis of full length GPCRs. Other biophysical measures, alternative to MIP, failed to explain constitutive internalization (Figure S23A-C). Consistent with the lack of effects of H8 substitution on agonist induced internalization (Figure 1H), MIP did not correlate with agonist-induced internalization (Figure S23D). The complex membrane binding profiles cluster 6 precluded extraction of MIP values predictive of constitutive internalization (Figure S23E). MIP is not significantly related to β-arrestin recruitment or reported G-protein coupling (Figure S24B+C). Together, these results indicate that the MIP of H8 selectively predicts constitutive internalization across this functionally important and structurally well-characterized subset of GPCRs.

Given the functional importance of constitutive internalization for dynamic surface expression of GPCRs, we utilized MIP of H8 as a proxy to map predicted internalization rates across all non-odorant GPCRs belonging to cluster 2-5 (Figure 6H and S23F). High MIP values are predominantly observed as phylogenetically related clusters within the rhodopsin family of receptors, including receptors for biogenic amines, chemokine receptors, and MAS-related receptors as well as free fatty acid receptors, alongside individual receptors such as NPBW2 and C5AR1. We also note that Secretin and Frizzled receptors largely contained a H8 predicted or found to be longer than 20 residues and the functional significance of this remains to be explored. MIP in GPCRs associated with Fast Endophilin-Mediated Endocytosis (FEME), that mediate endosomal signaling or function as mechanosensors display H8 MIP value distributions that do not deviate significantly from average H8 MIP for all GPCRs (cluster 2-5). On the other hand, GPCRs associated with primary cilia, which exhibit topologies with an inverse curvature to endocytic structures, display H8 sequences with significantly lower MIP (Figure S24A). Conversely, GPCRs expressed by viruses, which are known to elicit high constitutive internalization ^40^, mostly have high MIP values of their H8.

### Membrane insertion propensity is evolutionary conserved across distant organisms

To assess potential functional conservation of high MIP of H8, we used available data from UK biobank on naturally occurring variants in human GPCRs. While the cost of individual amino acid substitutions was similar in the H8 region compared to the remaining part of the C-termini (Figure 7A), the frequency of non-synonymous natural variants was significantly lower within H8s (Figure 7B). When analyzing differences in MIP resulting from these naturally occurring variants, we found that the observed substitutions give rise to a more conserved MIP than substitutions not observed (Figure 7C). Finally, the frequency of natural variants within H8 for individual receptors declines with increasing MIP, emphasizing the functional conservation of high MIP (Figure 7D).

**Figure 7.**
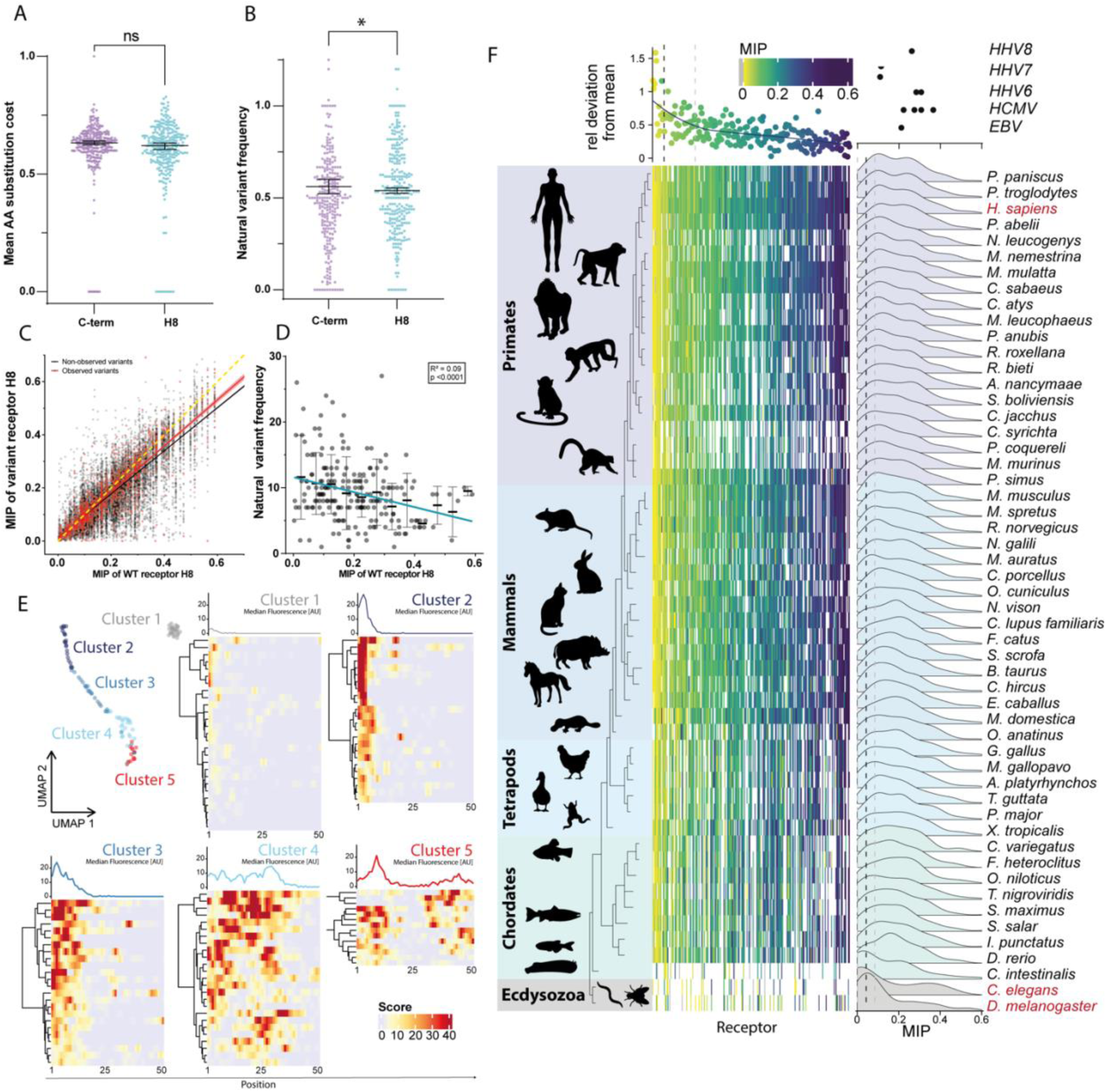
Functional conservation and evolutionary refinement of MIP of H8 in GPCRs. **(A)** Normalised BLOSUM62 ^45^ substitution cost of natural variants mapped to C-term vs H8. **(B)** Frequency of natural variants in the C-term vs H8 of human GPCRs of clusters 2-6 identified in individuals registered in the UK Biobank ^46^. **(A and B)** Wilcoxon non-parametric statistical test applied. Asterisks indicate p value <0.05 and ns: p value >0.05. **(C)** Scatter plot of the calculated MIP values of all identified non-synonymous natural variants (red dots) identified in the UK Biobank ^46^ and non-identified possible non-synonymous substitutions (black dots) as a function of the receptors wildtype MIP. Linear regression analysis of observed natural variants with a slope of 0.87 ±0.01 is closer to full conservation (Indicated by dashed yellow line, slope of 1) than for non-observed 0.81 ±0.004 (all other possible variants). This highlights a relative conservation of MIP. **(D)** Scatter plot showing frequency of non-synonymous natural variants of individual GPCRs as a function of MIP (each receptor represented by a dot), with a negative linear correlation indicating lowered variant frequency for receptors with high MIP (Cyan, R^2^ = 0.089, p <0.0001). Lines indicate average frequency of binned MIP intervals +/- SD. **(E)** UMAP-based clustering of combined *D. melanogaster*, *C. elegans* and *S. cerevisiae* GPCR C-termini based on position and strength of liposome binding motifs. Heatmaps of individual GPCR C-terminal liposome binding profiles within the 5 clusters and profile plots of their median fluorescence (Top) (See Figure S24 for list of individual receptor and species). **(F)** Evolutionary tree of GPCRs displaying MIP of H8 sequences identified in Human receptors and aligned across species (left). The ratio of variance vs. mean of all MIP decrease with increasing MIP (upper, left). (Upper, right) MIP values for viral GPCRs (Table S2). (Lower right) Density distribution of MIP values for individual species shift to the right during evolution.

Finally, we assessed the evolutionary origin and refinement of MIP in H8 of GPCRs. Using the membrane binding array, we found that GPCR C-termini from evolutionary distant organisms, including C. elegans, D. Melanogaster and S. Cerevisiae, show membrane binding profiles akin to H8 in human GPCRs (cluster 2 and 3) as well as non-binding (cluster 1) and more complex binding profiles (cluster 4 and 5) (Figure 7E and S25). This enabled us to address evolutionary refinement by aligning H8 sequences, as defined in human, across species back to Ecdysozoa (Figure 7F).

Individual receptors generally show strong conservation of MIP throughout evolution as indicated by the overall preservation of MIP ranking from yellow to purple across species. The relative variation across species is lowest for H8s with high MIP indicating evolutionary conservation of this feature of H8 (Figure 7F, top). Next, we assessed the distribution of MIP values for individual species across evolution (Figure 7F, histograms to the right). Intriguingly, in ecdysozoa the distribution of MIP values displays a single peak overlaying with values obtained from randomly generated sequences (indicated by the dashed line). However, the fraction of receptors with high MIP of H8 (> 0.3) gradually increases in higher organisms indicating functional significance (Figure 7F). This functional selection and refinement are evident from the multimodal distribution of H8 MIP values across human GPCRs, significantly different from a normal distribution (Figure 4H, p <0.0001, Shapiro-Wilk). Together, these findings demonstrate both evolutionary selection and refinement as well as functional conservation of the MIP value of H8 in GPCRs.

## Discussion

Here we show that AHs facilitate TMP association with areas of high membrane curvature similar to what has been demonstrated for cytosolic proteins. For TMPs, this association accelerates their constitutive trafficking in a manner that correlates with the MIP of their AHs. While this paper focused on endocytosis, we show evidence that this mechanism, due to the basic biophysical principle, is likely to apply to trafficking events throughout the endomembrane system involving highly curved membrane topologies and serve as a key determinant for steady-state distribution of GPCRs. We term this mechanism ToTAM.

Mechanistically, ToTAM is characterized by fast, cooperative cargo recruitment into small, spherical, high-density carriers and thus ToTAM displays elements of both membrane curvature sensing and facilitation of the cargo/carrier complex formation. In fact, ToTAM increases endocytosis of the transferrin receptor in trans, implying an active impact on endocytic processes. During this process, partial biochemical association with CME was observed, however, actual CLC accumulation prior to scission was low and increasingly asynchronous with higher MIP of the AHs. Together, our data suggest that ToTAM might hijack nucleation involving CLC and EPS15 to shortcut classical CME in an autonomous fashion that is complementary to the established adaptor based endocytic machinery.

The impact of high MIP H8 specifically on constitutive, and not agonist-induced, internalization of full length GPCRs, was causally demonstrated by experiments with chimeric receptors. Using large-scale liposome-binding data to exclude GPCRs with C-termini that did not display membrane-binding or complex membrane binding enabled us to correlate H8 MIP with full- length constitutive internalization across all human GPCRs. Highest constitutive internalization was predicted for the groups of chemokine receptors, biogenic amine receptors, MAS-related receptors, and free fatty acid receptors. It will be interesting to elucidate the functional implication of the predicted high constitutive turnover with e.g. ligand scavenging, membrane modulating ligands or continuous receptor availability.

Another recently described membrane-associated function of H8 is its role in conferring mechanosensitivity to GPCRs ^41^. Several of these receptors include a H8 with high MIP, including H_1_, AT_1_, and S1P_1_. A putative interplay between mechanosensation and high constitutive internalization due to membrane curvature sensing might reflect two aspects of the same biophysical principle and functional implications of these might unveil themselves perhaps in the context of the cardiovascular system. Moreover, GPCRs implicated in viral entry, as well as virally expressed GPCRs displaying high constitutive internalization, all except mGluR2 show high MIP ^42^, potentially advocating a novel molecular target for anti-viral therapy. Finally, the simple molecular nature of ToTAM is compatible with an early evolutionary origin, dating back prior to the yeast GPCRs characterized here, conceivably even prior to adaptor based trafficking. Ultimately, functional studies will be needed to settle the evolutionary relation between these two complementary trafficking mechanisms.

## Acknowledgments

We acknowledge the Proteomics Research Infrastructure, Core Facility for Integrated Microscopy and Flow Cytometry and Single Cell Analysis facilities at the University of Copenhagen. Further, we would like to thank Nabeela Khadim for technical assistance.

## Author contributions

Conceptualization: KLM, UG, JHS, RH

Methodology: KLM, JHS, RH, NRC, ALE, MDL, JR, AHL, MS, MP, KAA, CKH, JPC, DP, PB, VH, ASH Investigation: KLM, JHS, RH, JR, AHL, MS, MP, KAA, MDL, ALE

Visualization: KLM, JHS, RH, JR, AHL, MS, TTEN, DP

Funding acquisition: KLM, UG, AHL, PB, DP, VH Project administration: KLM

Supervision: KLM, DP, PB, CKH Writing – original draft: KLM, JHS, RH

Writing – review & editing: KLM, JHS, RH, JR, FH, AHL

