## supplementary figures and methods for "Membrane curvature association of amphipathic helix 8 drives constitutive endocytosis of GPCRs"

### Supplementary Materials

The following supplementary information is available for this paper.

Methods

Figures S1 to S24

Tables S1 and S2

Mathematical model “Acidification Kinetics Function”

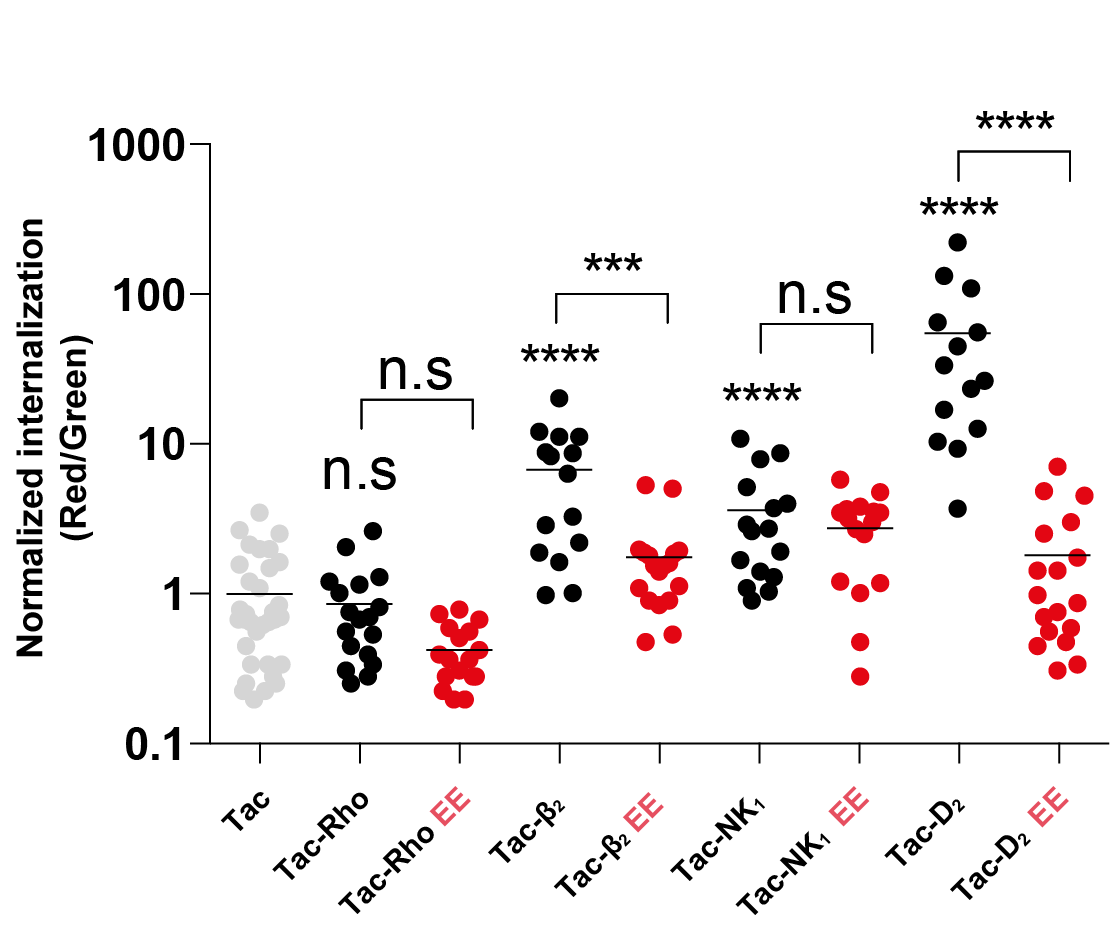

Figure S1. AHs mediate internalization of Tac in HeLa cells.

Quantification of internalization (Red/Green) from confocal images in HeLa cells (every dot represents a cell, compiled from three independent experiments). Statistical tests using Tukey’s multiple comparison. Four asterisks indicate p < 0.001 and three indicate p <0.01.

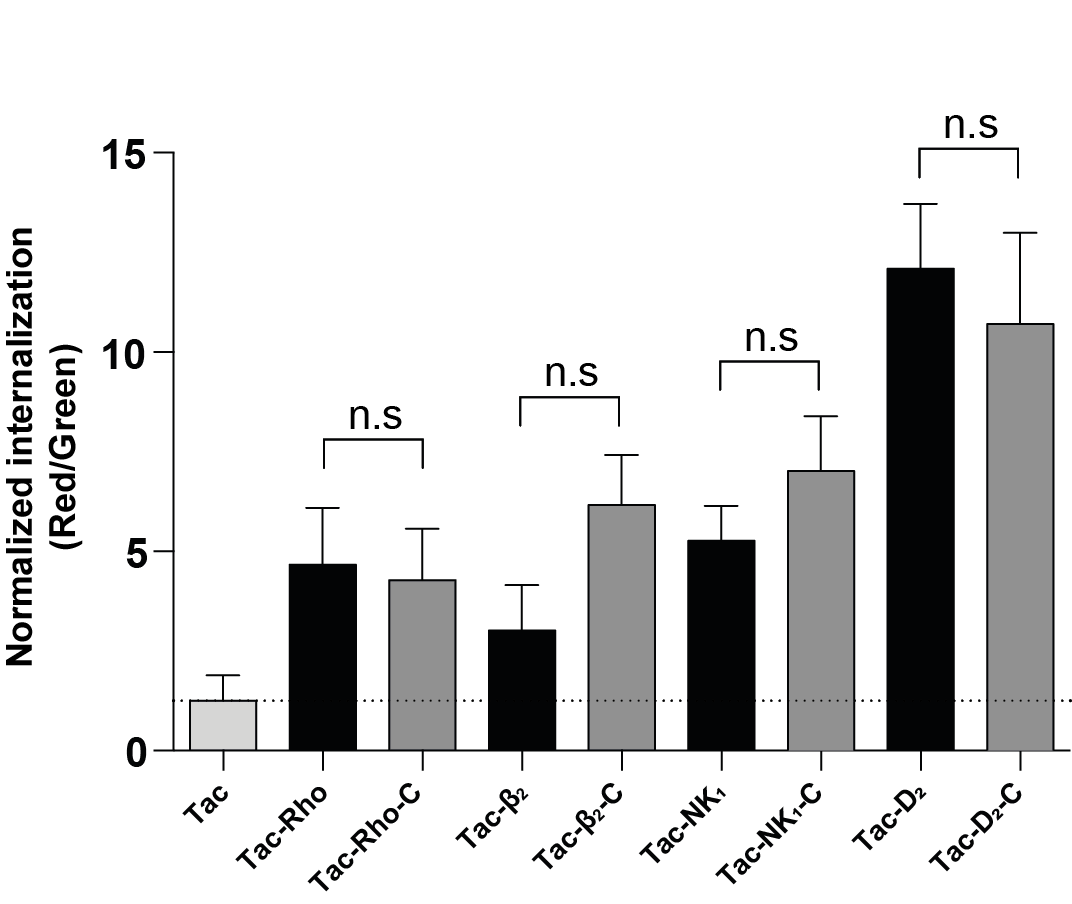

Figure S2. Internalization of Tac by AHs is independent of putative palmitoylation on C-terminal cysteines.

Quantification of internalization (Red/Green) from confocal images of HEK293 cells comparing Tac-AH constructs without and with truncation at the C-terminal cysteines. Mean ± SEM, N=3. Statistical tests using Tukey’s multiple comparison.

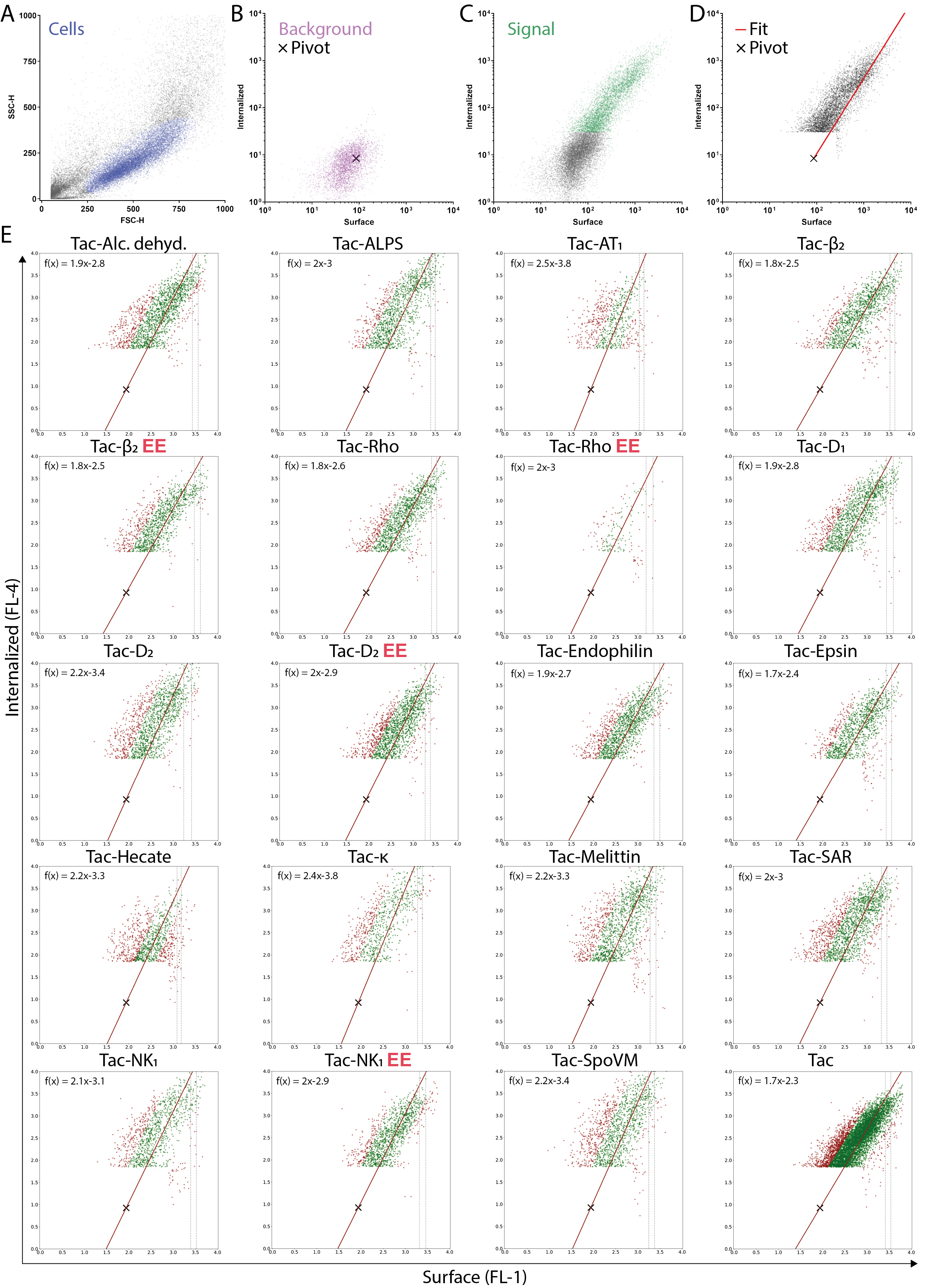

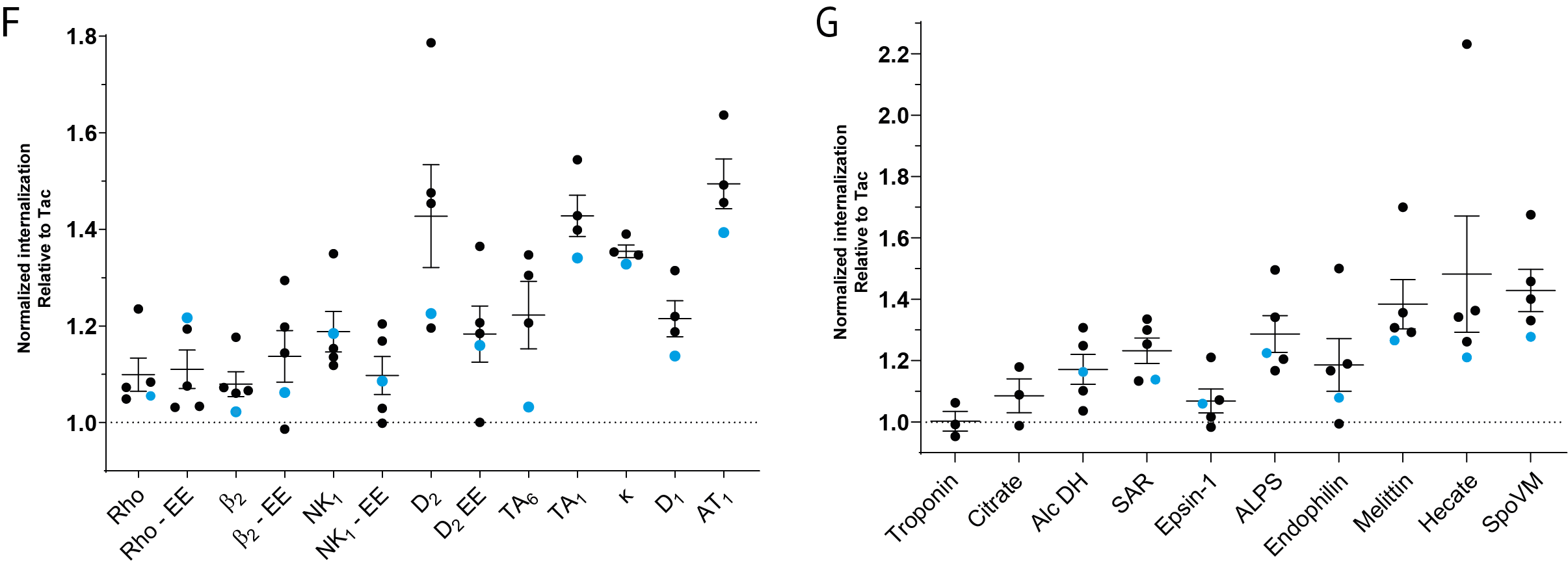

Figure S3. Flow cytometry analysis of antibody feeding assay using Tac-AH constructs.

**(A)** Initial gating of cell population given forward and side scatter signal.

**(B)** Determination of background fluorescent signal obtained from empty pcDNA3 transfected cells and pivot point for linear regression fitting. Each point represents a single cell.

**(C)** Representative signal in the two fluorescent channels split into background (gray, defined as 99-percentile of empty pcDNA3 transfected cells) and specific (green) signal.

**(D)** Specific signal after removal of background signal alongside linear fit fixed by the pivot point.

**(E)** Specific signal and fits (corrected using RANSAC; green dots remain, red dots are considered outliers) of the ratio of internalized vs. surface obtained from a representative run of Tac-AH construct in Figure 1.

**(F+G)** Normalized internalization ratios of Tac-AH constructs from (e). The representative flow cytometry run shown in (e) are highlighted in blue alongside remaining experimental replicates (black).

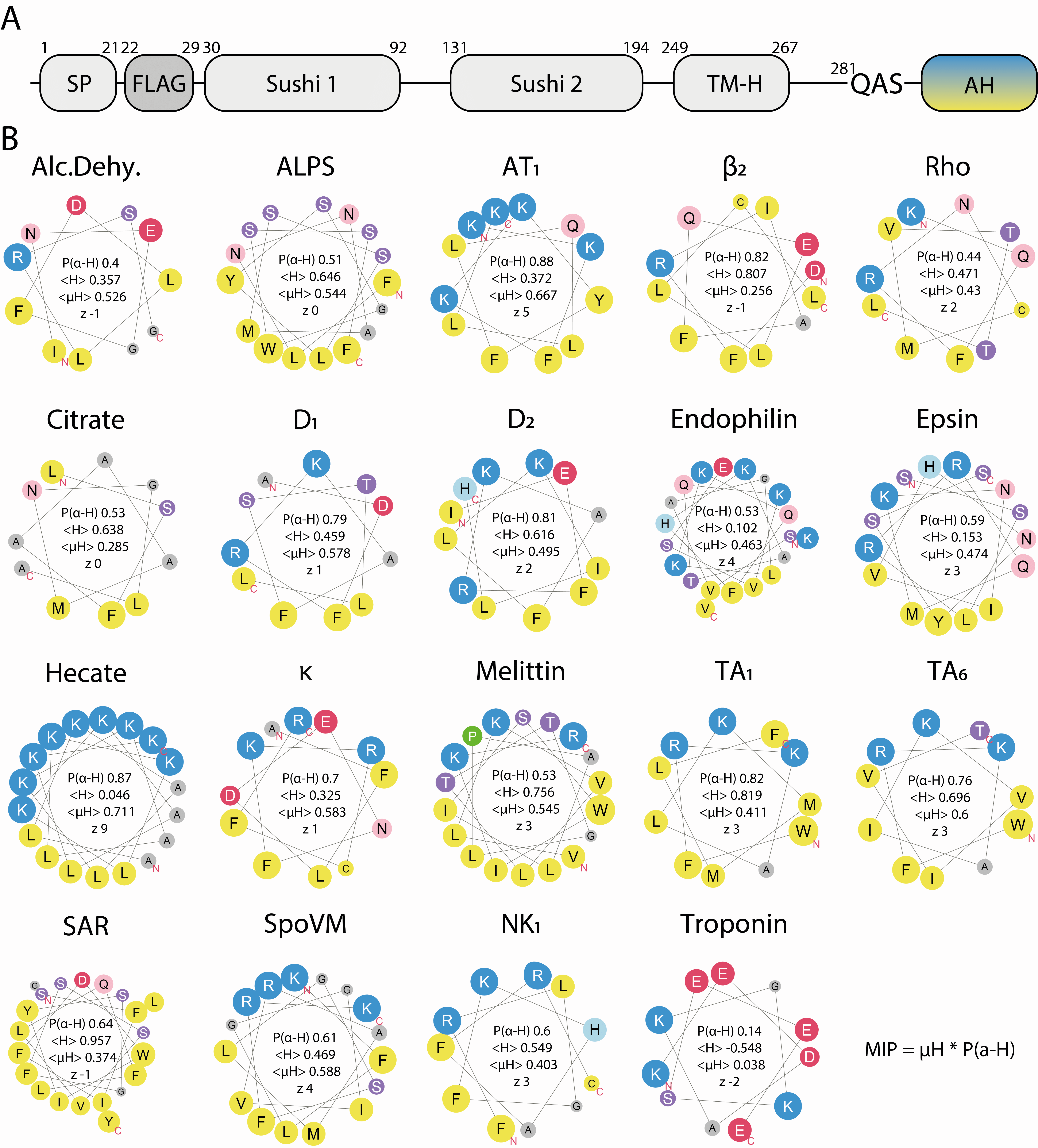

Figure S4. Helical wheel representation of the amphipathic helices and their biochemical parameters.

(A) Schematic outline of the Tac-AH constructs, with the N-terminal signal peptide (SP), Flag epitope tag (FLAG), the two extracellular Sushi domains, transmembrane helix (TM-H) and the different amphipathic helices (AH) inserted after the QAS sequence in the intracellular C-terminus of the construct.

(B) Helical wheel representations obtained using Heliquest ^43^ of the AHs in Figure 1 in alphabetical order alongside MIP calculation. Chemical nature of amino acid side chains is color coded (Blue, positively charged; Red, negatively charged; Green, constrained; Grey, small non-polar; light red/purple, hydrophilic; Yellow, hydrophobic. Size of circles indicate size of the side chain. P(a-H): a-helical propensity (obtained by NetsurfP); <H>: mean hydrophobicity; <µH>: hydrophobic moment; z: net charge.

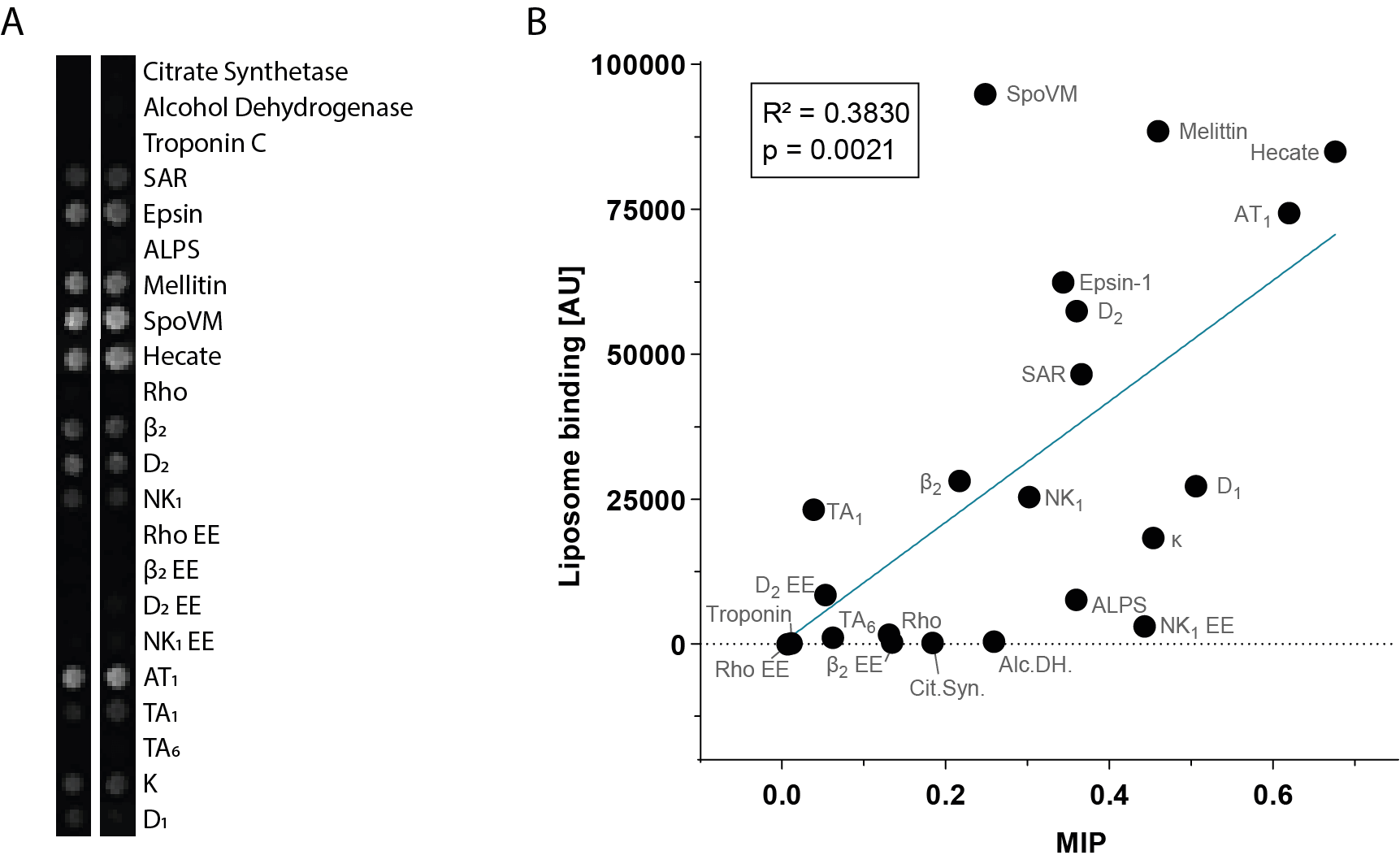

Figure S5. Correlation of MIP and experimentally determined membrane binding strength.

**(A)** Technical replicates showing binding of fluorescently labelled (DiD) liposomes made of bovine brain extract (Folch’s fraction type I) on a SPOT array with peptides corresponding to amphipathic helices.

**(B)** Liposome binding as a function of MIP predicted for individual AHs.

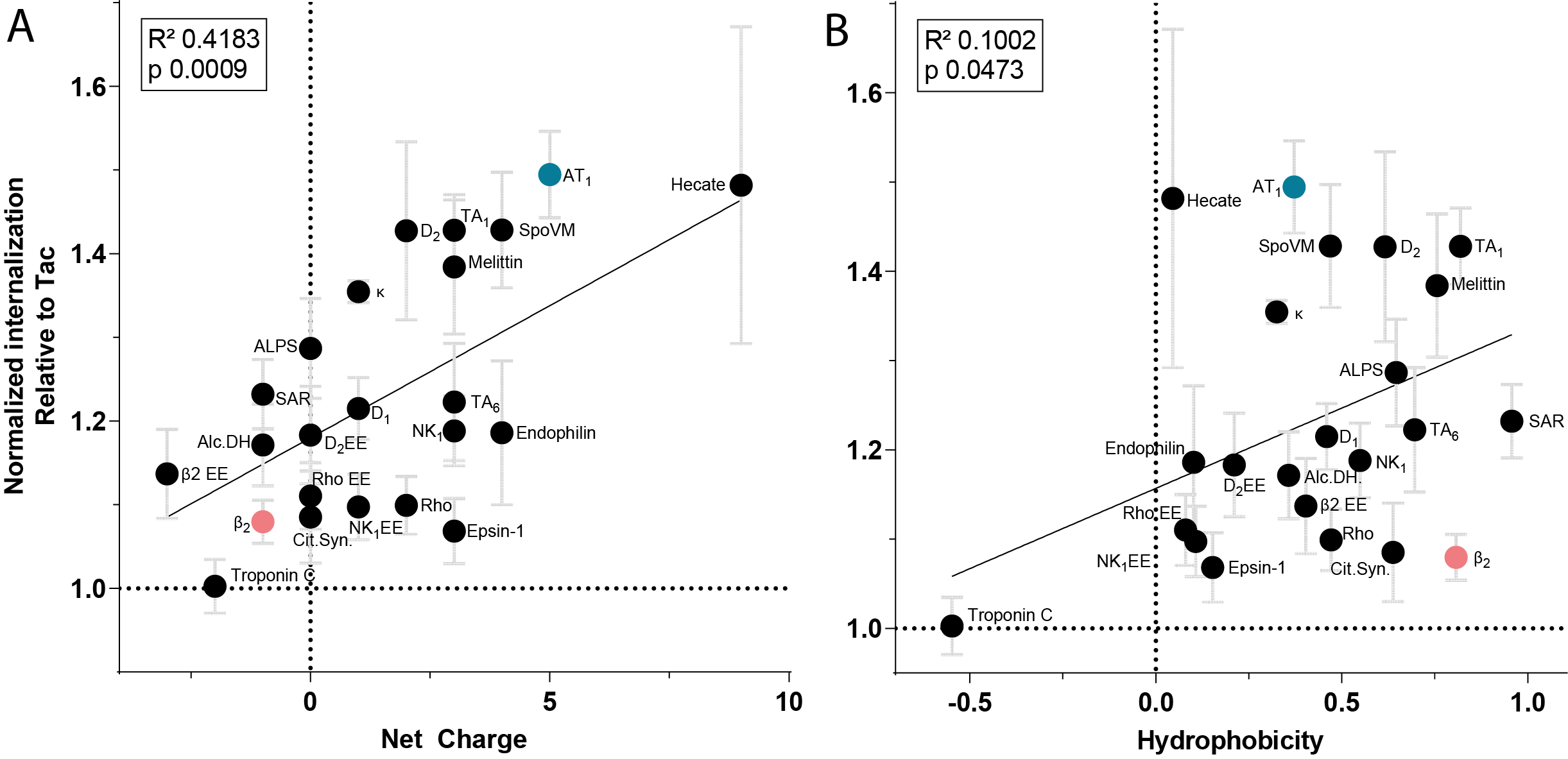

Figure S6. Relation of biophysical factors of AHs and their internalization.

**(A+B)** Linear correlation between net charge (z) (a) and mean hydrophobicity (<H>) (b) of AHs inserted in Tac and respective endocytic rates (relative to Tac) as in Figure 1F. Dashed vertical lines indicate 0 values.

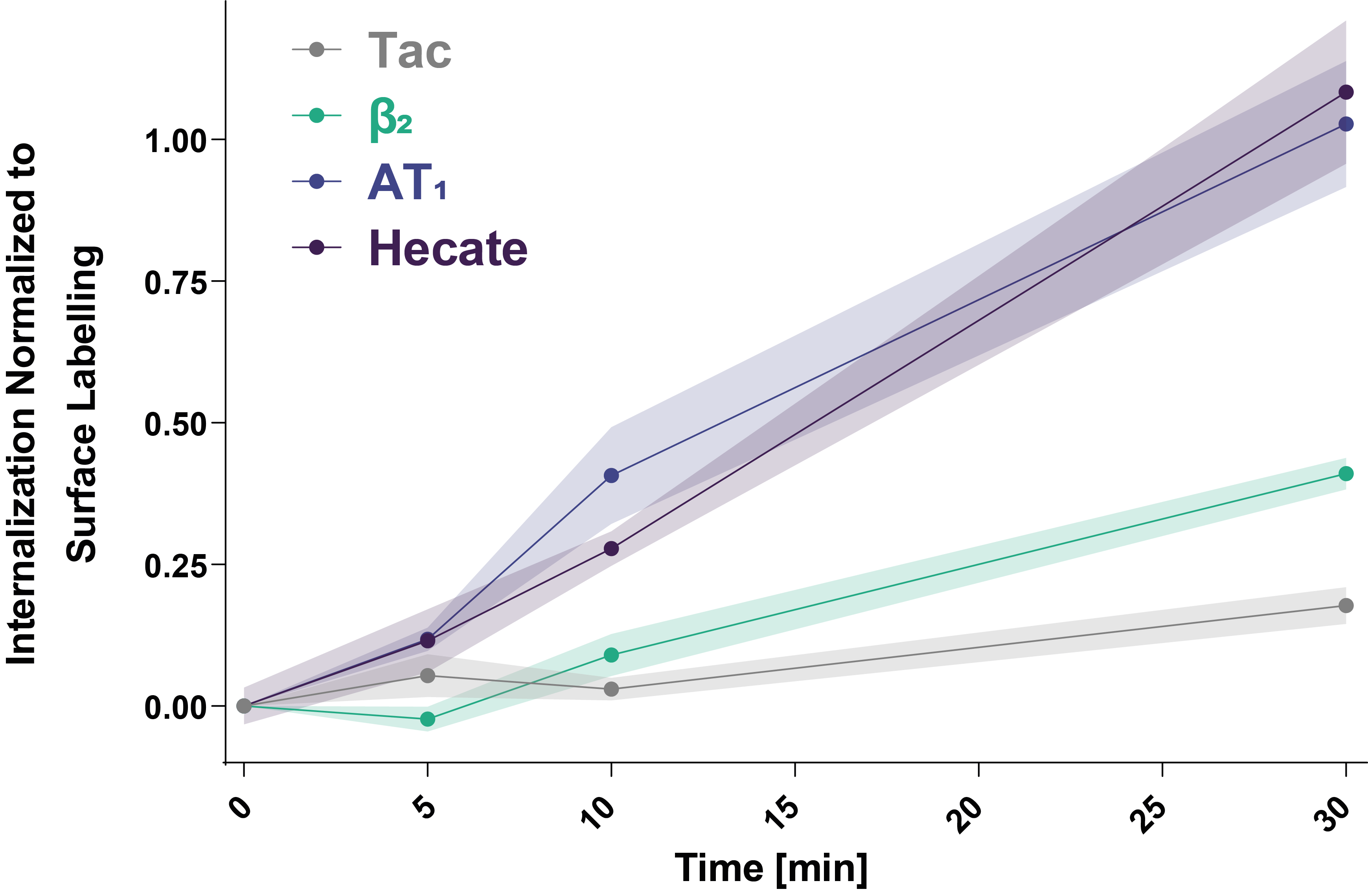

Figure S7. Time series of surface labelled Tac-AH internalization supports an active process.

Relative internalization of surface labelled Flag-Tac-AH constructs over a time-series of 0, 5, 10, and 30 minutes in the presence of Monensin in HEK293 cells as examined by flow cytometry. Relative internalization as the fluorescence measured after stripping of surface bound antibodies, normalized to a surface labelling control of the equivalent construct, and corrected for strip treatment. Results are representative of two experiments with 3 replicates each. Shaded area indicates SEM.

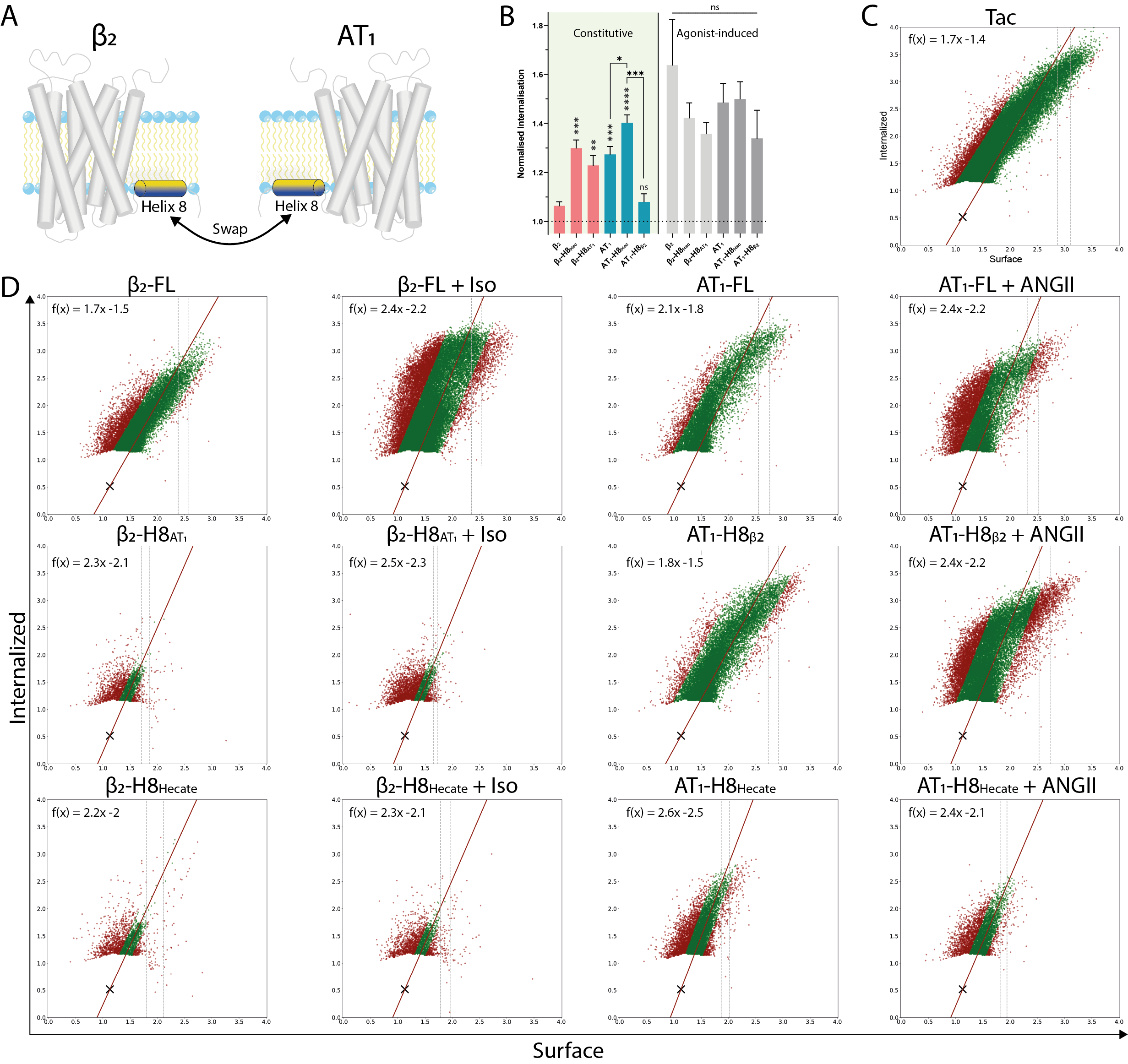

Figure S8. Flow cytometry analysis of full-length chimeric GPCRs.

**(A)** Schematic outline for the exchange of amphipathic H8 between β_2_ and AT_1_ receptors.

**(B)** Internalization rates of WT and chimeric receptors from Figure 1.

**(C+D)** Specific signal from a representative run after removal of background gated events with a RANSAC corrected linear correlation fit (green dots remain, red dots are considered outliers) for Tac (c) and the chimeric GPCR constructs in absence and presence of the respective agonists as indicated (d). Iso; isoproterenol, ANGII; Angiotensin II.

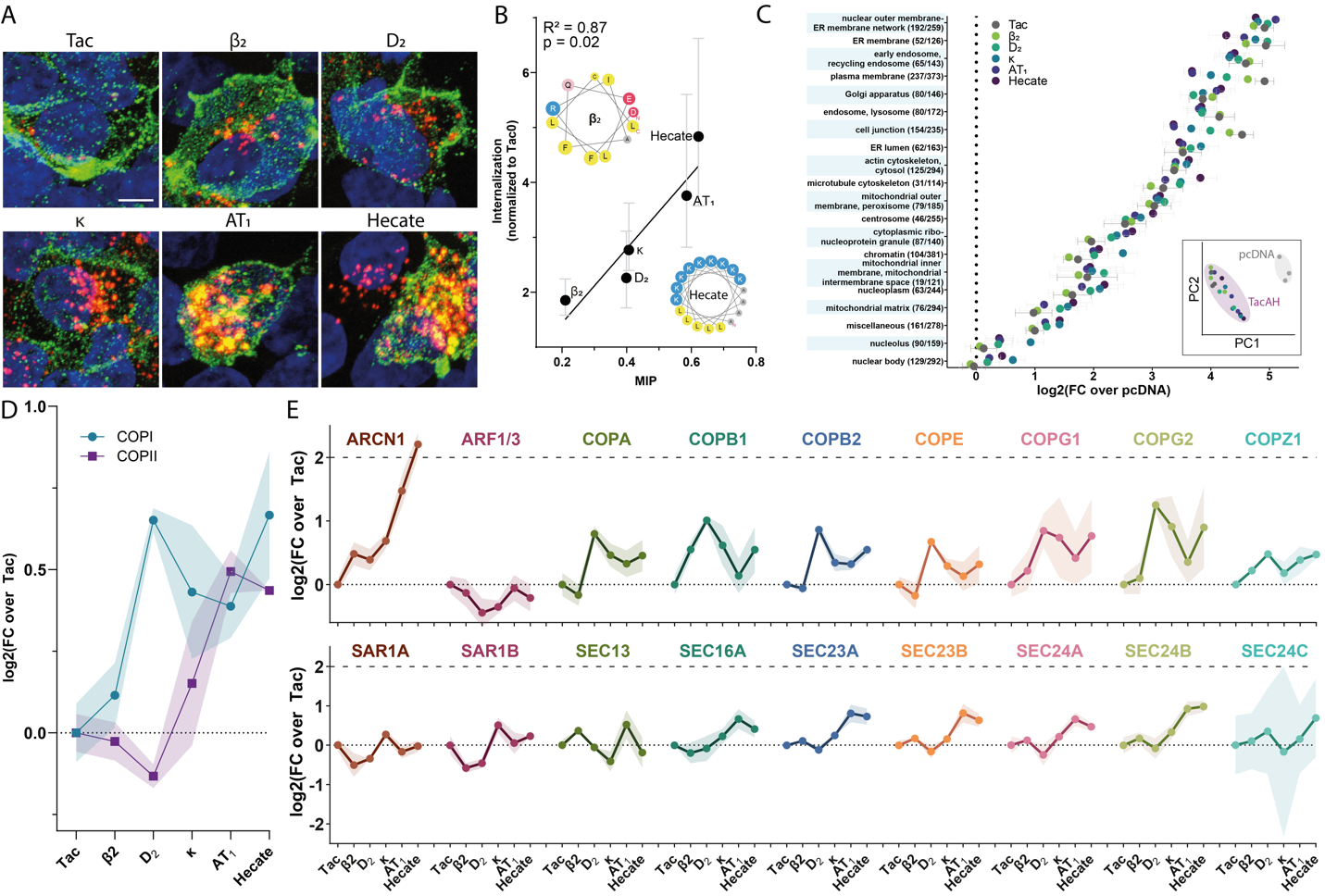

Figure S9. Association of TacAH-BioID2 constructs driven by ToTAM with coat associated proteins.

**(A)** Representative images of HEK293 cells transfected with Tac-H8-BioID2 constructs and subjected to antibody feeding (Green, surface; Red, internalized).

**(B)** Linear regression analysis of internalization as a function of predicted MIP shows preserved correlation for the BioID2 fusion constructs.

**(C)** Difference in abundance levels (log2 fold change) of TacAH-BioID2 constructs relative to pcDNA control for identified proteins belonging to subcellular localization groups used in ^26^ (MMF). (insert) PCA biplot showing variance of abundance levels in TacAH-BioID2 constructs versus pcDNA control.

**(D+E)** Difference in abundance levels (log2 fold change) relative to Tac for identified COPI+II machinery pooled (d) and individually (e).

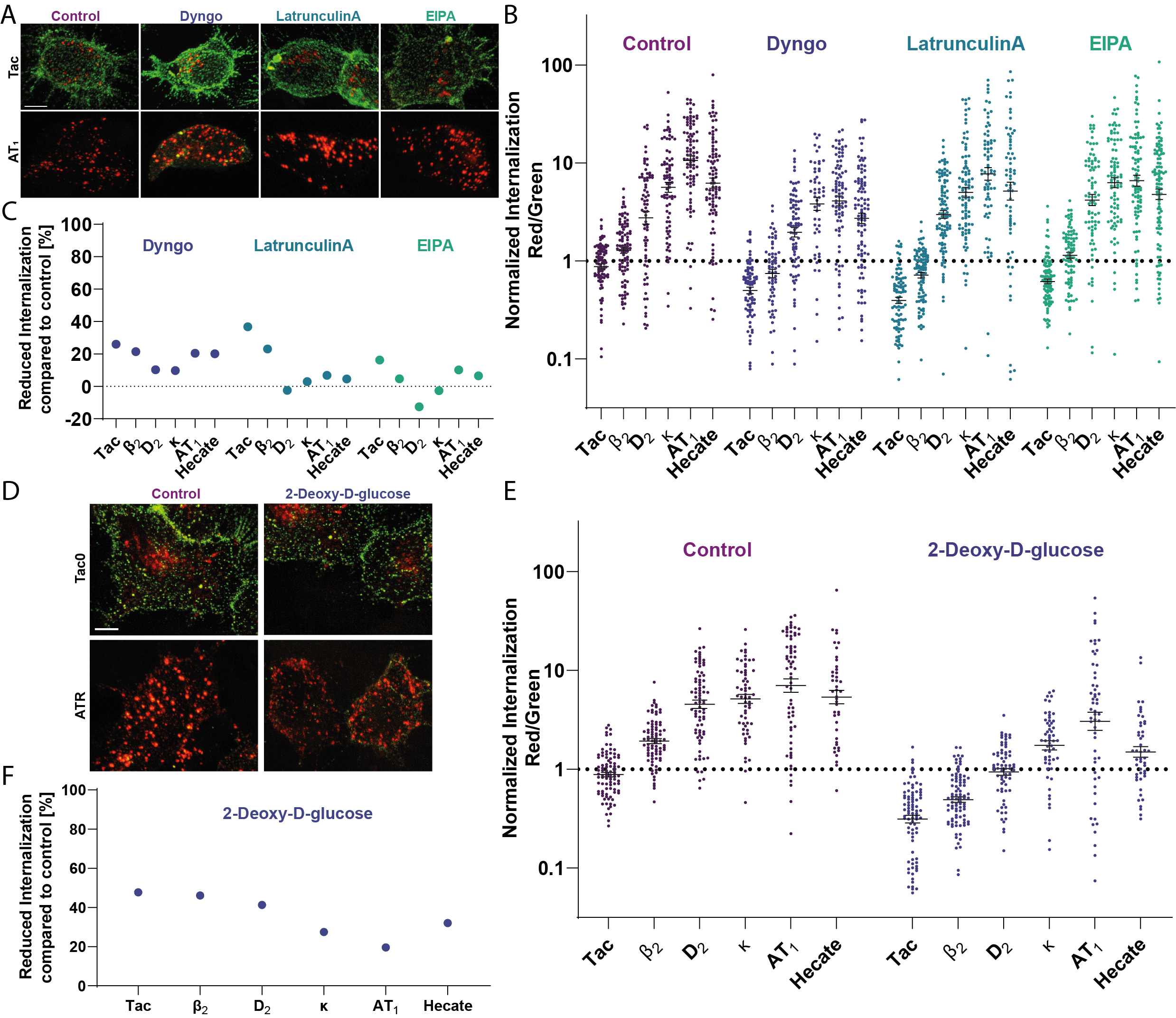

Figure S10. Pharmacological interference of endocytic machinery during ToTAM.

**(A)** Representative images of HEK293 cells transfected with Tac-AH constructs and subjected to antibody feeding (Green, surface; Red, internalized) in absence or presence of pharmacological inhibitors of endocytosis (10µM Dyngo; dynamin, 200nM LatrunculinA; actin polymerization, 1µM EIPA; macropinocytosis).

**(B)** Quantification of internalization (Red/Green) from confocal images (every dot represents a cell, compiled from three independent experiments).

**(C)** Quantification of internalization shown as reduction (in percentage) compared to non-treated cells (averages from b).

**(D)** Representative images of HEK293 cells transfected with Tac-AH constructs and subjected to antibody feeding (Green, surface; Red, internalized) during metabolic starvation with 10mM glycolysis resistant 2-Deoxy-D-glucose (ATP-depletion) vs 10mM D-glucose (Control).

**(E)** Quantification of internalization (Red/Green) from confocal images (every dot represents a cell, compiled from three independent experiments).

**(F)** Quantification of internalization shown as reduction (in percentage) compared to control cells (averages from e).

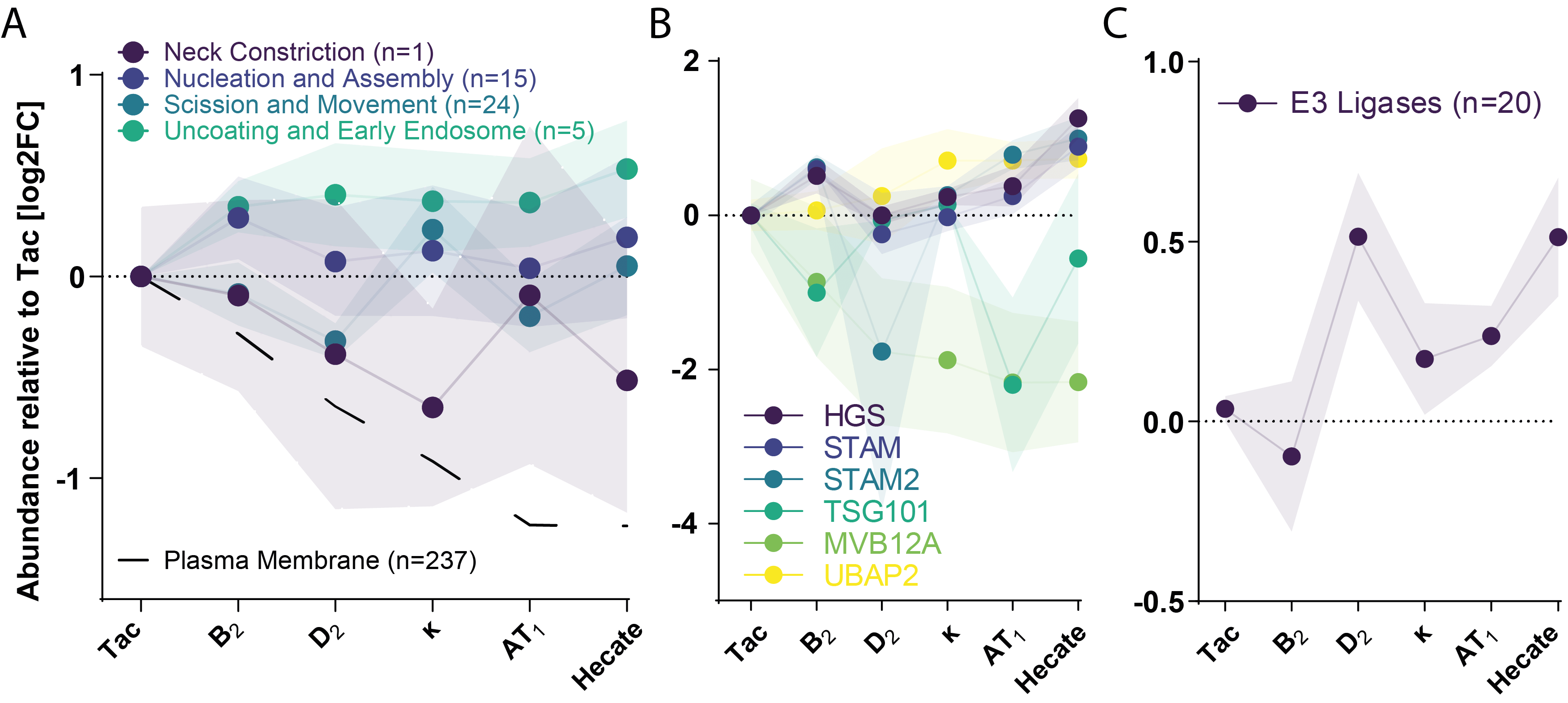

Figure S11. ToTAM driven association with other co-adaptor processes.

**(A-C)** Difference in abundance levels (log2 fold change) relative to Tac for all identified proteins of temporally defined modules of CME in ^31^(A), ubiquitin binding proteins of ESCRT machinery identified as listed in ^33^(B) and all identified E3 ubiquitin ligases (C).

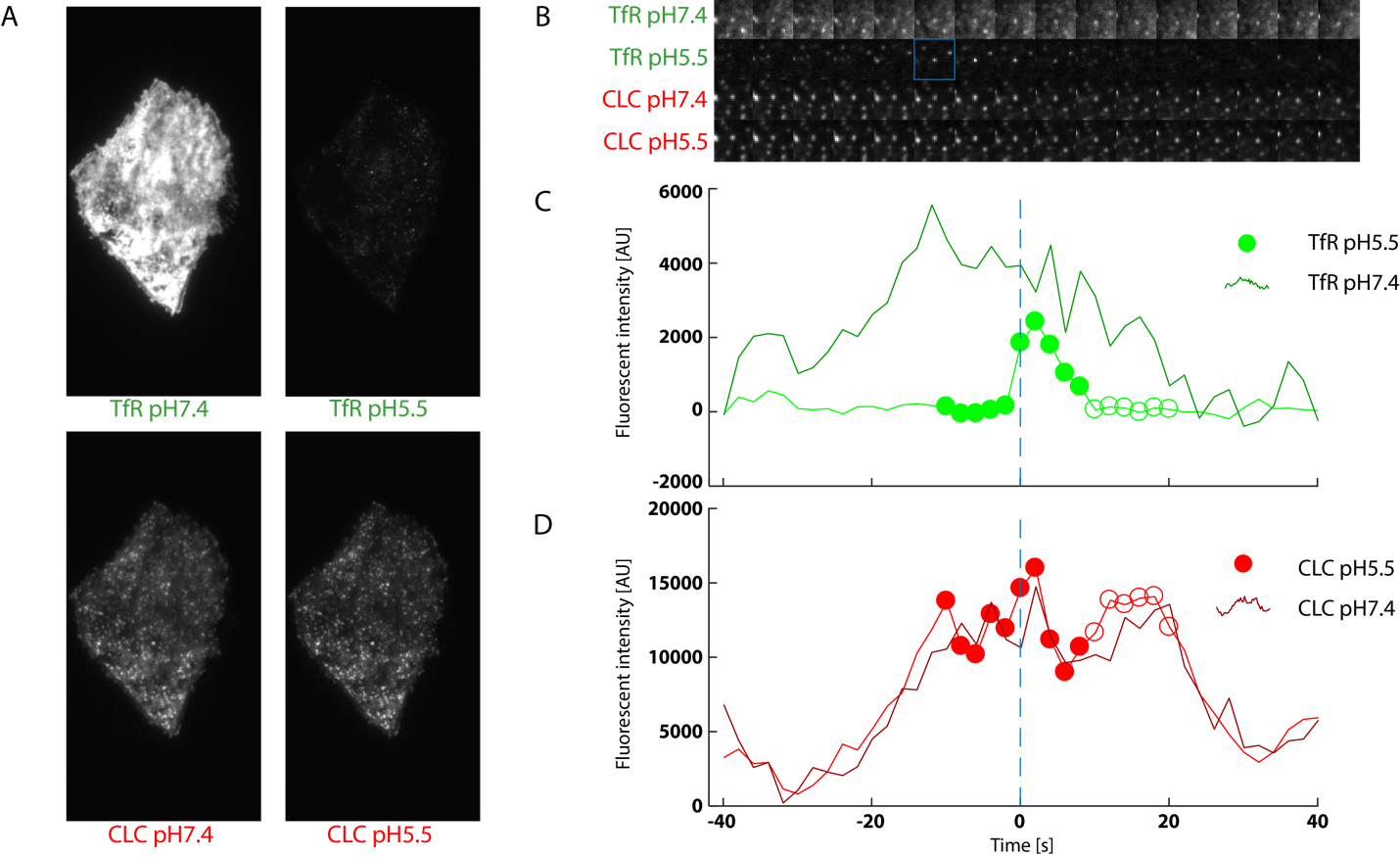

Figure S12. Example and outline of pulsed pH TIRF microscopy and analysis.

**(A)** Exemplary wide field micrograph, at initiation of experiment, of a BSC-1 cell expressing TfR-SEP (green legend) and CLC-mCherry (red legend) at pH 7.4 and 5.5 as indicated.

**(B)** Series of images taken with 2 sec intervals showing the fluorescent signal of TfR-SEP and CLC-mCherry leading up to and following an individual endocytic scission event (indicated by blue square in the TfR-SEP pH 5.5 channel).

(**C+D)** Quantification of the fluorescent traces of the pH 5.5 and pH 7.4 channel for TfR-SEP (**c**) and CLC-mCherry (**d**). Dots correspond to the timepoints shown in (**b)**.

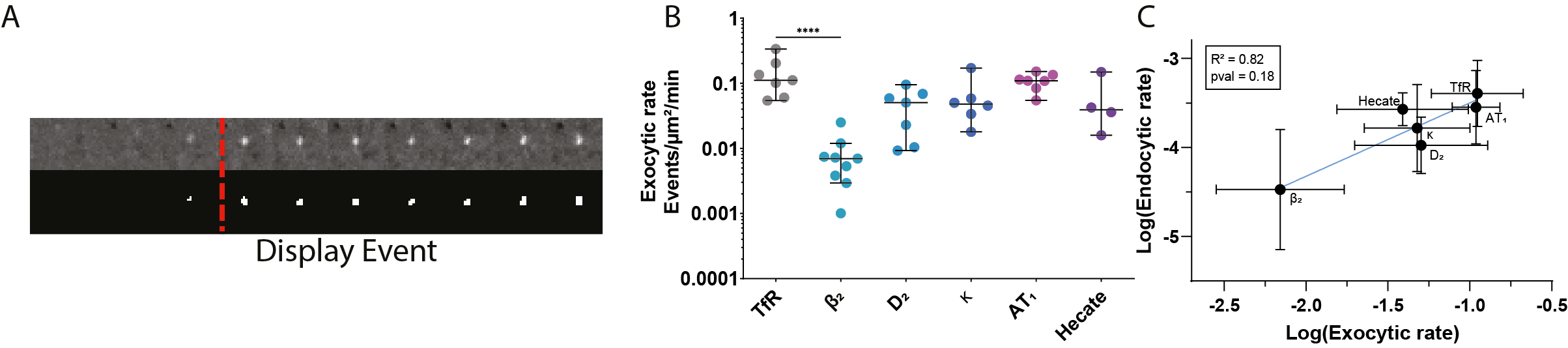

Figure S13. TacAH constructs with high MIP recycle efficiently.

**(A)** Representative images of exocytic events (see ^35^). Upper row shows SEP fluorescence during the progression of the exocytic event through eleven frames (four pre- and seven post-scission, as indicated by dashed lines). Lower row shows the mask used for quantification of the progression of these events.

**(B)** Exocytic rate quantified across cells expressing six different SEP-Tac-AH constructs. Each dot represents a cell. Statistical analysis using Kruskal-Wallis multiple comparisons test.

**(C)** Correlation measured between log-transformed exocytic-, and endocytic-rates for cells expressing either of the six selected SEP-Tac-AH constructs (R^2^=0.82, p<0.05).

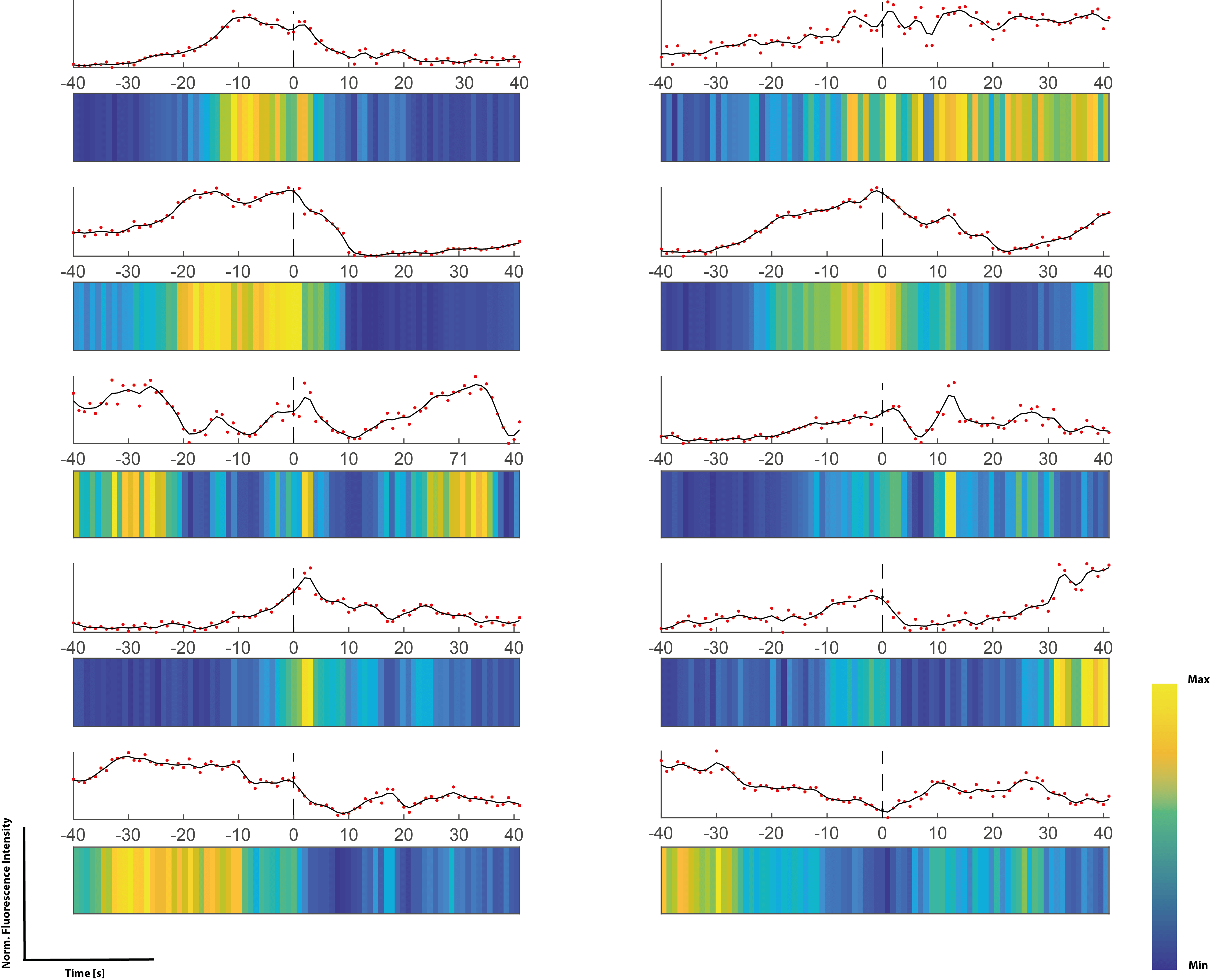

Figure S14. Normalized fluorescence traces of CLC-mCherry representing distinct activity profiles surrounding individual endocytic events.

(Upper) Quantification of single CLC-mCherry fluorescence spots from ROI defined by endocytic events in the TfR5.5-channel (qualified by scission as previously described ^35^. Red dots represent obtained the quantified fluorescence at each time-point, whereas the black line represents a smoothened linear model based on loess regression (black line). (Lower) The smoothened trace converted into a row in the raster-plots. Colormap indicates the values of smoothened trace at each timepoint.

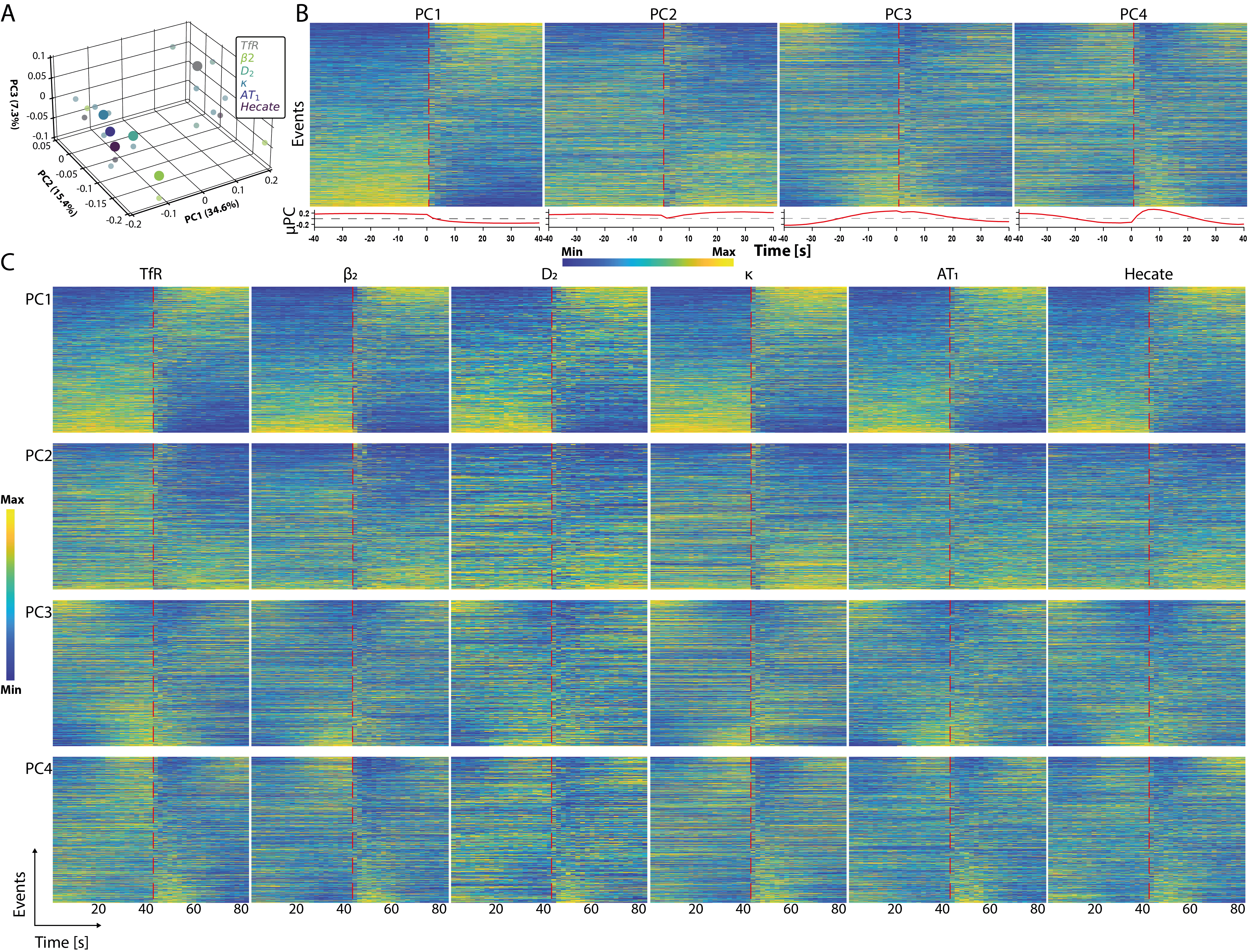

Figure S15. Rasterplots of CLC-mCherry traces sorted according to the four main principal components.

**(A)** Individual SEP-Tac-AH constructs evaluated based on a principal component analysis with average values of the three main components in a 3D-scatter plot. Larger dots represent the 3-dimensional locations, while the smaller shadow-dots indicate the projection on each 2D-plane to enable better comparison with regard to each component.

**(B)** Heatmaps of all CLC-trace profiles pooled together and sorted by each of the first four principal components. Black lines indicating time of scission. Below each heatmap is a representation of the principal component used to sort the global pool of traces.

**(C)** Rasterplots of CLC-trace morphologies collected from cells expressing each of the six SEP-Tac-AH constructs and sorted by each of the four main principal components. Heatmap values indicate values of the individual smoothened and normalized traces.

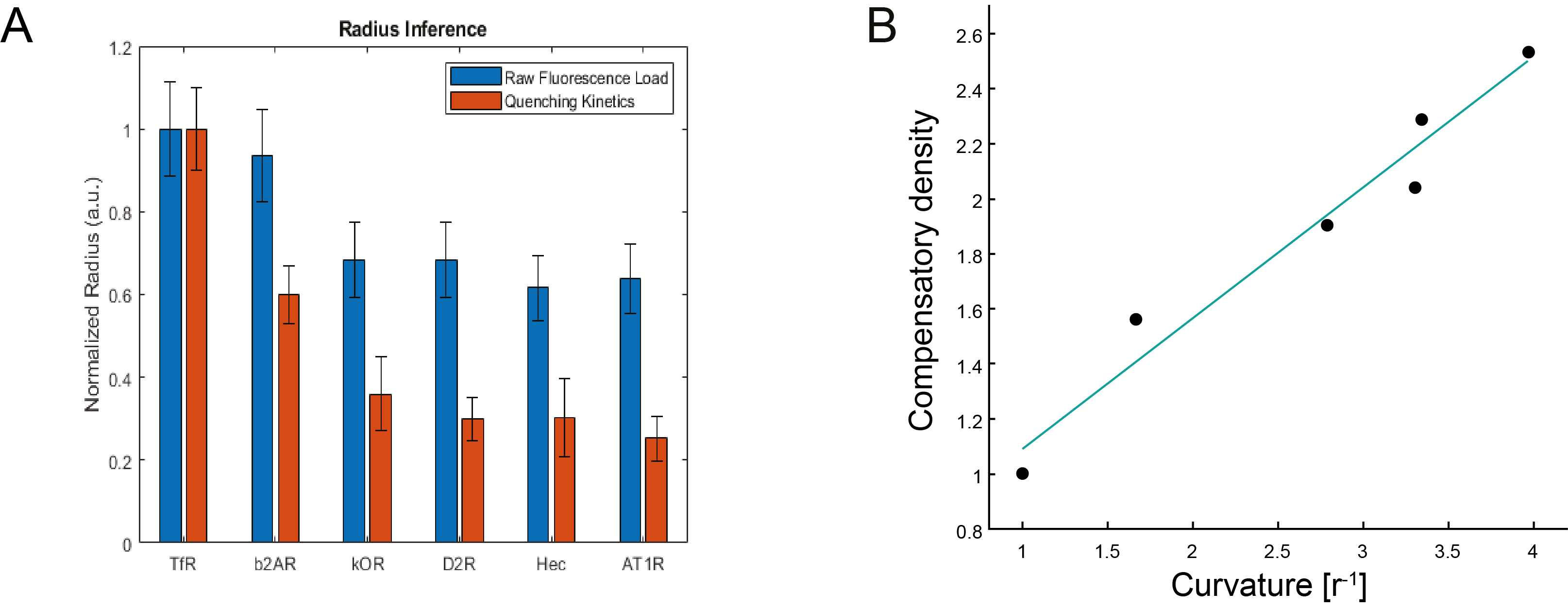

Figure S16. Higher density of cargo in smaller endocytic events identified using ppH-TIRFM

**(A)** Histograms illustrating the calculated average radii for spherical vesicular carriers associated with each of the six SEP-TacAH-constructs. Averages are either calculated directly from the observed fluorescence at time of scission (t=0, $\boldsymbol{F}\boldsymbol{\propto}\boldsymbol{r}^{\boldsymbol{2}}$, blue bars) or derived from the observed acidification kinetics (red bars). Mean values are normalized to TfR-SEP construct values, error bars represented as SEM.

**(B)** Discrepancy between the radii derived from absolute values (which is density-dependent) and the acidification kinetics (which is not density-dependent) is converted into relative measures of density increases and plotted against the calculated curvature of the carriers (here derived from acidification kinetics). The density of construct in the individual carriers increases linearly with curvature (r^2^ = 0.95, p = 0.00068).

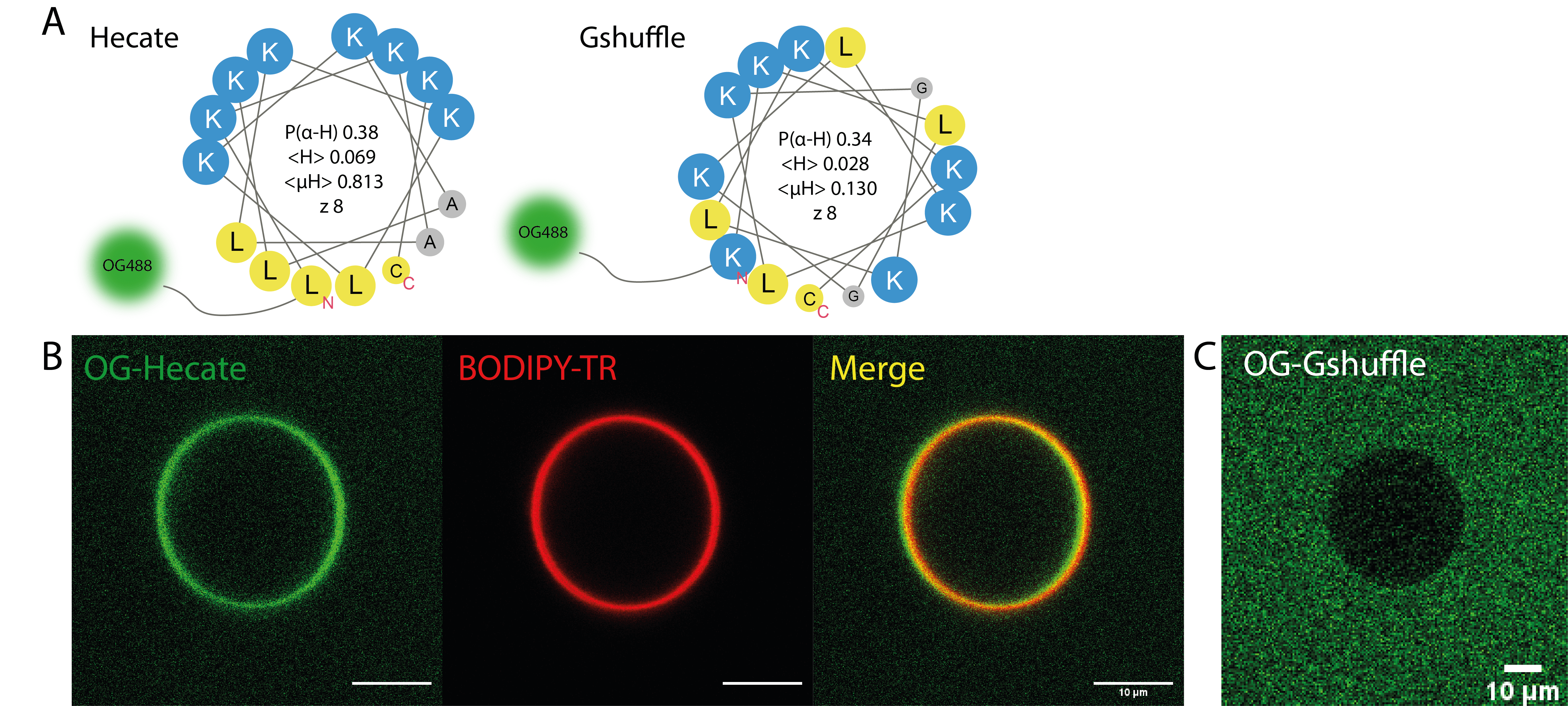

Figure S17. Binding of fluorescently labelled peptides to GUVs.

**(A)** Schematic illustration of Oregon Green 488 (OG488) labelled peptide sequences corresponding to a 15 amino acid stretch of Hecate AH (left) and the non-amphipathic Gshuffle (right, Hecate sequence with G for A substitution and randomized positions).

**(B)** Representative images of OG488 labeled Hecate AH sequence binding to BODIPY-Texas Red (TR) labeled Giant Unilamellar Vesicles (GUVs). Scale bar indicates 10 µm. c Representative image of bulk (1 µM) OG488 labelled Gshuffle in solution mixed with unlabeled GUVs showing no detectable binding of peptide to GUV. Scale bar indicates 10 µm.

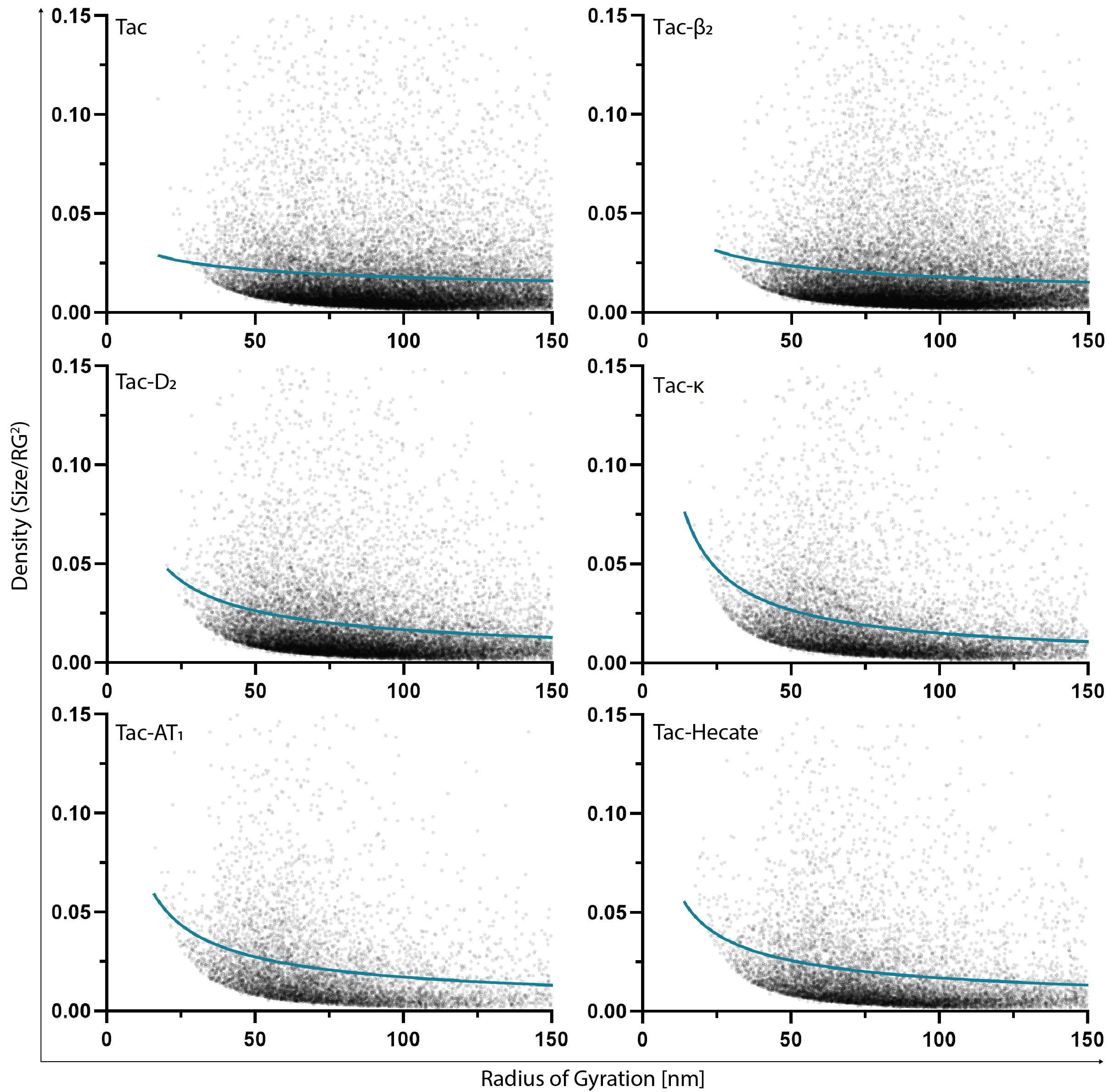
Figure S18. Increased frequency of small, high-density clusters for Tac-AH constructs with high MIP observed by 3D-dSTORM.

Non-linear regression analysis of cargo density vs size of individual clusters below 300nm diameter identified in 3D-STORM images of FLAG-TacAH constructs using the first element of a power series yielding f(x) = ax^b^.

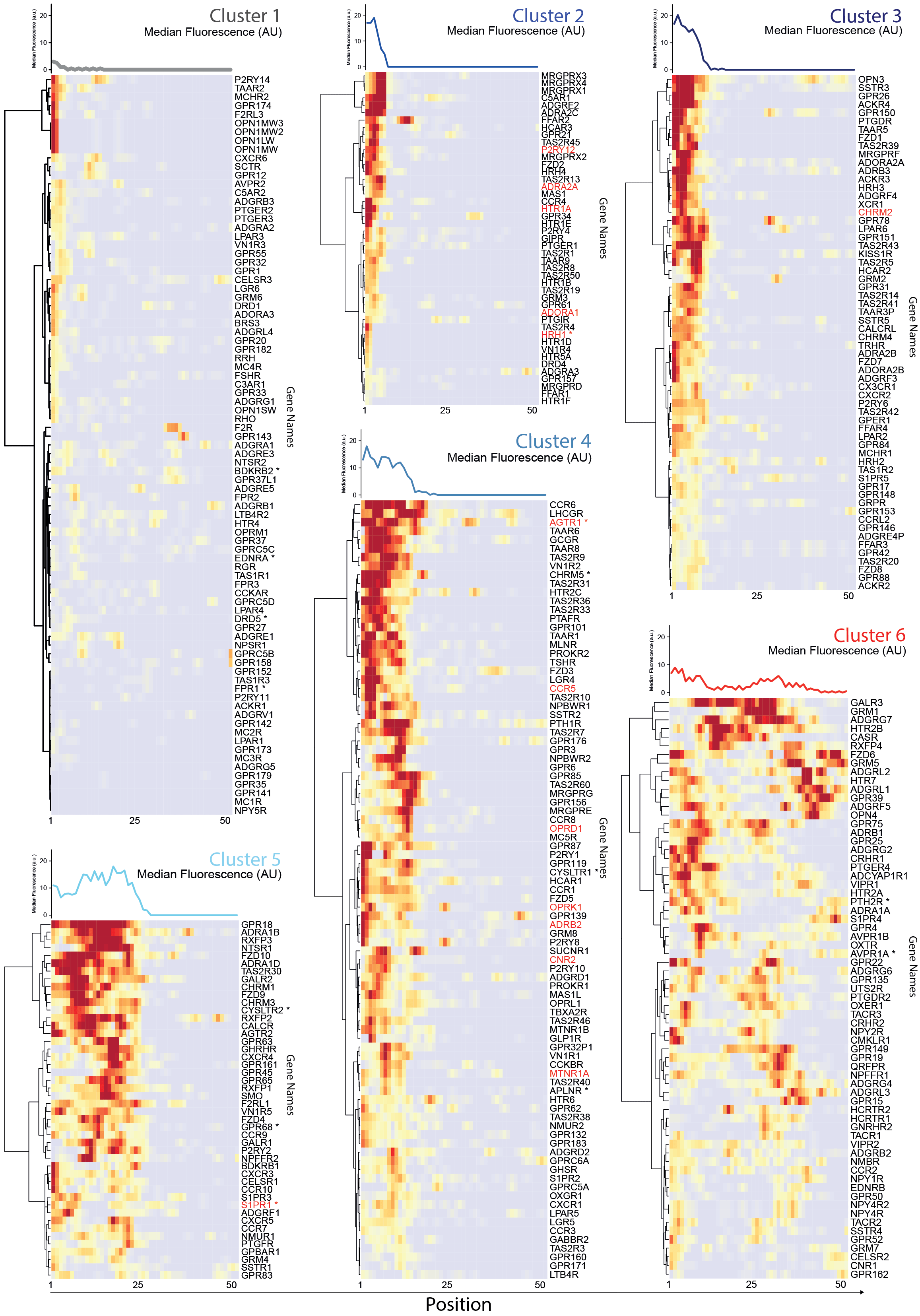
Figure S19. Membrane binding profiles of all human GPCR C-termini.

Membrane binding profiles of all human GPCR C-termini at 2 base resolution divided into 6 clusters identified by UMAP analysis (Figure 4D). (Upper) Median fluorescence at given position of all the C-termini within that cluster. (Lower) Hierarchical clustering of receptors identified in the individual clusters with the corresponding membrane binding profile and UniProt Gene Name listed ^47^. Receptors used in Figure 4F highlighted in red and receptors reported as mechanosensory highlighted with an asterisk.

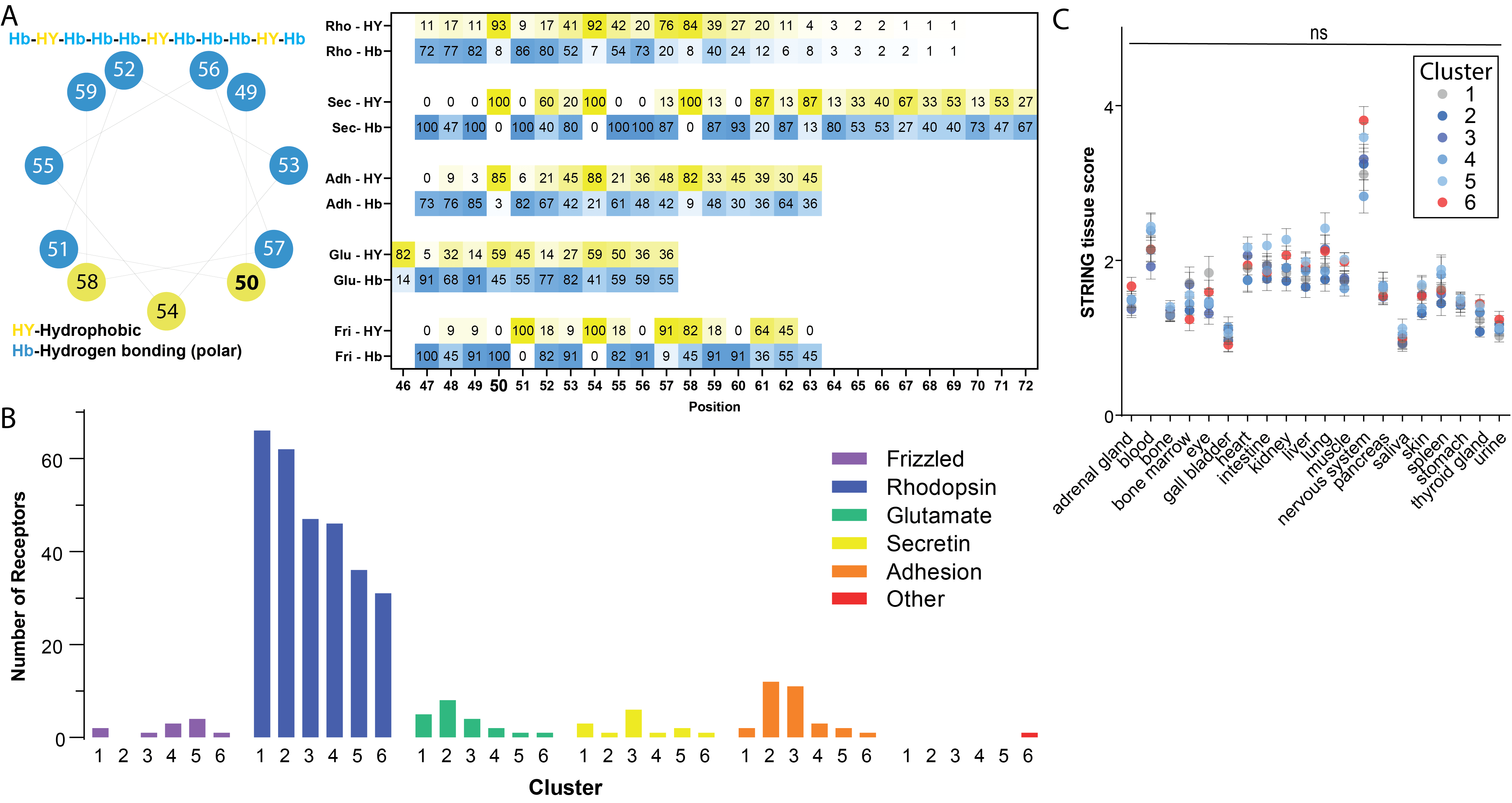

Figure S20. Membrane binding properties of H8 across GPCR families.

**(A)** (Left) Helical wheel representation showing consensus amphipathic distribution of conserved amino acid properties found across GPCR families. (Right) Heatmap of aligned H8 sequences indicating conservation of hydrophobic and polar residues in GPCRs by family in percentage of receptors as annotated on GPCRdb ^11^.

**(B)** Frequency representation of GPCRs of individual GRAFS families across the 6 identified clusters in Figure 4.

**(C)** Tissue expression scoring according to STRING database for receptors belonging to clusters 1-6. Lines indicate average score +/- SEM. One-Way ANOVA analysis identified no significant difference**.**

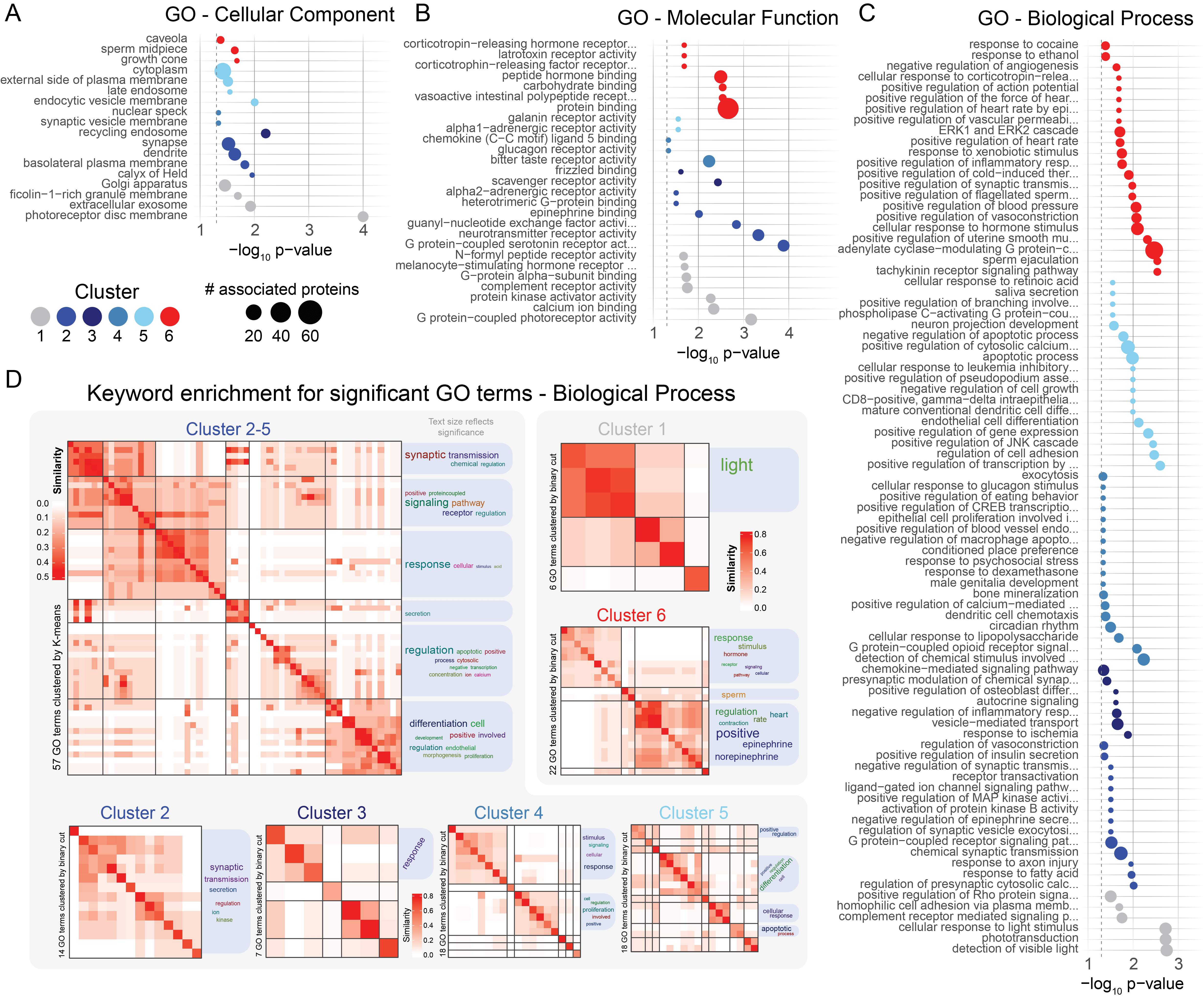

Figure S21. Enrichment analysis reveals distinct localization and function between GPCR clusters.

Conditional enrichment analysis of all non-olfactory GPCRs for the three Gene Ontology (GO) term families for each cluster of membrane binding profiles (Figure S19).

**(A)** Results for Cellular Component (CC)

**(B)** Results for Molecular Function (MF)

**(C)** Results for Biological Process (BP)

**(D)** To assess semantic similarity of BP terms in (C) we used Resnik GO term similarity scores identifying simplified terms for clusters 1-6 alongside pooled results for clusters 2-5 (bona fide H8 receptors).

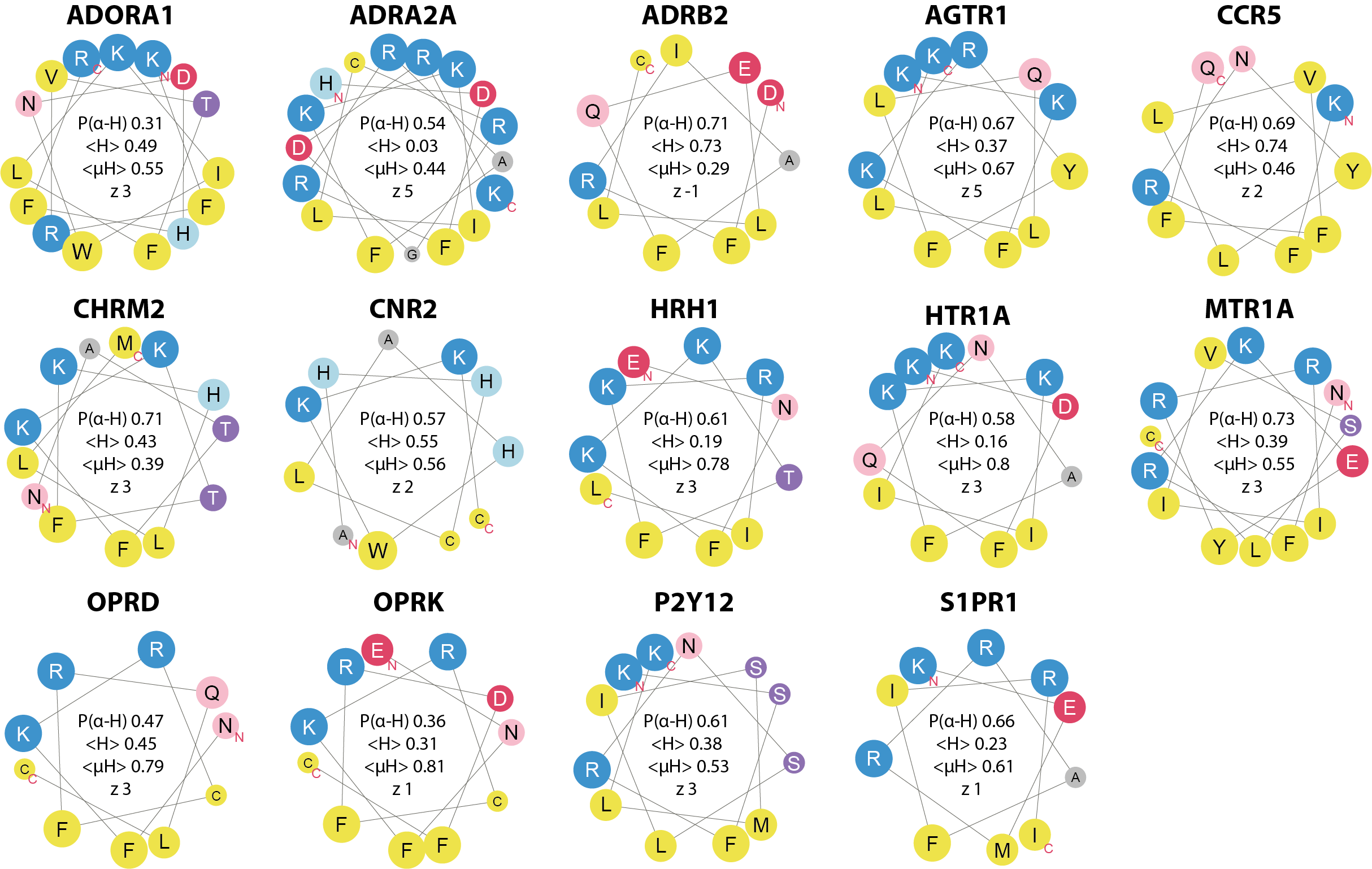

Figure S22. Helical wheel representation of amphipathic helices and their biochemical parameters in GPCRs of clusters 2-5.

Helical wheel representations obtained using Heliquest ^43^ of the AHs in Figure 4F in alphabetical order. The chemical nature of amino acid side chains is color coded (Blue, positively charged; Red, negatively charged; Green, constrained; Grey, small non-polar; light red/purple, hydrophilic; Yellow, hydrophobic. Size of circles indicate the size of the side chain. P(a-H): a-helical propensity (obtained by NetsurfP); <H>: mean hydrophobicity; <µH>: hydrophobic moment; z: net charge.

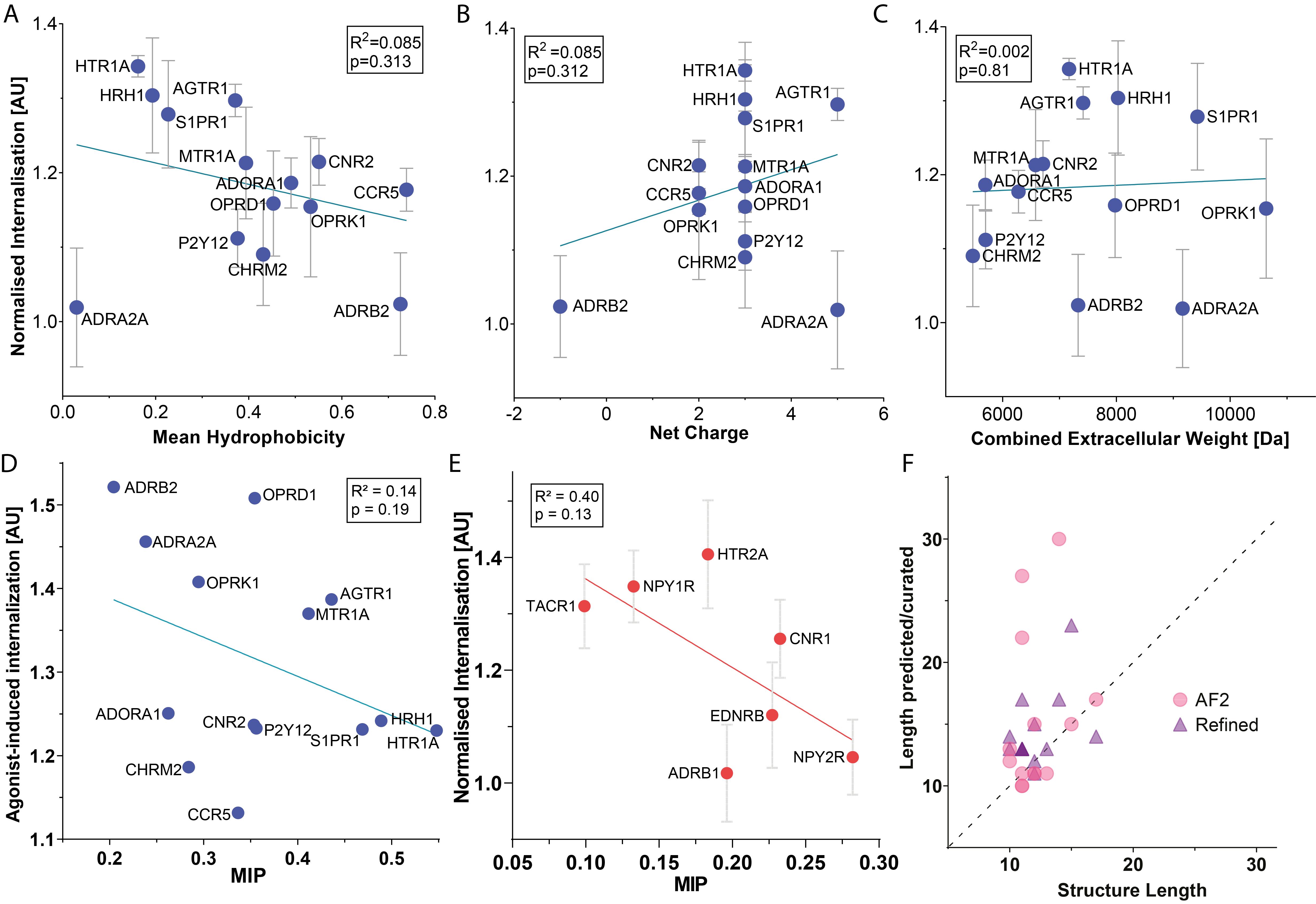

Figure S23. Explanatory value of other physical parameters identified in H8 and extracellular domains of GPCRs in cluster 2-5.

**(A-C)** Constitutive internalization of GPCRs of clusters 2-5 from Figure 4 as determined by flow cytometry in ^39^ as a function of mean hydrophobicity (A), and net charge (B) of H8, as well as the combined weight of extracellular residues (C).

**(D)** Agonist-induced internalization of receptors of clusters 2-5 from Figure 4 as determined by flow cytometry in ^39^ as a function of MIP.

**(E)** Constitutive internalization of receptors of cluster 6 as a function of MIP. f Correlation between the predicted length of H8 by AlphaFold2 ^44^(magenta) and refined structures by GPCRdb ^11^(purple) versus the observed length of H8 manually identified in structures.

**(A-C+E)** Mean ± SEM, N=3.

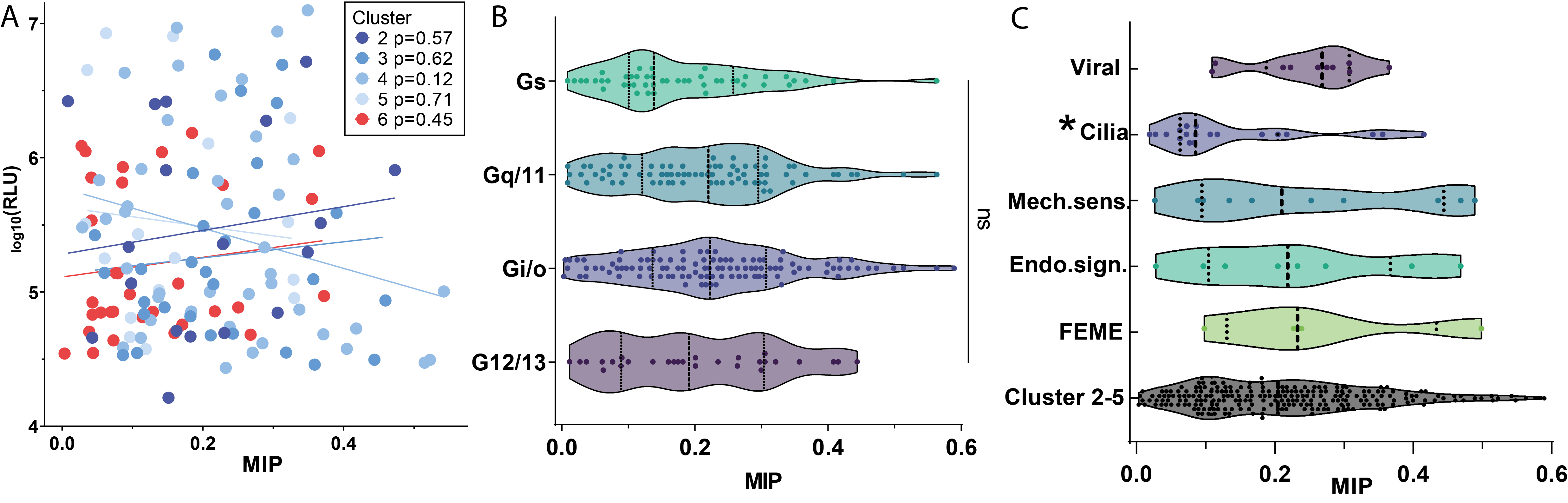

Figure S24. MIP of H8 sequences does not correlate with β-arrestin signalling or G-protein coupling, but anti-correlates with ciliary location. Change order

**(A)** Receptor signaling measured in relative luminescence units, as reported in ^48^ of receptors in clusters 2-6 as a function of MIP.

**(B)** The distribution of MIP of H8 in receptors of clusters 2-5 by reported G protein coupling ^49^. Lines indicate quantiles. One-Way ANOVA analysis identified no significant difference.

**(C)** The distribution of MIP values of human receptors of clusters 2-6 reported in relation to FEME ^30^, Endosomal signalling (Endo.sign.), Mechanosensation (Mech.sens.) ^41^ and Cilia as well as for MIP values of H8 sequences identified in viral GPCRs listed in ^40^ using NetSurfP3.0 ^23^ alongside the distribution of MIP values for combined human H8 sequences of clusters 2-5. Lines indicate quantiles. Asterisk indicates p<0.05 using Kruskal-Wallis multiple comparisons test versus Cluster 2-5.

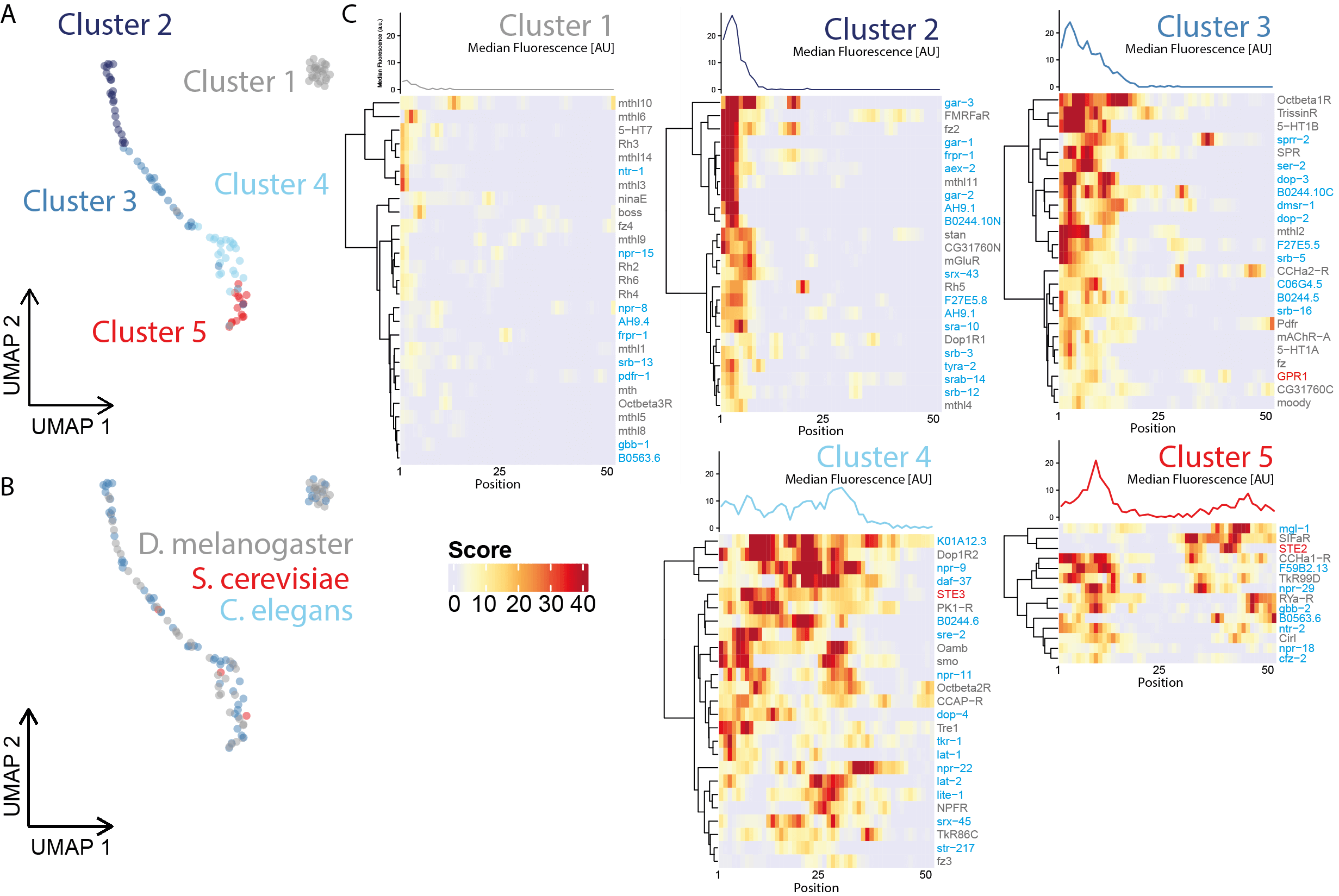

Figure S25. Selective drive for evolutionary conservation of MIP across distant species.

**(A)** UMAP-based clustering of combined *D. melanogaster*, *C. elegans* and *S. cerevisiae* GPCR C-termini based on position and strength of liposome binding motifs.

(**B**) Distribution of species in A.

**(C)** Heatmaps of individual GPCR C-terminal liposome binding profiles within the 5 clusters (lower) and profile plots of their median fluorescence (upper). Species of individual receptors highlighted by text color (*D. melanogaster*: grey, *C. elegans*: cyan and *S. cerevisiae*: red)*.*

| **Construct** | **Insert** |
| --- | --- |
| Alcohol dehydrogenase (Alc.DH) | INEGFDLLRSG |
| ALPS | FLNNAMSSLYSGWSSFTTGASKFAS |
| AT_1_ | GFLGKKFKKYFLQLLKYIPP |
| β_2_ | SPDFRIAFQELLCLRRSS |
| Rho | NKQFRNCMVTTLCCGKN |
| Citrate synthetase (Cit.Syn) | LSFAAAMNGLA |
| D_1_ | AFNADFRKAFSTLLGCYRLCPA |
| D_2_ | FNIEFRKAFLKILHC |
| Endophilin | MSVAGLKKQFHKATQKVSEKV |
| Epsin-1 | STSSLRRQMKNIVHNYSS |
| Hecate | FALALKALKKALKKLKKALKKAL |
| κ | AFLDENFKRCFRDFCFPLKM |
| Melittin | GIGAVLKVLTTGLPALISWIKRKRQQ |
| NK_1_ | NDRFRLGFKHAFRCCPFI |
| SAR | SFIFDWIYSGFSSVLQFLGLYKKTG |
| SpoVM | KFYTIKLPKFLGGIVRAMLGSFRKD |
| TA_1_ | FFYPWFRKALKMMLFGKI |
| TA_6_ | LFYPWFRKAIKVIVTGQV |
| Troponin C | KEDAKGKSEEE |

Table S1. Inserted amino acid sequences of H8 in TacAH constructs. Listed FLAG-TacAH constructs with the corresponding inserted amino acid sequences (one-letter code) harbouring AHs.

| **Name** | **Species** | **UniProt** | **MIP** |
| --- | --- | --- | --- |
| BILF1 | EBV | P03208 | 0,187071 |
| US28 | HCMV | P69332 | 0,320226 |
| US27 | HCMV | P09703 | 0,209586 |
| UL33 | HCMV | P16849 | 0,222506 |
| UL78 | HCMV | F5HET1 | 0,365568 |
| U12 | HHV6 | P52380 | 0,283792 |
| U51 | HHV6 | P52382 | 0,307771 |
| U12 | HHV7 | P52381 | 0,113649 |
| U51 | HHV7 | P52383 | 0,10925 |
| ORF74 | KSHV(HHV8) | Q98146 | 0,262279 |

Table S2. MIP values identified in virally encoded GPCRs. MIP values of putative H8 sequences in virally encoded GPCRs.

### Supplementary Materials

The following supplementary information is available for this paper.

Methods

Mathematical model “Acidification Kinetics Function”

### Methods

### Molecular Biology

Coding sequences for the amphipathic helices AHs (outlined below) were synthesized and cloned in frame into a previously described pcDNA3 Flag Tac construct (Madsen, Eriksen et al., 2008) using HindIII and XbaI sites. The amino acid sequence of Tac is largely equivalent to the Interleukin-2 receptor subunit alpha, with a FLAG-tag (DYKDDDDK) sequence inserted downstream of the signal peptide sequence and a trailing QASS sequence at the C-terminus (see Figure S4). The C-terminal serine is replaced with the following inserts containing putative helical sequences listed in Table S1. Full length ATR was a kind gift from Prof. Thue W. Schwartz (Metabolic Receptology, University of Copenhagen) and b2AR was kindly distributed by Prof. Mark von Zastrow (Dep. Of Psychiatry, UCSF, USA). Di-glutamate mutations were introduced following Quick Change® method (Stratagene, La Jolla, CA, USA).

Super Ecliptic pHluorin (SEP) constructs were produced using PCR amplification and subsequently inserted by restriction enzyme cloning. Chimeric constructs were produced using QuickChange method (Stratagene, La Jolla, CA, USA). PRESTO-Tango library was acquired from Addgene (Kit #1000000068). FLAG-TacAH-BioID2 constructs were prepared using SLIC method ^50^, amplifying the coding region of BioID2 ^25^(Addgene plasmid #74224), and inserting it downstream of FLAG-TacAH with a five amino acid linker sequence (AAGGS). All constructs were verified by DNA sequencing.

### Cell Cultures

Human Embryonic Cells (HEK293), BSC-1 cells and HeLa cells were cultivated in growth media (Dulbecco’s Modified Eagle’s Medium 1965 (DMEM) supplemented with 10% Fetal Bovine Serum (FBS) and 1% Penicillin/Streptomycin) and maintained at 37°C and 5% CO_2_ levels. Cells were sub-cultured every 3-4 days for maintenance to a maximum passage of 35 and never allowed to reach a confluency above 95%.

### Transient transfection for internalization assays

A day prior to transfection, cells were seeded on poly-L-ornithine treated coverslips in TPP polystyrene 6-well plates (Sigma-Aldrich) at a density of 3.5x10^5^ cells/well, unless otherwise stated. Transfection was performed using Lipofectamin 2000 and Opti-MEM (ThermoFisher). The manufacturer´s protocol was followed with a 3:1 lipofectamin to DNA ratio. A total of 0.5-1.5 µg/well was used and cells were incubated with transfection mix in Opti-MEM for 5 hours. Opti-MEM was replaced with growth media, and cells were used for experiments 48 hours after transfection.

### Antibody conjugation

Primary anti-FLAG M1 antibodies used for feeding assay were fluorescently labelled by NHS-ester fluorophore conjugation. A 90 µL antibody solution (1mg/mL) was added to 10 µL NaHCO3 (1M) and 5 µL NHS-ester fluorophore (Alexa Fluor®488 or Alexa Fluor® 647) (1mg/mL). The mixture was incubated for 1 hour at 25°C and antibodies were isolated using illustraTM NAP-5 columns (GE Healthcare Life SciencesTM), resulting in an average labeling rate of 4 and 8 dye molecules per antibody for 488 and 647 respectively, as measured on a Thermo ScientificTM NanoDrop 2000c.

### Antibody Feeding Assay for Confocal Imaging

On day of experiment cells were incubated with primary anti-FLAG M1 antibody (1 µg/mL in Opti-MEM, for 1 hour at 4°C). Cells were then exposed to pre-warmed (37°C) growth media containing 25 µM Monensin (Sigma-Aldrich) and kept at 37°C for 30 minutes. Cells were fixed in a 4% PFA solution in PBS for 10 minutes and subsequently washed with PBS. Next, cells were passivated using blocking buffer (5% GS in PBS) and incubated for 20 minutes at 25°C. Surface-bound M1 antibody was labelled using Alexa Fluor 488 conjugated goat anti-mouse secondary antibody (1:200 in 5% GS, 1 hour, 25°C). Cells were fixed again in 4% PFA in PBS for 10 minutes at 4°C, washed in PBS and permeabilized with 0.2% saponin and 5% GS in PBS (30 minutes at 25°C). Next, cells were incubated with Alexa Fluor 568 conjugated secondary goat anti-mouse antibody (1:500, 5% GS, 30 minutes 25°C). Finally, cells were washed three times with PBS and coverslips were mounted using DAPI Fluoromount-G (SouthernBiotech).

Selective interference of actin polymerization, macropinocytosis and dynamin constriction was performed by addition of 200nM Latrunculin A (Thermo Fischer), 1µM 5-[N-ethyl-N-isopropyl] amiloride (EIPA), and 10µM Dyngo-4A (Abcam) during 30-minute internalization period respectively.

Cells were imaged as z-stacks with increments of ~700 nm using either a Zeiss Confocor 2 confocal microscope (LSM 510) or Zeiss Axoimager M2 confocal microscope (LSM700) with a Plan-Apochromat oil immersion objective with a 63x magnification and NA of 1.4 (Carl Zeiss, Oberkochen, Germany). Fixed samples were imaged in frames of 1024x1024 resolution, 8bit, in line-scanning mode with 2 times averaging. Alexa Fluor® 568 dye was excited using 543/555-nm laser light from a helium-neon (HeNe) laser or diode source, and emitted fluorescent light was filtered using a 585-nm long pass filter or none. Alexa Fluor 488 dye was excited using a 488-nm laser line from an argon-krypton laser or diode, and the emitted light was filtered using a long pass 505–530-nm barrier filter or none. Channels were imaged separately. Image processing was performed using FIJI ^51^(National Institutes of Health). Region of interests (ROIs) were selected to accommodate entire cells throughout the stack, as inspected through green channel only (up to a max of 4 ROIs per image). The threshold was set to exclude background noise, kept constant throughout independent experiments, and all images were masked. Intensities were quantified independently for each stack in each channel and saved as comma separated files. A MatLab script was generated to automatically import data and calculate overall intensity ratios for each cell. Individual ratios for each cell were log_10_-transformed prior to averaging within each construct and normalization to Tac. Averages and standard error of mean was determined within each experiment on basis of a minimum of 10 individual cells. Linear regression analysis was performed using Prism (GraphPad Software, Inc.). All ROIs were saved for later inspection.

### Antibody Feeding Assay for Flow Cytometry

Cells were seeded in TPP polystyrene 6-well plates (Sigma-Aldrich), transfected as described above, and kept in plates until experiment. On day of experiment cells were incubated with Alexa Fluor 647 conjugated primary anti-FLAG M1 antibody (1 µg/mL, 1 hour at 4°C). Cells were then exposed to pre-warmed (37°C) growth media containing 25 µM Monensin and kept at 37°C for 30 minutes. For experiments with chimeric constructs, internalization media additionally included 10 µM isoproterenol (ISO) and 10 µM Angiotensin II (ANGII) for β2AR and AT1R respectively. Cell surfaces were stripped for antibody by addition of cold strip-buffer (0.5M NaCl, 0.2 acetic acid) and incubation for 5min followed by a wash in PBS and cold-labeling with Alexa Fluor 488 conjugated primary anti-FLAG M1 antibody (1 µg/mL, 1 hour at 4°C). Cells were collected using cold PBS and transferred to Eppendorf-tubes, spun down at 2000xg for 10 minutes and resuspended in 4% PFA prior to flow cytometry acquisition.

Samples were run on a FACSCalibur flow cytometer (BD Biosciences) and analyzed with FlowLogic software (Inivai Technologies). Briefly, empty pcDNA3 transfected cells without fluorophores were used to gate cells from debris through a combination of SSC-H and FCS-H signals (Figure S3). Unspecific fluorescence gating was set to include a maximum of 1% of events from untransfected cells otherwise treated identical to samples. Only events corresponding to gated cells with specific fluorescence signal were used for further analysis. Alexa Fluor 488 was excited using a 488 nm Argon laser and the emitted fluorescence filtered through a 530/30 BP filter and collected by a PMT (FL-1). Alexa Fluor 647 was excited using a 635 nm diode laser and the emitted fluorescence filtered through a 661/16 BP filter and collected by a PMT (FL-4). Extracted intensities in FL-1 and FL-4 channels were used for quantification. Briefly, a two-dimensional dot plot of the intensities of surface bound (FL-1) and internalized fluorophores (FL-4) for each investigated cell of each construct was analyzed using a python-based random sample consensus (RANSAC) algorithm from scikit-learn ^52^, giving an outlier optimized linear regression slope with minimized influence of outliers. The resulting slopes were normalized to the reference construct Tac yielding relative internalization.

The preparations for timeline experiments were done by incubating transfected cells with Alexa Fluor 647 conjugated primary anti-FLAG M1 antibody (1 µg/mL, 1 hour at 4°C) and then incubated at 37°C for indicated time-periods with pre-warmed (37°C) growth media containing 25 µM Monensin. Surface-bound antibodies were stripped by 5-minute incubation with cold strip-buffer (0.5M NaCl, 0.2 acetic acid). Cells were fixed and run on FACSCalibur flow cytometer for acquisition. Internalization was quantified as mean Alexa Fluor 647 signal subtracted by the mean signal obtained from samples at timepoint 0 in percentage of the same signal obtained for samples that were only cold labelled.

### Transferrin Uptake by Flow Cytometry

Cells were seeded in TPP polystyrene 6-well plates (Sigma-Aldrich), transfected as described above, and kept in plates until experiment. On day of experiment, cells were incubated with Alexa Fluor 488 conjugated primary anti-FLAG M1 antibody (1 µg/mL, 1 hour at 4°C) and Alexa Fluor 647 conjugated transferrin (25µg/ml) (Thermo Fischer). Cells were then exposed to pre-warmed (37°C) growth media containing 25 µM Monensin and kept at 37°C for 12 minutes. All cells were collected in cold PBS and transferred to Eppendorf-tubes, spun down at 2000xg for 10 minutes and resuspended in 4% PFA prior to acquisition in flow cytometer. Samples were run on a FACSCalibur flow cytometer (BD Biosciences) and analyzed with FlowLogic software (Inivai Technologies) similar to what was described for antibody feeding assays.

### Live cell imaging on pulsed pH (ppH) TIRF setup for endocytosis

Live cell imaging was performed at 37°C. Cells were perfused with HEPES buffered saline (HBS) solution (135 mM NaCl, 5 mM KCl, 0.4 mM MgCl_2_, 1.8 mM CaCl_2_, 20 mM HEPES, and 1 mM d-glucose, adjusted to pH 7.4 and 300–315 mOsmol/l). For the ppH protocol, a theta pipette ∼100-µm tip size achieved using a vertical Narishige puller (World Precision Instruments) was placed in proximity of the recorded cell. One outlet contained HBS (pH 7.4; see previous paragraph for recipe) and the other contained a solution where HEPES was replaced with MES (MES buffered saline solution) and buffered at pH 5.5. Solution flow was alternated every 2 s using electrovalves (Lee Company) switching in synchrony with image acquisition. TIRF imaging was performed on an inverted microscope (IX71; Olympus) equipped with an Apochromat N oil 60× objective (NA 1.49), a 1.6× magnifying lens, and an electron multiplying charge coupled device camera (QuantEM:512SC; Roper Scientific). Samples were illuminated by a 473-nm laser (Cobolt) for SEP imaging, as well as by a coaligned 561-nm laser for red FP imaging. Emitted fluorescence was filtered using the following filters (Chroma Technology Corp.): 620/60 m for mCherry imaging, and 525/50 m for SEP/GFP fluorescence. Simultaneous dual color imaging was achieved using a DualView beam splitter (Roper Scientific) and dual cameras. To correct for x/y distortions between the two channels, images of fluorescently labeled beads (Tetraspeck, 0.2 µm; Invitrogen) were taken before each experiment and used to align the two channels (see Quantification of fluorescence intensities). The camera was controlled by MetaVue7.1 (Roper Scientific).

### Detection and Quantification of endocytic events in ppH-TIRF

Detection and subsequent quantification of fluorescent protein assemblies was done as previously described ^35^. Briefly, spots were subjected to segmentation based on wavelet transform Multidimensional Image Analysis (MIA). The objects detected were then tracked using a simulated annealing algorithm to identify endocytic events. The output of this tracking was a series of coordinates corresponding to the center of mass of the objects, with unique identifiers (event numbers). All the observed fluorescent assemblies in the TfR5.5 movies were screened to identify genuine endocytic events using routines programmed in Matlab 7.4 (Mathworks). To qualify as bona fide events each candidate event required a TfR5.5 vesicle (i) that persisted for at least three frames (i.e., 8 s) following appearance, (ii) that appeared at least 20 frames after the start of the movie, or 20 frames before its end, to ensure quantification of signals for 80 s before and after the vesicle's appearance, (iii) that appeared and remained at more than seven pixels (0.7 µm) from the edge of the image, to ensure proper quantification (see below), (iv) that appeared de novo, and was not produced by the fusion of two objects or the dissociation of an object into two, (v) that overlapped, on appearance, with a pre-existing cluster detected in the segmented TfR7.4 movie, (vi) whose fluorescence was bigger than a defined SNR of 5 wherein SNR = (F_0_−av)/std, where F_0_ is the fluorescence at time 0, and av and std are the average fluorescence and standard deviation, respectively, in the five frames before vesicle appearance, and (vii) that was close to maximal fluorescence at the time of appearance. We calculated the slope of the fluorescence change in the first three frames of vesicle appearance and discarded the events where this slope was greater than 0.1, which corresponds to a 10% increase in fluorescence.

### Characterization of trace profiles from ppH-TIRF

While the spatial and temporal extension of the fluorescent traces in the TfR5.5-channel is defined directly by the formation and acidification of single vesicle carriers, the fluorescent traces describing clusters in TfR7.4, CLC5.5 and CLC7.4 can encompass and span several events. The resultant profile characteristics therefore contain important information of the underlying mechanistic of each construct. Hence, we initially assessed the CLC-profiles and TfR7.4-profiles connected to each TfR5.5-event by normalizing each of the traces individually, smoothening the traces by a loess model to remove noise, and ultimately generating collected raster-plots for each construct (Figure S13, Figure 2J, Figure 3A).

In the case of TfR7.4-profiles, we used these collections to estimate the overall recruitment rates and collaborative nature of the clusters comparatively by performing logistic fitting of the profiles just before scission (t = [-20:0]) to an adapted version of the Langmuir-Hill-equation: θ = 1/(1+(T50/t))^H, where θ is the fraction of total cluster-size estimated at the time of scission, t is the time from -20 to time of scission, T50 the time where cluster is grown to half size, and H is the Hills-coefficient. These fits allowed to extract the Hills-coefficient as a measure for the collaborative recruitment of constructs during accumulation.

As the collected CLC-profiles (CLC5.5/7.4 interleaved) contained information of the overall activity surrounding the individual events (Figure S13, Figure S14), we performed a principal components analysis (PCA) on all the normalized traces pooled together to evaluate the principal components giving rise to the observed profiles. Together, the first four principal components contained 63.9% of the total variance and was used to sort and the individual raster-plots and evaluate the governing mechanics pertaining to each construct

### High frequency live cell imaging for exocytosis studies

Live cell imaging was performed at 37°C. Transfected BSC-1 cells were grown in LabTek2® chambers and imaged using a Nikon Eclipse Ti-E epifluorescence/TIRF microscope (Nikon) equipped with a 405 nm laser (coherent) and a triple line 405/488/561 beamsplitter. Cells were kept in growth medium prior to imaging and media was substituted for temperature-equilibrated PBS buffer to reduce background fluorescence during imaging.

The emission signal from pHluorin-constructs and CLC-mCherry was split by a 509nm beamsplitter (AHF analysentechnik AG), the pHluorin-signal was filtered using a 466/40nm BP filter, and the CLC-mCherry signal by a LP filter. Light was directed to a dual view adaptor collected by two EM-CCD cameras (iXon3 897, Andor).

### SPOT assay

Peptide arrays were prepared as described in ^53^ with sequences identical to the Flag-TacAH-constructs inserts. We developed and optimized a protocol for liposome binding ^54^. In brief, plates were coated with 5% BSA prior to binding, washed and exposed to a 0.01mg/mL dilution of BBV liposomes containing 0.5% biotin. Liposomes were made by dissolving Folch bovine brain lipids in chloroform together with 0.5% DOPE-biotin, evaporating samples with N_2_ and rehydrating in buffer to a final concentration of 1 mg/mL. After 8 freeze-thaw cycles, the liposomes were sonicated briefly at low intensity, subjected to another freeze-thaw cycle, and stored at 4°C until use. Storage time was always kept below 36 hours. The arrays were incubated with liposomes for 30 minutes at 25°C, washed three times in PBS and exposed to 5 mL of a 0.01mg/mL avidin-HRP solution to cover the entire array. Arrays were allowed to equilibrate for 30min at room temperature, washed three times in PBS and moved to an Alpha Innotech MultiImage^TM^ Light Cabinet, where it was covered with ECL substrate and imaged. Total intensities were quantified with ImageJ.

### Data Retrieval of Membrane Proximal C-terminal regions in GPCRs

We extracted the complete set of Homo sapiens (taxid: 9606) gene products from the UniProtKB Swiss-Prot database with the Gene Ontology (GO) annotation “part of integral component of membrane” (GO:0016021) and “G protein-coupled receptor activity” (GO:0004930) from the EBI GO Annotation database (GOA) (<https://www.ebi.ac.uk/GOA>, 2020-08-17). This resulted in a dataset containing 807 distinct gene products. Gene products annotated with “olfactory receptor activity” (GO:0004984) were removed resulting in 410 gene products. The resulting gene products were exported as a Gene Product Association Data file. The UniProtKB accession numbers were used to extract their respective topological data from the Universal Protein Resource Knowledgebase (UniProtKB) (<https://www.uniprot.org/>). The following extraction, sorting and analysis of data was performed using the programming language R (version 4.0.2) in RStudio (version 1.4.1013). Sequences were analyzed for annotated regions of secondary structure and intermembrane regions. The identified regions were removed from the sequences and only the regions residing in proximity to transmembrane domains were kept in the dataset. We added all C-termini from S. Cerevisiae, C. elegans and D. Melanogaster G-Protein Coupled Receptors (GPCR) to the dataset (140 GPCRs as obtained from the GO resource), enabling an evolutionary conservation analysis. The resulting sequences were split into 16mer cassettes overlapping by 14aa (2aa resolution), which resulted in 15198 unique peptides.

### High-Density Peptide Array Synthesis and Scanning

Synthesis was performed by Schafer-N ApS, Lersö Park Allé 42, DK-2100 Copenhagen as described by Schafer-Nielsen and coworkers ^55^. All unique *in-silico* extracted 14aa overlapping 16mer GPCR-specific peptides were randomly split into 6 evenly sized groups and synthesized onto 6 equally sized sectors on the array. Each peptide was synthesized in a 20x20µm field, separated from other fields by 10µm from all sides. As a positive control for liposome binding Hecate (LKKALKKLKKALKKAL) was synthesized in the 17 peptide fields encompassing each of the corners of the 6 sectors on the array. All peptides were synthesized onto the array with a short C-terminal linker KSGG. 510 negative controls, fields not containing peptides, were spread randomly across the array. The array was blocked using 5 % BSA in PBS with 2 mM DTT for 1h on a shaker at 120 rpm and then washed 3 times in PBS for 20 min. The array was incubated with DiD-labelled liposomes diluted 1:100 in PBS (pH 7.4) for 1h at RT. Subsequently the array was washed twice in PBS for 5 min and an additional time for 20 min. Following 3x 30sec washes with 0.1% Tween-20 in PBS the array was dried by centrifugation (30sec) and scanned at 1µm resolution with an InnoScan 900 laser-scanner (INNOPSYS) with an excitation wavelength of 635nm and gain setting 6 with the high-power setting to obtain the image used for this analysis.

### Conditional GO term enrichment analysis

All non-olfactory Human GPCRs represented were selected from the array data by exclusion of Olfactory GPCRs annotated with GO term GO:0004984 (olfactory receptor activity). GO terms were imported via pantherdb.org using rbioapi package in R (datetime: 23/09/2024 18:08). Conditional enrichment was run for each cluster using the topGO package in R algorithm='weight01', test='fisher'. The Resnik GO term similarity was calculated in each cluster using the simplifyEnrichment package in R with the background population set to GPCRs manually. Similarity matrix was clustered using a binary cut or kmeans (see file name). Keyword enrichment was performed for each cluster (or cluster 2-5 combined)(Fisher test vs GPCR GO terms). GO term similarity matrix was plotted with significantly enriched cluster associated keywords for clusters with n>4 terms

### Liposome Preparation

Liposomes were generated using a heterogenous lipid extract from bovine brain (Type I, Folch fraction I, Sigma-Aldrich). 0.5mg lipid extract was dissolved in 1mL of chloroform, before addition of 2% DiD (Invitrogen™) (w/w). Lipids were dried with N_2_-gas while rotating, creating a thin lipid film, followed by vacuum dehydration for 12 hours to remove residual chloroform. Lipids were rehydrated in a sterile filtered phosphate buffered saline (PBS) (pH 7.4) to a final concentration of 1 mg/mL. The rehydrated lipids were put through 8 freeze-thaw cycles using liquid nitrogen and a 40-50°C water bath. The liposomes were extruded through a 19 mm polycarbonate Whatman^TM^ Nuclepore^TM^ Track-Etched membrane with a pore size of 1 µm using a LiposoFast liposome extruder (Avestin). The liposomes were extruded 18 times before being diluted 1:1000 in sterile PBS.

### 3D-dSTORM – Sample preparation

BSC-1 cells were seeded on coverslips in TPP polystyrene 6-well plates (Sigma-Aldrich), transfected as described above, and kept in plates until experiment. On day of experiment cells were fixed using a 4% PFA solution (10 minutes at 4°C), washed in PBS (3x) and passivated using blocking buffer (1% BSA and 5% goat serum solution in PBS) for 20 minutes prior to incubation with anti-FLAG M1 antibody (1 µg/mL in 1% BSA and 5% goat serum solution for 1 hour at 25°C, ThermoFisher). Cells were washed in PBS (3x), fixed in a 4% PFA solution (10 minutes at 4°C), washed again (3x), before being passivated for 20 minutes and incubated with secondary Alexa Fluor 647 conjugated anti-mouse antibody (1 µg/mL in 1% BSA and 5% goat serum solution for 45 minutes at 25°C, ThermoFisher). Finally, cells were washed in PBS (3x), fixed using 4% PFA solution (10 minutes at 4°C) and washed once in PBS containing 20 mM Glycine and 15 mM Ammonium Chloride to quench free aldehydes, followed by wash in PBS (2x). The fixed samples were then transferred to a quick release imaging chamber and imaged in a buffer containing 560 g/mL Glucose-oxidase, 34 g/mL Catalase, 2 mM cyclooctatetraene, 1% 2-Mercaptoethanol, 50 mM Tris-HCL (pH 8), 10 mM NaCl and 10% Glucose in water.

### 3D- dSTORM – Optical setup and image acquisition

Single-molecule localization imaging was performed on a Zeiss Elyra PS.1 inverse microscope. Samples were illuminated using 405 nm and 642 nm HR Diode lasers with maximum power levels of 5 mW and 150 mW respectively. The 642 nm laser was used at maximum power, while 405 nm laser was used to reactivate Alexa Fluor 647 fluorophores and power was incrementally increased by 0.1 percentage points. Light was collected in a Plan-Apochromat 63x oil objective with an effective numerical aperture of 1.4. Emitted fluorescence was collected in the same objective and further magnified using a 1.6x tube lens. Emission was filtered using a 655 LP filter and captured by a cooled EMCCD camera (Andor - iXon DU897) at 55.6 Hz. For 3D localizations, we applied a double phase ramp module (Zeiss) for phase ramp imaging localization microscopy (PRILM). 3D information was acquired from the z-position dependent rotational angle of the two lobes emanating from each fluorophore given the double phase ramp module ^56^. All 3D-SMLM experiments were conducted in highly inclined and laminated optical sheet (HILO) illumination mode. A total of 30000 frames were recorded for each video.

### 3D-dSTORM – Analysis

Localizations were fit using the spliner implementation of the 3D-DAOSTORM code package ^57^. Calibrations were made using 100 nm fluorescent TetraSpeck beads. The beads were attached to a coverslip by initial addition of 50 µL 1M MgCl_2_ followed by deposition of 0.9 µL beads diluted in 500 µL water. Clustering of the data was performed using a DBSCAN algorithm, identifying clusters within a 60 nm radius of each localization with at least 15 qualifying neighbors. Given the reduced axial resolution compared to lateral resolution, we set the weighting of axial position versus lateral to 0.5. Several parameters were extracted for each cluster, including max size in X,Y,Z-direction, radius of gyration and density of localizations.

### Transient transfection for BioID2-MS assay

A day prior to transfection, low passage (below 15) HEK293 cells were seeded in TC treated T-175 culture flasks (Sigma-Aldrich) at a density of 5x10^6^ cells/flask. Transfection was performed using Lipofectamin 2000 and Opti-MEM (ThermoFisher). The manufacturer´s protocol was followed with a 3:1 Lipofectamin to DNA ratio. A total of 20 µg/well was used and cells were incubated with transfection mix in Opti-MEM for 5 hours. Opti-MEM was replaced with biotin supplemented growth media (50:50 F10+DMEM (ThermoFisher)), and cells were used for experiments 24 hours after transfection.

### Proximity biotinylation pull-down assay

On the day of experiment biotin containing growth media was decanted and cells were washed twice with PBS. The second wash was performed with ice cold PBS (4°C) containing complete protease inhibitor cocktail tablets and phosphatase inhibitor cocktail 3 (Sigma-Aldrich). Cells were subsequently kept at 4°C for the rest of the purification steps. Cells were collected and spun down at 300xg for 5 minutes. Supernatant was decanted and the pellets were snap-frozen in liquid nitrogen. Pellets were resuspended in 1 mL of ice-cold RIPA-lysis buffer (150 mM NaCl, 0.1% sodium dodecyl sulfate (SDS), 0.2% NP-40 and protease inhibitor in 50 mM Tris-HCL at pH 7.5) and homogenized using a hand-held homogenizer gun. The homogenate was incubated for 2 hours with end-over-end rotation. The lysates were centrifuged at 13000xg for 15 minutes. The resulting supernatant was transferred to fresh tubes and incubated with 50 µL/sample prewashed (in RIPA buffer) magnetic streptavidin-coated Dynabeads slurry for 1 hour with end-over-end rotation. Liquid was removed from Dynabeads using a magnetic DynaMag rack. Subsequently, beads were washed using a regimen of 2x low- (100 mM KCL and 0.1% TX-100 in 20 mM Tris-HCL at pH7.5), 2x high- (500 mM KCL and 0.1% TX-100 in 20 mM Tris-HCL at pH 7.5), 2x low- and 1 moderate-salt buffer (150 mM KCL and 0.1% TX-100 in 20 mM Tris-HCL at pH 7.5) prior to a wash in PBS, transfer to fresh tubes and a final wash in PBS. On-bead digestion using trypsin and subsequent LC-MS/MS analysis was performed by the Proteomics Research Infrastructure (PRI) facility of the Faculty of Health and Medical Sciences at the University of Copenhagen.

### Analysis and data presentation

Raw LC-MS/MS data files were directly loaded as input in MaxQuant software (version 1.6.15.0)^58^. The Andromeda search analyzed the runs against the Human reference proteome UP000005640 (downloaded from UniProtKB database in March 2021) and a 1% FDR peptide and protein cutoff was used. We specified carbamidomethylation on cysteines as fixed modification and oxidation (M) and N-term acetylation were set as variable modifications. Trypsin was selected as proteolytic enzyme, allowing a maximum of 2 missed cleavage sites. Results were filtered for a minimal peptide length of seven amino acids. We enabled match-between runs with a matching time window of 0.7 min and alignment time window of 20 min. Results were filtered for a minimal peptide length of seven amino acids. The resulting protein library was loaded into Perseus (version 1.6.15.0) for label-free quantification (LFQ) analysis. Data was filtered to remove proteins only identified by site as well as potential contaminants and reverse hits. We selected for proteins that were identified in all three replicates in at least one group, log_2_-transformed LFQ intensities and imputed missing values based on quantile regression imputation of left-censored data (QRILC) method using the imputeQRILC function of imputeLCMD package written for R.

The resulting LFQ intensities were used to calculate differential abundance (Difference) of proteins compared to Tac0-BioID2 and profile plots of the resulting mean differences were used to highlight trends throughout groups. To control for endogenous biotinylation levels, we analysed ratios of Tac0-BioID2 normalized LFQ values of TachAH-BioID2 constructs versus Tac0-BioID2 normalized LFQ values obtained for untransfected cells.

$$Abundance relative to Tac={LFQ}_{TacAH}-{‾LFQ}_{Tac}$$

$$Normalized abundance relative to pcDNA= \frac{{LFQ}_{TacAH}}{{‾LFQ}_{Tac}}- \frac{{LFQ}_{pcDNA}}{{‾LFQ}_{Tac}}$$

### Immunoblotting

Protein lysates were mixed with 5xsodium dodecyl sulfate polyacrylamide gel electrophoresis (SDS-PAGE) loading buffer (50 mM DTT and 355 mM MBE), boiled at 95°C for 5 min before loaded into an Any-kD (Mini-ProTEAN® TGXTM, BIO-RAD) and run at 115V for 1 hour. The size-separated proteins were transferred to polyvinylidene difluoride membranes (BIO-RAD) for 10 min at 25 V on a trans-blot turbo transfer system (Bio-Rad) and blocked for 1 hour in 5% milk in wash buffer (PBS with 0.05% Tween-20). Membranes were incubated with primary antibodies (anti-FLAG M1 antibody) for 1 hour at 25°C or overnight at 4°C, washed 3 times for 15min and incubated with HRP-conjugated secondary anti-mouse antibodies for 30 min. Membranes were washed 3 times for 15 min before developing using either the SuperSignal ELISA Femto Substrate (Thermo Fisher Scientific) or ELC Prime Western Blotting system (Sigma-Aldrich) and captured with a cooled CCD camera.

### Gel-assisted GUV formation

The general procedure for Gel-assisted giant unilamellar vesicles (GUV) formation followed a previously published protocol ^59^. In brief, a 5% (w/w) solution of PVA (molecular weight above 146 kDa, MERCK) was prepared by stirring PVA in 20mM TRIS (pH 7.4) and 150 mM sucrose while heating to 90°C until appearing clear to the eye. We applied a thin film of 15µL PVA solution to a plasma-cleaned microscope coverslip (20 mm x 20 mm, Menzel-Gläser) and left the solution to dry for 30 minutes at 50°C. We spread 10 µL of lipids (1 mg/mL) on the dried PVA and evaporated the solvent for 60 minutes under vacuum. A lipid composition of [79.3 mol% DOPC:10 mol% DOPS:10 mol% cholesterol:0.5 mol% BODIPY-TR:0.3 mol% PEG-Biotin] dissolved in chloroform was used. Next, we applied 1 mL of growth buffer solution (5 mM MOPS (pH 7.4), 20 mM NaCl, 255 mM sucrose in water adjusted to 300 mOsm by addition of sucrose) and left GUVs swelling for 30-45 minutes at 20°C. After swelling period, GUVs were detached by gentle tipping against the bottom of the chamber and transferred to Eppendorf tubes using a P1000 pipette with a cut tip to decrease chance of bursting. The prepared GUV solution was used the earliest after a 15-minute equilibration period and kept at 4°C for a maximum of 3 days.

### Pulling nanotubes from GUVs

The general procedure for pulling nanotubes followed a previously published protocol ^60^. In brief, we prepared micropipettes by pulling borosilicate glass capillaries (GC100-15, Harvard Apparatus) with a Sutter P-2000 (Sutter instruments). The tip was refined using a microforge, resulting in a final inner diameter of 5-7 microns, corresponding to ~1/3 of typical GUV diameter investigated. Prior to experiments, both the experimental chamber and the micropipette for micromanipulation of GUVs were passivated using a 5 mg/mL solution of highly pure ß-casein (C6905, Sigma-Aldrich) in experimental buffer (20 mM MOPS (pH 7.4), 100 mM NaCl, 90 mM glucose in water adjusted to 310 mOsm by addition of glucose) and leaving the solution for more than 30 minutes to minimize the chance of GUVs adhering to the glass surface and bursting. After passivating, the experimental chamber was washed 3 times in experimental buffer and a few µL of streptavidin-coated polystyrene-beads (3-3.4 micron diameter, SVP-30-5, Spherotech) were added to the chamber. During the passivating step, we prepared a mix of 1 µM Oregon Green labelled peptide and 10% of the GUV solution prepared as described above in experimental buffer and let the mix incubate for 30 minutes prior to addition to the experimental chamber. We let the GUVs settle and deflate by leaving the buffer to evaporate for ~15 minutes, allowing them to be “floppy”, and then sealed the experimental chamber using mineral oil. Aspiration pressure of the micromanipulation pipette was then equilibrated using a polystyrene bead prior to every tube pulling recording. We then aspirated a GUV, trapped a bead using optical tweezers and positioned both at ~20 microns from the glass surface. Membrane tensions was relieved without losing the GUV from the micropipette and the GUV was carefully positioned using a piezo-actuator in proximity to the bead to allow for streptavidin-biotin bonds to form. Careful separation of GUV and beads then allowed nanotube formation and the aspiration pressure was increased to recreate an aspiration tongue. Movement of the bead was recorded using a bright field microscope for a minute, followed by confocal imaging, and recording of the water tank height position relative to equilibrium. This step was repeated multiple times at increasing aspiration pressure. Quantitative evaluation of membrane curvature sorting was then calculated as a sorting coefficient, S given by:

$$S=\frac{\frac{{Intensity}_{Protein in tube}}{{Intensity}_{Protein in GUV}}}{\frac{{Intensity}_{Lipid in tube}}{{Intensity}_{Lipid in GUV}}}$$

Where Intensity_Protein in tube_ and Intensity_Protein in GUV_ represent the fluorescence intensities of proteins and Intensity_Lipid in tube_ and Intensity_Lipid in GUV_ represent the fluorescence intensities of lipids on the tube and GUV respectively. These intensities were calculated given the difference in summed intensities observed over a range of line scans analyzed for regions of interest for GUV and tube (Figure 3H).

### Statistical Analysis

Analysis was performed using GraphPad Prism software v6.01. Statistical comparisons across groups were done with a one-way ANOVA in the case of confocal microscopy data. Averages from Flow Cytometry data were analyzed with one-way ANOVA using Dunnett’s correction for multiple comparisons, except for grouped data, as in the case of EE-mutants and Dyngo-treatment. Here a two-way ANOVA with Sidak or Tukey’s correction for multiple comparisons was applied. Linear regression was performed as standard, and goodness of fit was indicated with R-square and p values.

### Mathematical model “Acidification Kinetics Function”

Henderson-Hasselbach derived quenching dynamics

- Following the quenching-kinetics of pHluorin as a function of vesicular size

$$I_{SEP}=c_{SEP}*[SEP-]$$

$$pH=pKa+log(\frac{\left[ SEP- \right]}{\left[ SEPH \right]})$$

$$-\log\left( \left[ H+ \right] \right)= -\log\left( Ka \right)+\log\left( \frac{\left[ SEP- \right]}{\left[ SEPH \right]} \right)= log(\frac{\left[ SEP- \right]}{\left[ SEPH \right]*Ka})$$

$$\left[ SEP- \right]= \frac{Ka}{[H^{+}]}[SEPH]$$

$$\left[ SEPH \right]={[SEP]}_{tot}-[SEP-]$$

$$\left[ SEP- \right]*\left( 1+\frac{Ka}{\left[ H^{+} \right]} \right)= \left[ SEP \right]_{tot}*\frac{Ka}{[H^{+}]}$$

$$\left[ SEP- \right]={[SEP]}_{tot}*\frac{\frac{Ka}{[H+]}}{(1+\frac{Ka}{\left[ H^{+} \right]})}$$

Assumption 1: The density of V-ATPase in the membrane is not influenced by the construct:

$$\frac{d[H^{+}]}{dt} \propto\frac{4\pi r^{2}}{\frac{4}{3}\pi r^{3}}= \frac{1}{3r}$$

Since the relative radius is a constant that we will ultimately need to derive from the fit of each average trace:

$$\left[ H^{+} \right]\propto\frac{1}{3r}*t$$

Assumption 2: At timepoint t_0_, immediately upon scission, pH = 7.4, pKa = 6.9, i.e:

@t=0:

$${10}^{0.5}= \frac{\left[ SEP- \right]}{\left[ SEPH \right]}=\frac{\left[ SEP- \right]}{(\left[ SEP \right]_{tot}-\left[ SEP- \right])}$$

$$\left[ SEP- \right]_{t=0}={\frac{{10}^{0.5}}{{(1+10}^{0.5})}\left[ SEP \right]}_{tot}\approx0.76*{[SEP]}_{tot}$$

We want to calculate the relative amount of [SEP-] at time = t:

$$\frac{\left[ SEP- \right]_{t}}{{[SEP-]}_{t=0}}=f(t)$$

$$\frac{d[SEP-]}{dt}=\frac{{([SEP-]}_{t}-\left[ SEP- \right]_{t=0})}{(t-t_{0})}$$

To derive f(t) we first look at how [SEP-] changes with [H^+^], and thereby time t:

$$\frac{d\frac{[SEP-]}{{[SEP]}_{tot}}}{d[H^{+}]}=\left[ \frac{\frac{Ka}{[H+]}}{(1+\frac{Ka}{\left[ H^{+} \right]})} \right]^{'}$$

g(x) = $\frac{Ka}{[H^{+}]}$ ; g’(x) = -$\frac{Ka}{{[H^{+}]}^{2}}$ ; h(g) = $\frac{g}{(1+g)}$ ; h’(g) = $\frac{1}{{(1+g)}^{2}}$ ;

Chain rule: h(g)’ = h’(g)*g’ = $\frac{1}{\left( 1+\frac{Ka}{\left[ H+ \right]} \right)^{2}}$ * (- $\frac{Ka}{{[H^{+}]}^{2}}$) = $-\frac{\frac{Ka}{{[H^{+}]}^{2}}}{\left( 1+\frac{Ka}{\left[ H+ \right]} \right)^{2}}$

As expected, a decline of the relative amount of [SEP-] with increasing [H^+^]

Since we already deduced the relationship between radius, time and [H^+^]-increase above, we can then state the change of [SEP-]_rel_ with time t by substituting [H^+^] with $c_{t}\frac{t}{3r}$:

$$\frac{d{[SEP-]}_{rel}}{dt}=-\frac{c_{t}}{\left( \frac{t}{3r} \right)^{2}}*\frac{1}{\left( \frac{c_{t}}{\frac{t}{3r}}+1 \right)^{2}}=-\frac{9r^{2}c_{t}}{t^{2}}*\frac{1}{\left( \frac{3rc_{t}}{t}+1 \right)^{2}}=-\frac{9r^{2}c_{t}}{\left( 3rc_{t}+t \right)^{2}}$$

That means that as time passes, the relative concentration of [SEP-] will continuously decline, but with a progressively slower kinetic, as we also observe.

Since the radius is a time-independent constant in these measures, we can easily substitute and integrate:

$$c_{1}=9r^{2}c_{t}$$

$$c_{2}=3rc_{t}$$

$${[SEP-]}_{rel}= -c_{1}\int\frac{1}{t^{2}*\left( \frac{c_{2}}{t}+1 \right)^{2}}dt= -c_{1}\int\frac{1}{\left( c_{2}+t \right)^{2}}dt=\frac{c_{1}}{\left( c_{2}+t \right)^{2}}+c_{3}$$

Which allows us to fit the TfR5.5-data directly in our ppH-tracks:

$${[SEP-]}_{rel}= \frac{\left( 3r \right)^{2}c_{t}}{\left( 3rc_{t}+t \right)^{2}}+c_{ig}$$

NB: What we fit is the arbitrary intensity of SEP:

$$I_{SEP}=c_{quench}*\left[ SEP- \right]+c_{dist}*\left[ SEP- \right]$$

Since we assume that the contribution of the helical construct to the average speed with which it diffuses away from the membrane is negligeable compared to the effect on size-dependent quenching-kinetics, we can focus only on the first term.
